# Adipocyte JAK2 mediates aging-associated metabolic liver disease and progression to hepatocellular carcinoma

**DOI:** 10.1101/681809

**Authors:** Kevin C. Corbit, Camella G. Wilson, Dylan Lowe, Jennifer L. Tran, Nicholas B. Vera, Michelle Clasquin, Aras N. Mattis, Ethan J. Weiss

## Abstract

Non-alcoholic fatty liver disease (NAFLD) and steatohepatitis (NASH) are liver manifestations of the metabolic syndrome and can progress to hepatocellular carcinoma (HCC). Loss of Growth Hormone (GH) signaling is reported to predispose to NAFLD and NASH through direct actions on the liver. Here, we report that aged mice lacking hepatocyte *Jak2* (JAK2L), an obligate transducer of Growth Hormone (GH) signaling, spontaneously develop the full spectrum of phenotypes found in patients with metabolic liver disease, beginning with insulin resistance and lipodystrophy and manifesting as NAFLD, NASH and even HCC, independent of dietary intervention. Remarkably, insulin resistance, metabolic liver disease, and carcinogenesis are prevented in JAK2L mice via concomitant deletion of adipocyte *Jak2* (JAK2LA). Further, we demonstrate that GH increases hepatic lipid burden but does so indirectly via signaling through adipocyte JAK2. Collectively, these data establish adipocytes as the mediator of GH-induced metabolic liver disease and carcinogenesis. In addition, we report a new spontaneous model of NAFLD, NASH, and HCC that recapitulates the natural sequelae of human insulin resistance-associated disease progression. The work presented here suggests a attention be paid towards inhibition of adipocyte GH signaling as a therapeutic target of metabolic liver disease.

## INTRODUCTION

The global prevalence of NAFLD, a state of excess liver lipid accumulation in the absence of chronic alcohol intake, is estimated to be ∼24% (1). A subset of patients with NAFLD go on to develop non-alcoholic steatohepatitis (NASH) with associated inflammation and fibrosis. NASH is now the leading cause of liver transplantation in the United States and predisposes to HCC (2), the most common malignancy of liver worldwide (3). Currently, there are no FDA-approved treatments for NASH, which could be a reflection of the lack of faithful animal models that fully recapitulate its clinical presentation (4, 5).

NASH pathogenesis is complex and multi-factorial, though the risk factors are nearly identical to those predisposing to the metabolic syndrome (4). It was revealed nearly two decades ago that NAFLD is significantly associated with insulin resistance (IR) (6), and NAFLD-associated hepatic IR is independent of adiposity and glucose intolerance (7). While the association of IR with fatty liver is strong, the cause-and-effect relationship is unclear. Hepatic fat content results from a balance of fatty acid influx, de novo lipogenesis (DNL), secretion of triglycerides, and catabolism of fatty acids by beta-oxidation (8). Given that these four processes occur within the liver itself, the majority of research and treatment paradigms for NAFLD, NASH, and HCC have been focused on liver intrinsic mechanisms.

How IR promotes hepatic lipid accumulation is debated. Insulin suppresses gluconeogenesis and induces DNL in liver (9). Thus, the predicted consequence of IR is increased glucose production and reduced lipogenesis. However, it is well recognized that the former and not latter pathways are affected in diabetics, as high levels of gluconeogenesis co-exist with increased hepatic lipid burden in the setting of IR. This phenomenon has been termed ‘selective insulin resistance’ and been attributed to both liver-intrinsic and –extrinsic mechanisms (10, 11).

Prolonged fasting/starvation, such as in anorexia nervosa (AN), and lipodystrophy (LD) are other conditions are also associated with fatty liver (12, 13). Interestingly, these states of adipose tissue dysfunction have IR as a commonality and, especially in the case of LD, have biochemical features more closely aligned with prevalent forms of IR than those caused by defects in insulin signaling itself (13). This suggests that adipose tissue dysfunction may be a common feature, and possible a driver, of various form of IR.

As is the case with protracted fasting, NAFLD and NASH are associated with adipose tissue dysfunction (13, 14). GH is a starvation-induced hormone that controls hepatic (and circulating) IGF1 levels and adipose tissue lipolysis during fasting (15, 16) and is elevated in LD patients (17). Malnutrition, LD, AN, and NAFLD are associated with hepatic GH resistance, which may seem paradoxical, but in fact hepatic GH resistance leads to elevated circulating GH, via loss of IGF1-mediated feedback inhibition, which can act on non-liver, GH-responsive tissues. In fact, treatment of AN patients with GH indeed fails to increase insulin-like growth factor (IGF1) levels, confirming hepatic GH resistance, but does further decrease fat mass, indicating that adipose tissue remains GH responsive in the clinically GH ‘resistant’ state (18). Thus, elevated GH activity on adipose tissue may be a commonality among GH resistant and IR states, including starvation, LD, NAFLD/NASH.

GH is a major regulator of glucose and lipid metabolism (19, 20). Congenital loss of global GH signaling increases insulin sensitivity and adiposity and may decrease the incidence of diabetes and cancer (21, 22). Conversely, exposure to GH acutely (19) or chronically, in the setting of acromegaly, induces insulin resistance (23). Somewhat paradoxically, adult-onset of GH deficiency also predisposes to fatty liver and NASH (24), possibly by a different mechanism via direct effects on hepatic DNL (25). However, a unifying model of how GH impinges on insulin sensitivity to mediate glucose and lipid metabolism has not been universally accepted.

We previously reported that mice with hepatic GH resistance, via hepatocyte-specific deletion of *Jak2* (JAK2L), developed fatty liver in a GH-dependent manner and had early signs of NASH by 20-weeks of age (26). Here, we report that aged JAK2L mice are insulin resistant, lipodystrophic, and subsequently develop severe NASH and HCC. These phenotypes recapitulate the features of metabolic liver disease and are entirely dependent on adipocyte JAK2, as mice lacking both hepatocyte and adipocyte *Jak2* (JAK2LA) retain insulin sensitivity and maintain liver homeostasis. Treatment with recombinant GH increases liver tri-(TAG) and di-acylglycerol (DAG) levels in control but not in mice lacking adipocyte *Jak2* only. Collectively, we demonstrate that adipocytes are the target of GH-induced changes in liver metabolism. Further, we provide a new model of metabolic liver disease that is independent of dietary intervention.

## RESULTS

### Hepatic GH resistance promotes age-associated insulin resistance via adipocyte signaling

We aged cohorts of control (CON, N=16), JAK2L (N=14), and JAK2LA (N=17) mice to between 70-75 weeks of age and determined glucose homeostasis in the fed and fasted states. Similar to our previous results in younger mice (27), induction of hepatic GH resistance through hepatocyte-specific deletion of *Jak2* in JAK2L and JAK2LA mice essentially eliminated detectable circulating IGF1 (Figure 1A). This abolished IGF1-mediated negative feedback on central GH production and resulted in ∼200 times higher fasting serum GH levels in both JAK2L and JAK2LA animals compared to the CON cohort (Figure 1B). Blood glucose levels varied little between the three genotypes, with only JAK2LA mice having statistically lower levels of fed glucose compared to CON mice (Figure 1C). CON mice appropriately showed lower serum insulin levels following on overnight fast, however JAK2L animals had both fed and fasting hyperinsulinemia (Figure 1D). This led to a large increase in HOMA-IR in the JAK2L mice that was normalized in JAK2LA animals (Figure 1E). Insulin tolerance testing (ITT) revealed augmented responsiveness in JAK2LA mice as compared to CON and JAK2L cohorts (Figure 1F). While HOMA-IR and ITT results were not concordant in these cohorts, HOMA-IR is more closely correlated with hepatic than peripheral insulin sensitivity (28), consistent with our previous published work using hyperinsulinemic-euglycemic clamps in JAK2L mice (20). Therefore, aged mice lacking hepatocyte *Jak2* are GH resistant and develop insulin resistance in an adipocyte *Jak2*-dependent manner.

**Figure 1:**
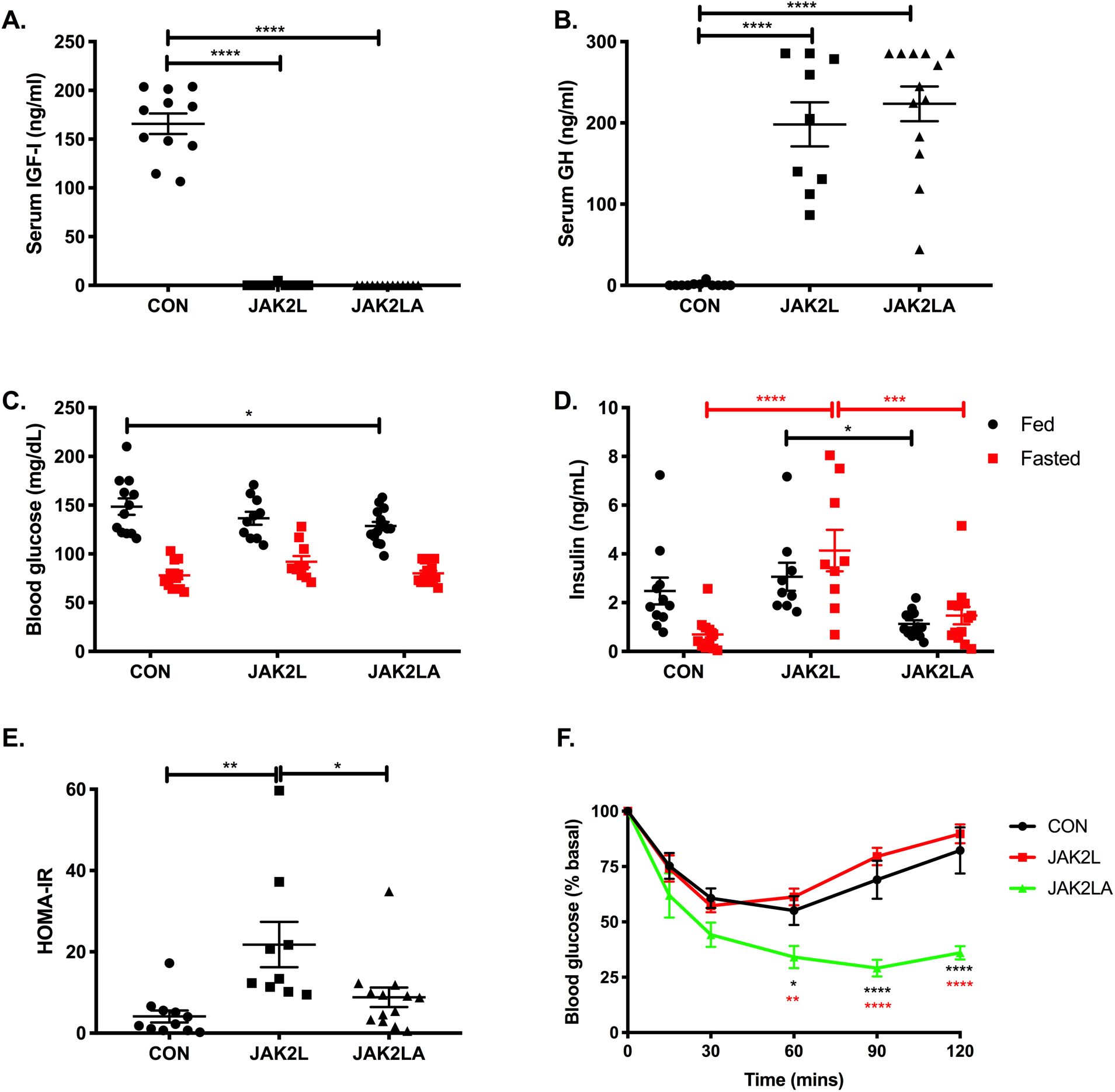
hepatic GH resistance does not correlate with insulin resistance in mice lacking adipocyte *Jak2*. Serum (A.) IGF-1 and (B.) GH levels in 16-hour fasted control (CON), JAK2L, and JAK2LA mice. (C.) Blood glucose and (D.) serum insulin levels in ad lib fed (black) and 16-hour fasted (red) mice. (E.) Homeostatic assessment model of insulin resistance (HOMA-IR) values. (F.) Insulin tolerance testing in control (CON, black), JAK2L (red), and JAK2LA (green) mice. N=9-13 (A, B, D, and E), 10-15 (C), and 6-8 (F). \**p*<0.05, \*\**p*<0.01, \*\*\**p*<0.001, \*\*\*\**p*<0.0001 by 1Way (A, B, and E) and 2Way ANOVA (C, D, and F).

### JAK2L mice are lipodystrophic and have defective adipose tissue signaling in response to feeding

Aged JAK2L mice weighed less than the CON and JAK2LA cohorts in both the fed and fasted states (Figure 2A). Interestingly, JAK2L mice lost more weight following an overnight fast, consistent with the role of GH as a catabolic ‘starvation’ hormone (Figure 2B). Dual-energy absorptiometry (DEXA) scanning revealed an increase in lean mass and loss of fat mass in JAK2L mice that was normalized in the JAK2LA cohort (Figure 2C). While relative visceral (epididymal pads) fat mass did not statistically differ between the groups (Figure 2D), a large reduction in subcutaneous (inguinal pads) fat was observed in JAK2L animals, while JAK2LA mice had increased relative subcutaneous fat mass (Figure 2E). Histological sectioning revealed smaller adipocytes and sclerotic tissue in JAK2L inguinal fat pads (Figure 2F). In contrast, JAK2LA fat pads were histologically devoid of fibrotic lesions and contained adipocytes of a size comparable to CON (Figure 2F). At the molecular level, acute re-feeding induced mTORC1 activity, a major regulator of the fasting-fed transition (29), in inguinal adipose tissue (Figure 2G). The adipose mTORC1 response to re-feeding was entirely abolished in JAK2L but not JAK2LA mice (Figure 2G). Collectively, high levels of circulating GH in JAK2L mice is associated with lipodystrophy and aberrant fasting-fed transitional adipose tissue signaling that is governed by adipocyte JAK2.

**Figure 2:**
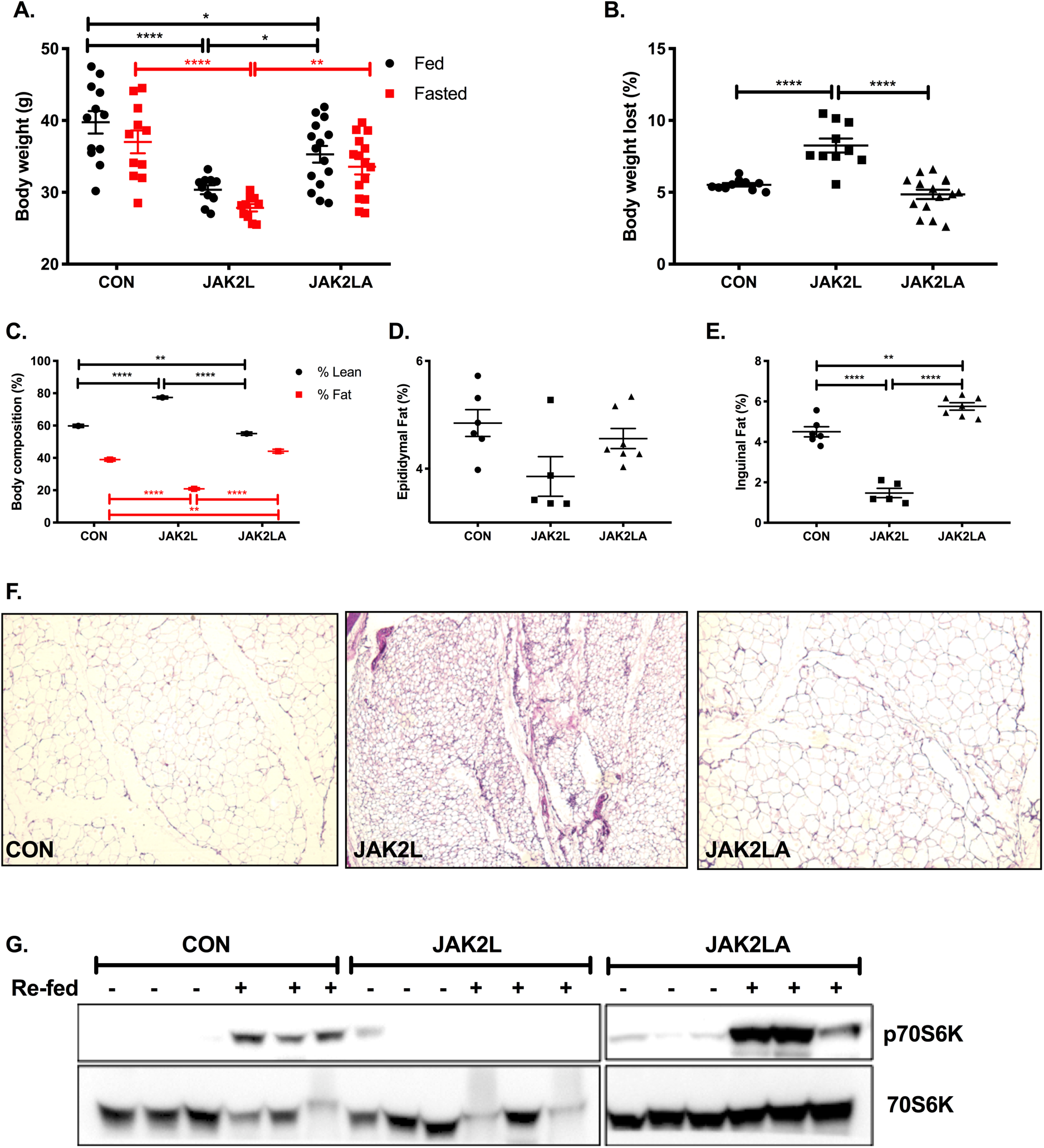
JAK2L mice are lipodystrophic and have a defective fasted-to-fed response in adipose tissue. (A.) Body weight in grams (g) of ad lib fed and 16-hour fasted control (CON), JAK2L, and JAK2LA mice. (B.) Percent body weight lost following a 16-hour fast. (C.) Percent lean and fast mass. Amount of (D.) epididymal and (E.) inguinal fat mass as a percent of total body weight. (F) H&E staining of inguinal fat pads from 16-hour fasted mice. (G) Inguinal adipose levels of phosphorylated (T389) p70S6K (top) and total p70S6K (bottom) in 16-hour fasted (-) or 30-minute re-fed (+) as determined by Western blot. N=10-15 (A and B), N=12-17 (C), N=5-7 (D and E). \**p*<0.05, \*\**p*<0.01, \*\*\*\**p*<0.0001 by 1Way (B and E) or 2Way (A and C) ANOVA.

### Loss of hepatocyte Jak2 promotes increased hepatic lipid burden and dyslipidemia in an adipocyte Jak2-dependent manner

Given the association of IR and lipodystrophy with hepatosteatosis, we examined the livers of aged CON, JAK2L, and JAK2LA mice. As a percent of total body weight, JAK2L animals had increased liver weight compared to both the CON and JAK2LA cohorts (Figure 3A). In contrast, JAK2LA mice had decreased liver weight (Figure 3A). Both total hepatic triglycerides (Figure 3B) and cholesterol (Figure 3C) were increased in JAK2L mice in an adipocyte *Jak2*–dependent manner. Markers of liver injury, including ALT (Figure 3D), AST (Figure 3E), and AP (Figure 3F) were increased in JAK2KL animals, and only AST levels were not entirely normalized in the JAK2LA cohort. Neither fasting serum NEFA (Figure 3G) nor triglycerides (Figure 3H) differed between the groups. However, fasting total cholesterol (Figure 3I) as well as HDL-C (Figure 3J) and LDL-C (Figure 3K) levels were increased in JAK2L mice. Again, these levels were corrected by concomitant deletion of adipocyte *Jak2*. Thus, hepatic GH resistance promotes increased liver lipid burden, liver injury, and dyslipidemia via effects on adipocytes.

**Figure 3:**
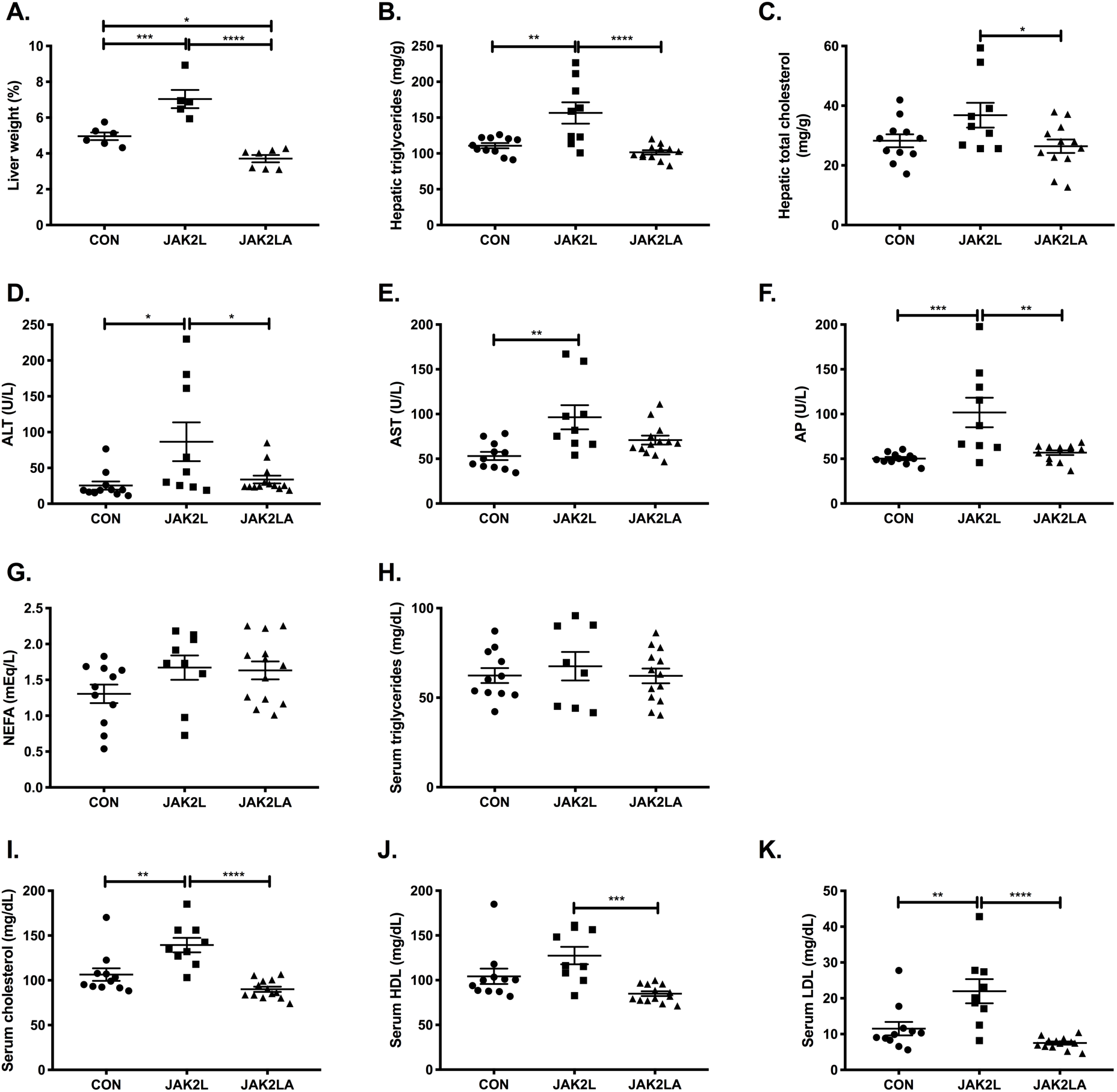
Loss of hepatocyte *Jak2* promotes liver damage and dyslipidemia in an adipocyte *Jak2*-dependent manner. (A) Liver weight as a percent of total body weight in 16-hour fasted control (CON), JAK2L, and JAK2LA mice. Hepatic (A.) triglycerides and (C.) total cholesterol levels in 16-hour fasted mice. (D). Alanine aminotransferase (ALT), (E.) Aspartate aminotransferase (AST), (F.) Alkaline phosphatase (AP), (G.) Non-esterified fatty acids (NEFA), (H.) Triglycerides, (I.) Cholesterol, (J.) High density lipoprotein (HDL), and (K.) Low density lipoprotein levels in 16-hour fasted serum. N=5-7 (A), N=9-12 (B and C), N=9-13 (D-K). \**p*<0.05, \*\**p*<0.01, \*\*\**p*<0.001, \*\*\*\**p*<0.0001 by 1Way ANOVA.

### JAK2L mice spontaneously develop age-associated NAFLD and NASH in an adipocyte Jak2-dependent manner

Livers collected from the aged cohorts were sent to a pathologist for a blinded histological assessment. H&E stained liver sections of aged CON mice demonstrated moderate lipid accumulation, which was slightly increased by hepatocyte-specific deletion of *Jak2* (Figure 4A, B and C). In contrast, few lipid droplets were observed in mice with loss of both hepatocyte and adipocyte *Jak2*. Trichrome staining revealed some collagen deposition in the CON cohort, while fibrosis was more widespread in liver sections from JAK2L animals (Figure 4B). Similar to lipid droplets, Trichrome staining was reduced in JAK2LA mice. Upon quantification, ∼54% of CON hepatocytes had lipid droplets, as compared to 74% and 9% for the JAK2L and JAK2LA cohorts, respectively (Figure 4A and C). For CON and JAK2L livers, steatosis was primarily localized to zone 3, whereas in JAK2LA livers, lipid-laden hepatocytes were also found in the azonal and panacinar regions. Ballooning, an indication of cellular stress, was found to be rare in CON liver sections. In contrast, histological ballooning was much more common in centrizonal sections of JAK2L mice. The severity of ballooning was significantly reduced, but not entirely corrected, by concomitant deletion of adipocyte *Jak2* (Figure 4D). Inflammatory lymphocytic foci were present in ∼2/3 of sections examined from CON livers, while foci were identified in all JAK2L sections (Figure 4E). The number of inflammatory foci were reduced in JAK2LA sections as compared to JAK2L but were not completely normalized to CON levels. Scoring for fibrosis (Figure 4F) and Brunt (Figure 4G) staging resulted in highly significant increases for the JAK2L, but not JAK2LA, cohort as compared to CON mice. Thus, JAK2L mice develop NASH with aging with retained JAK2 functioning in adipocytes.

**Figure 4:**
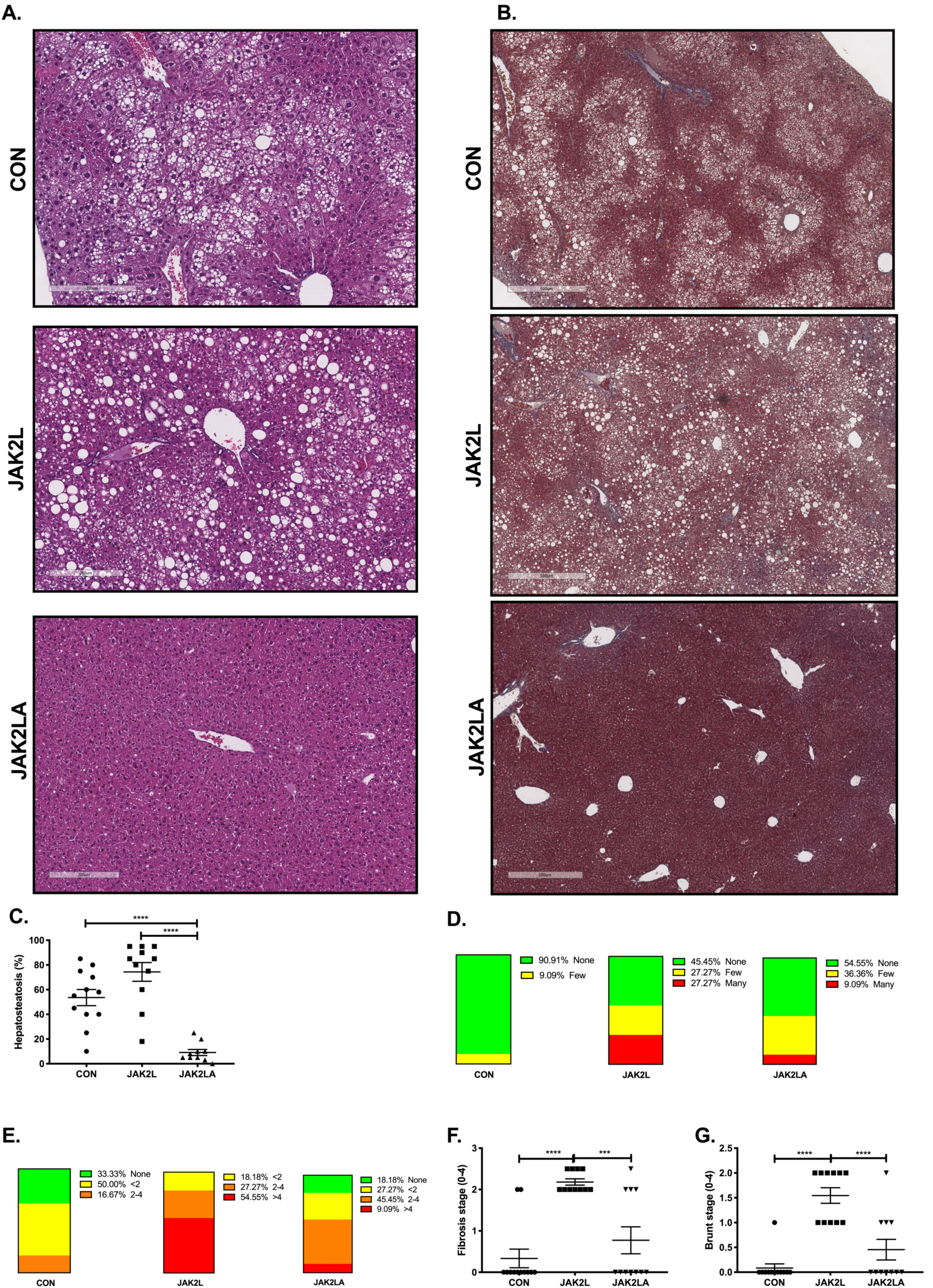
Loss of hepatocyte *Jak2* promotes NAFLD and NASH in an adipocyte *Jak2*-dependent manner. (A.) H&E and (B.) Trichrome staining of liver sections from control (CON), JAK2L, and JAK2LA mice. (C.) Percent hepatosteatosis, (D.) Ballooning, and (E.) Inflammatory loci observed in liver sections. (F.) Fibrosis staging score. (G.) Brunt staging score. N=10-12. \*\*\**p*<0.001, \*\*\*\**p*<0.0001 by 1Way ANOVA.

### JAK2L mice spontaneously develop HCC

Given that a percentage of NASH patients develop HCC, we allowed cohorts of CON (N=17), JAK2L (N=24), and JAK2LA (N=20) mice to age until natural death (here, defined as veterinarian-mandated euthanasia). Upon necropsy, livers from both CON and JAK2L were pale and large, while JAK2LA mice had smaller, red-colored livers (Figure 5A). We noticed that a number of livers from JAK2L mice had large growths (Figure 5B). Immuno-histochemical assessment of the tumors found them to be fibrotic (Figure 5C) and positive from both Glutamine Synthetase and Glypican-3 (Figure 5D), indicative of HCC (30). Tumor sections were negative for CK19 (Figure 5D), favoring a diagnosis of HCC over cholangiocarcinoma (31). Quantification of HCC incidence in our cohort (Figure 5E) revealed a statistical increase in JAK2L mice (Chi-square test, *p*=0.0077), with an average age of discovery of ∼704 days. In contrast, HCC incidence in the JAK2LA cohort did not differ from control mice. In summary, mice with high levels of circulating GH due to hepatocyte-specific deletion of *Jak2* spontaneously develop HCC with age. In contrast, high levels of GH in the setting of dual hepatocyte and adipocyte *Jak2* deficiency appear protected against age-associated HCC.

**Figure 5:**
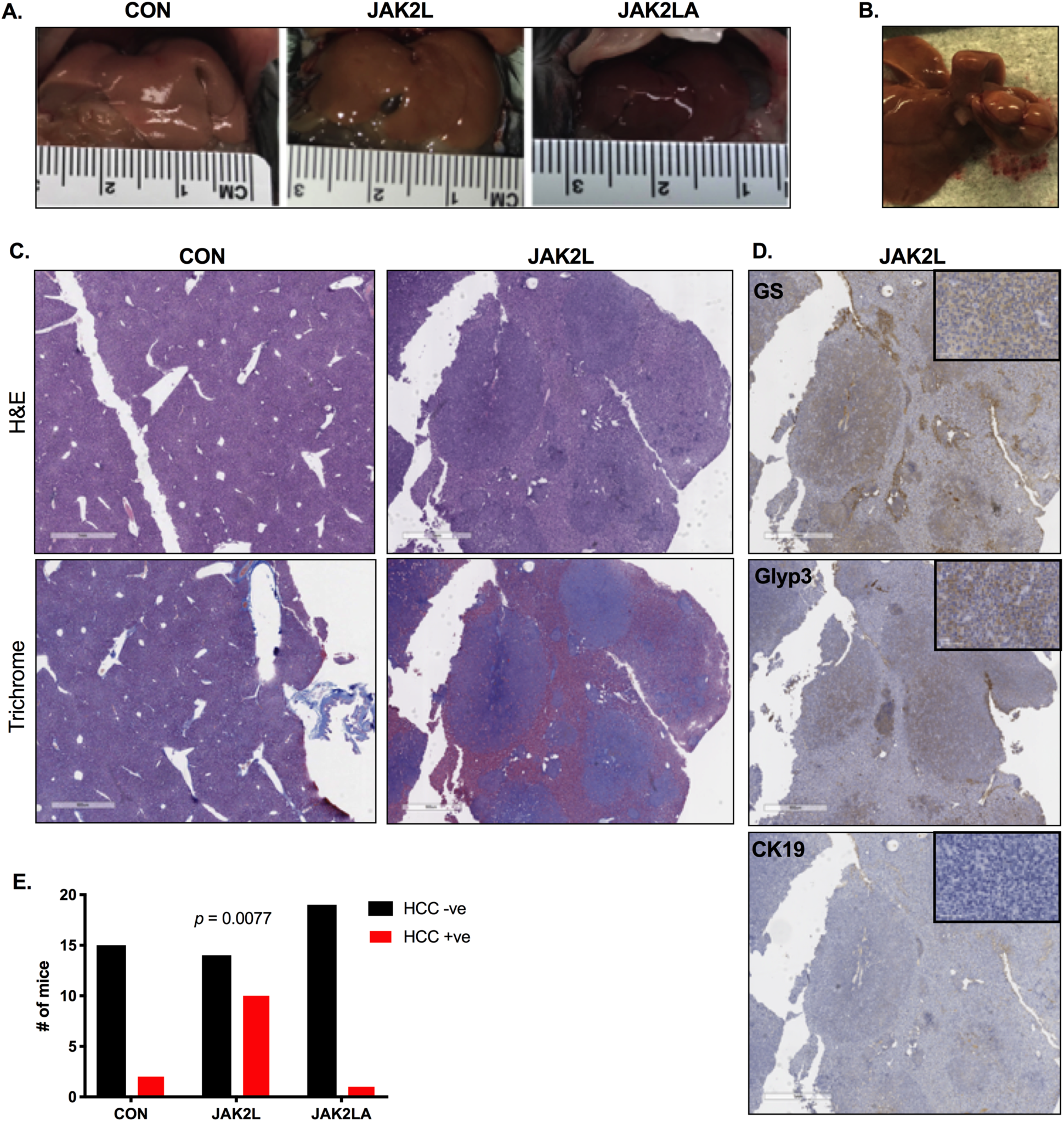
loss of hepatocyte *Jak2* promotes HCC in an adipocyte *Jak2*-dependent manner. (A.) Pictures of gross livers from control (CON), JAK2L, and JAK2LA mice. (B.) Liver nodules on a JAK2L liver. (C.) H&E and Trichrome staining of liver sections from CON mice and JAK2L nodules. (D.) Immunohistochemistry (high magnification shown in inset) of sections from JAK2L liver nodules stained with anti-Glutamine Synthetase (GlutSyn), anti-Glypican3 (Glyp3), and CK19. (E.) Contingency evaluation of HCC incidence showing the number of mice with HCC negative (HCC –ve) and positive (HCC +ve) tumors, N=17-24. *P* value determined by Chi-square testing.

### GH increases hepatic lipid burden through adipocyte Jak2

JAK2 is known to transduce signals from cytokines other than GH. Therefore, the phenotypic changes in hepatic metabolism observed in JAK2L and JAK2LA mice may not be entirely due to loss of GH signaling. To more definitely address the role of GH, we injected control (CON) and mice lacking adipocyte JAK2 (JAK2A) with vehicle or GH daily for seven consecutive days. We chose JAK2A over JAK2LA mice in order to specifically address the role of adipocyte JAK2, without any confounding effects of combinatorial hepatocyte deletion, in mediating metabolic changes following GH exposure. Livers were collected and subjected to an LCMS panel targeting ∼200 lipids. Heat maps derived from the top 99% of lipid species detected show that GH treatment grossly augmented hepatic TAG (Figure 6A) and DAG (Figure 6B; for raw lipidomics data and concentration waterfall plots see Supplemental Table 1 and Supplemental Figure 1, respectively) species in CON mice. One week of GH treatment increased total hepatic TAG (Figure 6C) and DAG (Figure 6D) by an average of ∼18% and ∼15%, respectively, in CON but not JAK2A mice. Hepatic cholesterol ester (CE) species were less affected by GH treatment (Figure 6E). However, GH treatment in CON mice did produce statistically higher levels of total hepatic CE than JAK2A animals (Figure 6G). GH treatment did not differentially affect total hepatic ceramide (CER) when comparing CON and JAK2A cohorts (Figures 6F and 6H). Therefore, acute GH treatment increases hepatic lipid burden but does so via adipocyte signaling.

**Figure 6:**
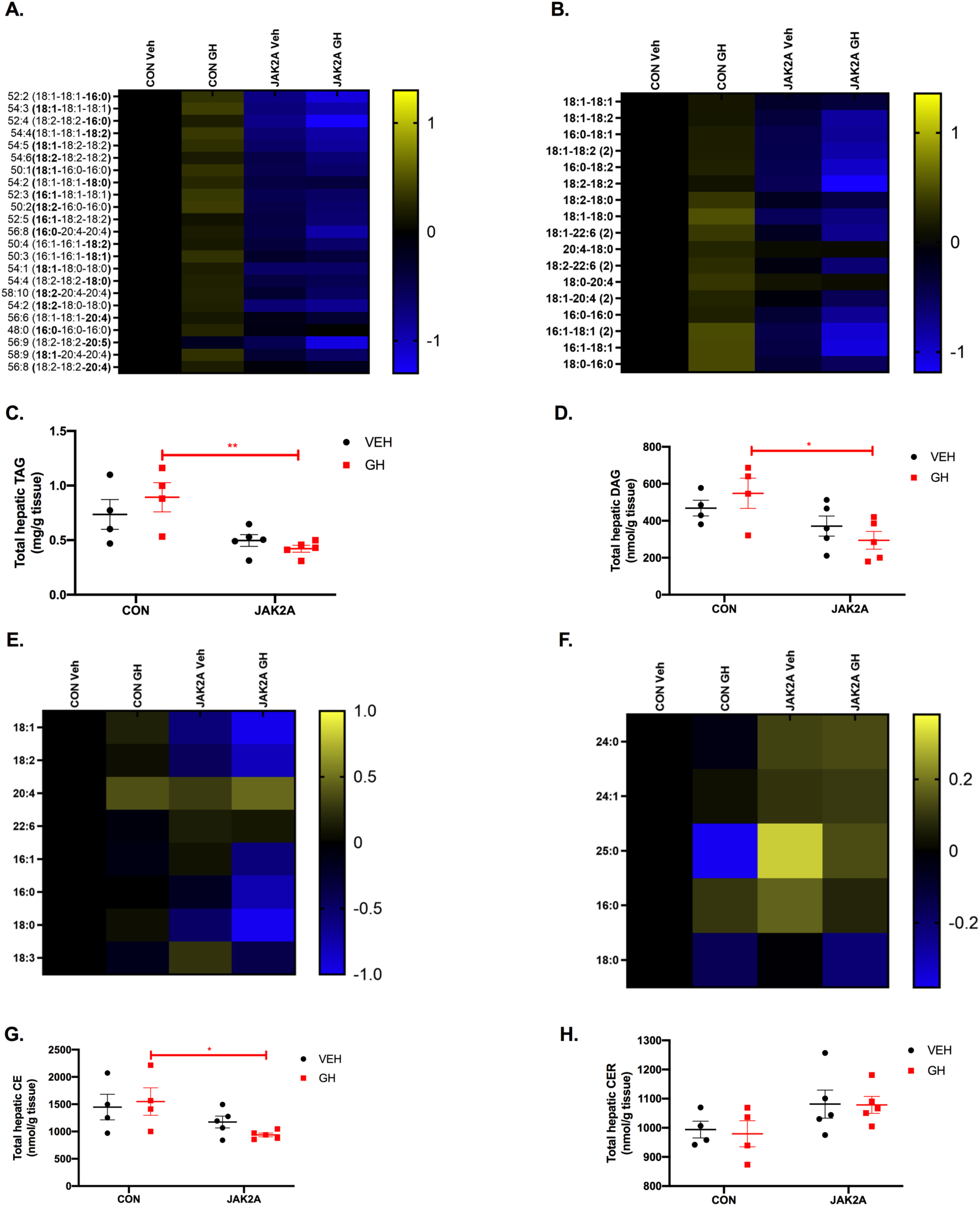
Growth hormone treatment induces hepatic lipid deposition via adipocyte JAK2. Lipidomics heat maps of individual hepatic (A.) Triacylglycerol (TAG), (B.) Diacylglycerol (DAG), (E.) Cholesterol ester (CE), and (F.) Ceramide (CER) species in vehicle (Veh)- or Growth hormone (GH)-treated control (CON) and JAK2A mice. Total hepatic (C.) TAG, (D.) DAG, (G.) CE, and (H.) CER levels. \**p*<0.05. \*\**p*<0.01 by 2Way ANOVA. For lipidomic heat maps, all values of individual lipid species are expressed as the log2 ratio to vehicle-treated CON mice and visualized by the log2 scale to the right of the heat map. N=4-5.

## DISCUSSION

The association between IR and type 2 diabetes (T2D) with hepatosteatosis is strong (32). However, it presents a number of physiological quandaries. Chief among them is why fatty liver develops in the setting of IR (33). The normal physiological roles of insulin action on the liver are to suppress hepatic glucose production (HGP) and to stimulate TAG synthesis. Thus, the prediction would be that patients with T2D would have hyperglycemia (from the inability of insulin to suppress HGP) concomitant with “normal” liver lipid levels (from the inability of insulin to induce TAG synthesis). Nevertheless, patients with T2D, while indeed exhibiting increased gluconeogenesis, also display increased rates of TAG synthesis, suggesting the latter pathway remains active while the former loses insulin responsiveness. This paradox has been termed “selective hepatic insulin resistance” and has been primarily attributed to branch points (i.e. one arm regulating HGP and a second mediating TAG synthesis) of insulin signaling within the liver itself (10, 11). TAG synthesis via hepatic DNL has been proposed as the potential driver of fatty liver disease (34) and a target of hepatic GH action (25). The natural conclusion would be to treat T2D and metabolic liver disease by targeting liver intrinsic mechanisms.

An alternative hypothesis is that substrate delivery can drive HGP and TAG synthesis independent of direct effects of insulin on the liver (35). From this vantage, diseases associated with hepatic IR could be treated by either shutting off the supply of substrate or inhibiting substrate uptake. In support of this, a study on NAFLD patients demonstrated that the majority of hepatic TAG arises from NEFA, more than twice of that derived from DNL (36). Furthermore, it was reported in high fat diet-fed rats that NEFA but not hepatic TAG accumulation is the cause of liver cell injury (37). Regardless, registered interventional NASH trials almost exclusively examine the effects of agents targeting mechanisms within the liver. Therefore, the effects of abrogating substrate supply or uptake remain unexamined, although preclinical models are supportive of pursuing this paradigm (37, 38). Here, we show that inhibition of adipocyte JAK2 signaling prevents the sequelae of events associated with high levels of circulating GH, beginning with insulin resistance and progressing through HCC. Specifically, the potential, if any, of inhibiting adipocyte GH signaling for the treatment of established NAFLD, NASH, and HCC are unknown.

Short term starvation is a natural state of IR and fatty liver (39). During nutrient deprivation, GH induces IR and promotes a lipolytic response (15). It has been hypothesized that GH resistance, which results from high circulating GH levels, is an adaptive response to states of under-nutrition to maintain euglycemia (40). Adipocyte (41) but not hepatocyte (42) IR results in LD and can promote fatty liver. This suggests that starvation and LD-induced fatty liver may share a common mechanism involving GH-mediated adipocyte IR. Interestingly, while AN is characterized by GH resistance (i.e. elevated circulating GH in the face of low plasma IGF1), treatment of AN patients with GH further induces fat loss. This suggests that AN is actually a state of *hepatic* GH resistance while adipose tissue of AN patients remains GH-responsive. Consistent with this, we now show that mice with elevated GH (JAK2L) lose more body weight following a 16-hour fast and are lipodystrophic. We interpret this as an unrestrained chronic state of adipose tissue insulin resistance and a condition mimicking metabolic starvation. At the molecular level, adipose tissue of JAK2L mice fail to induce mTORC1 following acute re-feeding, suggesting that GH impinges on mTORC1 activity. Interestingly, loss of mTORC1 specifically in adipocytes induces IR, fatty liver disease, and LD (43), a phenocopy of the JAK2L mice. These phenotypes associated with loss of hepatocyte *Jak2* -IR, adipose tissue mTORC1 activation, LD, and fatty liver - are rescued in JAK2LA mice. Therefore, adipocytes govern GH-mediated catabolism and fat mobilization and subsequent perturbation of liver metabolism.

Adult GH deficiency (AGHD) commonly results in the development of NAFLD and NASH (24). The association of AGHD with fatty liver has been attributed to the ability of GH to inhibit hepatic de novo lipogenesis (DNL); hence, in the absence of liver GH action, DNL would be increased (25). This is in contrast to our previous work using *in vivo* ^2^H2O labeling, as we found no increase in DNL in mice with fatty liver lacking hepatic GH signaling (26). Instead, using precise tissue-specific genetic models, we have determined that fatty acid uptake, via Cd36, is responsible for liver intrinsic mechanisms driving fatty liver in mice with disrupted hepatic GH signaling (38). Further, hepatosteatosis following hepatic GH resistance is dependent upon circulating GH, and since GH cannot transduce hepatic signals in the setting of hepatic GH resistance, a tissue other than liver must mediate fatty liver development (26). Here, our lipidomics data demonstrate that one-week of GH treatment promotes lipid deposition in livers of mice with intact hepatic GH signaling but not in mice lacking adipocyte GH signaling. Thus, both loss of hepatic GH sensitivity as well as continued hepatic exposure to GH in GH-sensitive animals promotes fatty liver. This may seem contradictory but that hepatic GH resistance results in high levels of circulating GH, similar to peripheral administration of recombinant GH. As we show here, hepatosteatosis induced by either GH resistance or GH treatment is governed by adipocyte JAK2. Regardless, the association of AGHD with liver fat and dysfunction has led to the dominant paradigm of augmenting GH levels to treat NAFLD and NASH. We propose that this approach will likely worsen the pathophysiology of established NAFLD and NASH, as was reported for patients with anorexia nervosa (18) and HIV-associated LD (44, 45).

Cordoba-Chacon *et al.* recently reported that knockdown of hepatocyte *Ghr* in adult mice (aHepGHRkd) promotes fatty liver and NASH via increased DNL without severe alterations in systemic metabolism or adipose tissue lipolysis (46). This led them to conclude that liver disease from loss of hepatocyte GH signaling occurs via liver autonomous means. However, it is difficult to make these conclusions in the face of high circulating GH, as reported in their aHepGHRkd mice, which can act on non-liver tissue, as well as insulin resistance (hyperinsulinemia and hyperglycemia). Previously, we controlled for augmented circulating GH in mice lacking hepatocyte GH signaling by global disruption of GH secretion (26). These studies demonstrated that circulating GH mediated onset of fatty liver in JAK2L mice, demonstrating that cells other than hepatocytes were responsible for development of hepatosteatosis. aHepGHRkd did not display increased plasma NEFA, which is consistent with our study here, as we did not observe a correlation between circulating fasting NEFA levels and liver pathology. However, given that circulating NEFA levels are the net result of release and uptake, and that loss of hepatic GH signaling induces *Cd36* (which increases NEFA uptake), it is difficult to make any conclusions on the state of lipolysis. To directly address this, Cordoba-Chacon *et al.* used adipose explant cultures to demonstrate that aHepGHRkd had normal basal and stimulated rate of lipolysis. This is also consistent with our previously published results showing that GH does not affect basal lipolysis but instead specifically interferes with the ability of insulin to suppress lipolysis (19). Thus, without directly attending to high circulating GH levels present in aHepGHRkd mice via concomitant loss of adipocyte *Ghr* or global *Gh* disruption it is premature to conclude that fatty liver and NASH resulting from loss of hepatocyte *Ghr* occurs via liver autonomous means.

The role of JAK2/STAT5 signaling in HCC is well documented. Liver-specific *Stat5* knockouts (STAT5L), like our JAK2L mice here, succumb to HCC upon aging without carcinogen or dietary intervention (47). In a study by Yu *et al.,* STAT5L mice developed HCC at age 17 months, although it was a small cohort of four mice. In addition, a previous study examined liver tumorigenesis in JAK2L mice and reported a 68% incidence at 60 weeks of age, although it is unclear if the tumors were HCC (48). Here, we report that 10/24 (∼42%) of JAK2L mice develop HCC with an average age of discovery of 704 days. Contrary to these findings, one report found that loss of hepatocyte JAK2 *protects* against liver tumorigenesis (49). However, the work by Shi *et al.* used chemical and dietary perturbations to induce liver disease and carcinogenesis. We propose that the reports, including our own, on aging-related HCC are mediated by adipocyte GH signaling and governed by IR while orally-administered carcinogens and dietary models directly affect hepatocytes. Our work here demonstrates that JAK2L mice succumb specifically to the sequalae of events leading to metabolic liver disease and, while it is experimentally cumbersome to wait ∼2 years for HCC development, may provide a more faithful model of human IR-associated HCC.

Acromegaly predisposes to cancer, and cancer has become a leading cause of acromegaly-associated deaths, despite an overall increase in life expectancy due to new treatments (50). The most parsimonious rationale for the mechanism of cancer onset due to high levels of GH is reduced plasma IGF1. However, we report here that mice with hepatic GH resistance indeed succumb to HCC, despite undetectable circulating IGF1. Further, while treatment of Laron Syndrome (LS) patients (loss of GH signaling) with recombinant IGF1 does normalize linear growth rates, restoration of IGF1 levels by pharmacological means does not prevent the immunity from cancer appreciated in those with LS (51). Collectively then, we propose that GH predisposes to cancer via its role in metabolism and not IGF1 production, possibly through actions on adipose tissue. Adipose tissue dysfunction, and IR in particular, is highly associated with cancer and adipocytes have been proposed as mediators of tumor cell behavior (52). How insulin resistant adipocytes promote oncogenesis is unknown, but a role for fatty acid receptors has been proposed (53). In addition, metabolic re-programming in HCC is mediated by the master oncogene *Myc* (54) and MYC levels themselves are induced by cellular exposure to fatty acids (55). Interestingly, a single bolus of GH increased hepatic *Myc* expression within one hour of administration (56). Therefore, it is tempting to speculate that adipocytes release a factor in a JAK2-dependent manner that subsequently induces hepatic *Myc*, leading to a favorable metabolic environment for development of HCC. Given that HCCs are ‘addicted’ to sustained MYC expression (57), it will be interesting to test if inhibition of adipocyte JAK2 will lead to regression of established HCC, what would ultimately be akin to a kind of ‘metabolite addiction’.

## METHODS

### Study Approval

All procedures performed on animals were in accordance with regulations and established guidelines and were reviewed and approved by the Institutional Animal Care and Use Committee at the University of California, San Francisco.

### Animals and diets

The generation of JAK2L and JAK2LA mice employing *Albumin:CRE* and *Adiponectin:CRE* was previously described (20). For our studies reported here, *Jak2^lox/lox^* (58) were used as controls and backcrossed onto the C57BL/6 background for at least nine generations. All mice in these studies were fed PicoLab Mouse Diet 20 (Lab Diet #5058; percent calories provided by protein 23%, fat 22%, carbohydrate 55%) throughout their lifetime.

### Study designs

Mice were aged until 70-75 weeks for metabolic studies. Blood was collected by retro-orbital puncture at 17:00 prior to food removal as the fed state. The following morning at 09:00 mice were sacrificed and blood and tissues collected as the fasted state. For re-feeding studies, mice fasted overnight were sacrificed or re-fed for 30 minutes prior to tissue collection by placing a pellet of food into the cages. Tissues were flash frozen on liquid nitrogen for Western blot, lipidomic, and transcriptomic analyses. Tissues were fixed with 10% neutral-buffered formalin for 24 hours for histological analyses. Blood glucose and serum insulin levels were determined by glucometer readings (Bayer Contour) and ELISA (Alpco), respectively. Serum IGF-1 (R&D MG100) and GH (Millipore EZRMGH-45K) levels were determined by ELISA. For insulin tolerance testing, mice were fasted for four hours (09:00-13:00) followed by intraperitoneal injection of 2 U/kg insulin (Novolin® Novo Nordisk, Bagsvaerd Denmark). Blood glucose levels were determined by tail prick using a hand-held glucometer at the times indicated. HOMA-IR was calculated as a ratio of fasting blood glucose (mg/dL):fasting serum insulin (mU/L) divided by 405 (59). Total fat mass was determined by Dual-energy X-ray absorptiometry. All clinical chemistry was done on terminally-collected serum from 16-hour fasted mice and done at the University of California, Davis Comparative Pathology Lab Core. For acute GH studies, CON and JAK2A mice were injected (i.p.) with vehicle (0.03 M NaHCO3, 0.15 M NaCl, pH 9.5) or 5 mg/kg recombinant mouse GH (Dr. A. F. Parlow, National Hormone and Peptide Program, UCLA, Torrance, CA USA) daily at 09:00 for 7 days and sacrificed 4 hours after a fast and the final injection.

### Hepatic TAG and cholesterol measurements and Western blots

Bits of frozen inguinal adipose pads were weighed and re-suspended in 1X cell lysis buffer (Cell Signaling Technology 9803) with protease (Roche 4693132001) and phosphatase (Roche 4906845001) inhibitor cocktail at a final volume of 50mg/ml and homogenized using the OmniTH homogenizer. 20 ul of lysates were used to determine cholesterol (Wako 999-02601) and triglyceride (Infinity reagent, Thermo TR22421) levels. Alternatively, lysates were ran out on 4-12% gradient gels and probed with anti-phospho(T389) p70S6K (Cell Signaling 9205) and anti-p70S6K (Cell Signaling 2708) antibodies. Western blots were developed with SuperSignal West Femto reagent (Thermo 34095) and developed using a ChemiDoc Imaging System (Bio-Rad).

### Liver histology and pathological analyses

Tissues were collected and fixed in 10% neutral-buffered formalin for 24 hours. Subsequently, tissues were washed and stored in 70% ethanol until embedding and sectioning. Sectioning and staining were carried out by the UCSF Liver Center core. H&E, Trichrome, reticulin, and immunohistochemical stains were reviewed and evaluated by a blinded GI/Liver pathologist. Livers were evaluated and scored according to methods by Kleiner and Brunt (60). Liver tumors were evaluated by H&E, Trichrome, and reticulin patterns and further evaluated by immunohistochemical staining for CK19, Glutamine Synthetase, and Glypican 3. Immunohistochemical staining was performed by the UCSF Liver Center core.

### Lipidomics

Lipidomics were done exactly as described (20).

### Statisitcs and graphics

All statistical tests and figures were done using GraphPad Prism v8.0.

## Acknowledgements

This study was supported by National Institutes of Health (NIH) Grants 1R01DK091276 (to E.J.W.) and DK076169 (E.J.W.), We also gratefully acknowledge the support of the James Peter Read Foundation, the University of California, San Francisco (UCSF) Cardiovascular Research Institute, the UCSF Diabetes Center (P30 DK063720), and the UCSF Liver Center (P30 DK026743). We would like to thank Dr. Kay-Uwe Wagner from the University of Nebraska for kindly providing the *Jak2* conditional mice.

## Author contributions

J. L. T. and K. C. C. carried out experiments for Figures 1–5. J. L. T. and C. G. W. did the ITTs for Figure 1. J. L. T. and D. L. performed experiments for Figure 6. N. M. did all liver pathological analyses. N. B. V. and M. C. performed lipidomics, made heat map and waterfall plots, and provided Supplemental Table K. C. C. carried out all ELISAs and Western blots. K. C. C. and E. J. W. conceived of the experiments, made the figures, performed statistical analyses, and wrote the paper.

## Notes

**Conflict of interest statement** N. B. V. and M. C. are employees and shareholders of Pfizer, Inc.

