## Supplemental files for "Adipocyte JAK2 mediates aging-associated metabolic liver disease and progression to hepatocellular carcinoma"

### Supplemental Figure 1

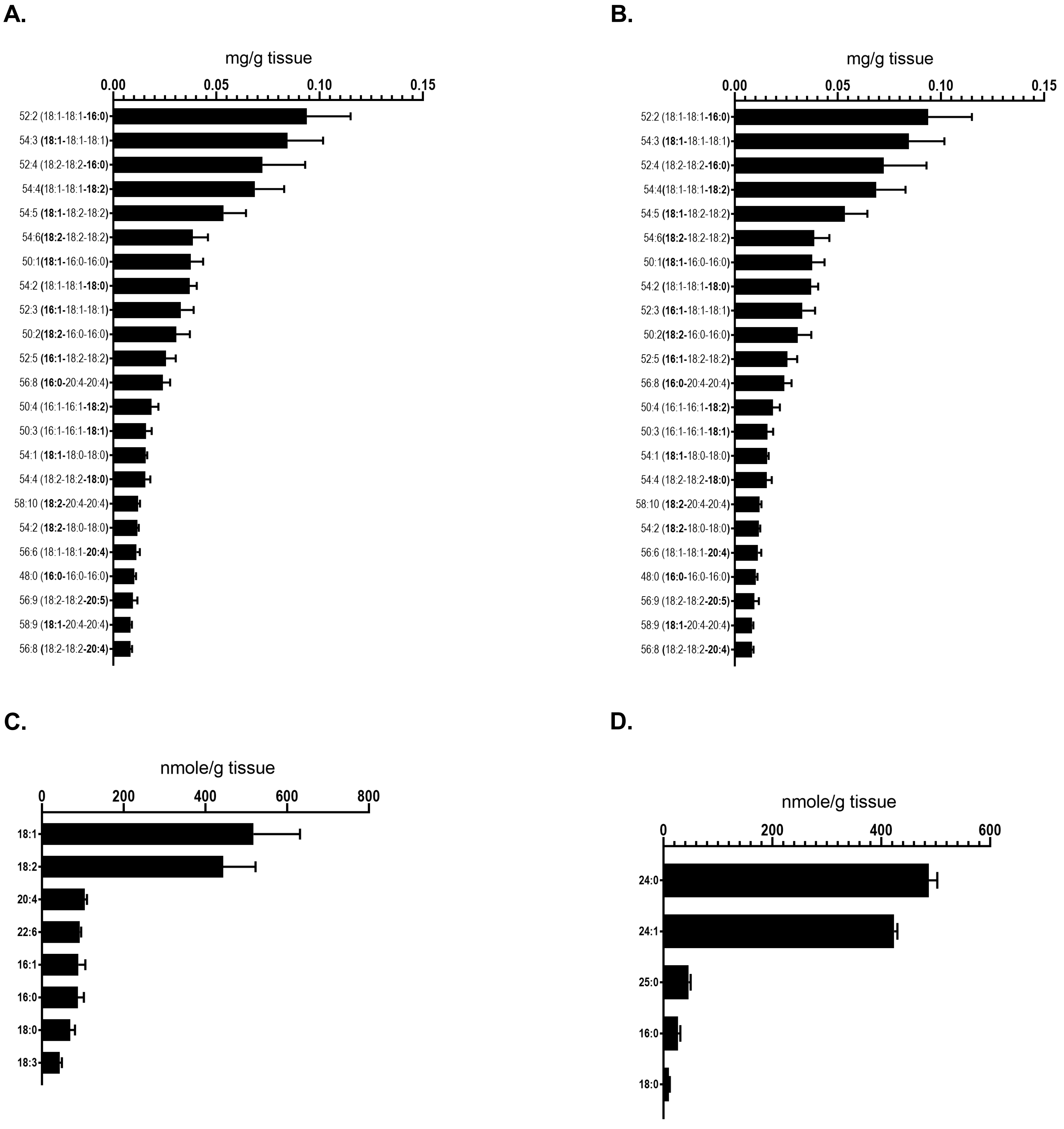

**Supplemental Figure 1: concentrations of individual lipid species shown in lipidomics heat maps.** Waterfall plots of the 99% most abundant (A.) Triacylglycerol, (B.) Diacylglycerol, (C.) Cholesterol ester, and (D.) Ceramide species.

| Componen | Total.Carbo | Lipid.Class | 2 (Control - | 6 (Control - | 13 (Control | 16 (Control | 21 (Control | 24 (Control |
|---|---|---|---|---|---|---|---|---|
| | | | CON-Veh | CON-Veh | CON-Veh | CON-Veh | CON-Veh-I | CON-Veh-I |
| CER 16:0 | 16:0 | Ceramide | 21.76873 | 38.55723 | 17.87069 | 27.36116 | 93.86485 | 17.62738 |
| CER 18:0 | 18:0 | Ceramide | 8.032578 | 13.74015 | 7.290517 | 11.46742 | 13.93507 | 7.139164 |
| CER 24:0 | 24:0 | Ceramide | 463.4499 | 524.4314 | 460.0862 | 502.0064 | 514.1623 | 513.7643 |
| CER 24:1 | 24:1 | Ceramide | 427.4914 | 438.7045 | 413.9224 | 415.618 | 498.1847 | 408.8213 |
| CER 25:0 | 25:0 | Ceramide | 37.38302 | 54.31137 | 42.64655 | 49.94061 | 40.51979 | 41.76426 |
| CER 18:1 | | Ceramide | 0.770342 | 1.059198 | 0.566897 | 0.879197 | 2.457097 | 0.768821 |
| Total | | Ceramide | 958.8959 | 1070.804 | 942.3833 | 1007.273 | 1163.124 | 989.8852 |
| CE 16:0 | 16:0 | Cholesteryl | 102.0849 | 63.55599 | 62.65006 | 122.335 | 92.20838 | 69.82915 |
| CE 16:1 | 16:1 | Cholesteryl | 90.77004 | 44.82555 | 92.34428 | 128.0056 | 69.78482 | 67.99116 |
| CE 18:0 | 18:0 | Cholesteryl | 93.28444 | 47.39088 | 49.99094 | 85.22585 | 56.28992 | 54.10493 |
| CE 18:1 | 18:1 | Cholesteryl | 535.5684 | 290.1185 | 423.1257 | 824.0723 | 538.0974 | 408.1206 |
| CE 18:2 | 18:2 | Cholesteryl | 441.7182 | 275.8611 | 401.4496 | 655.9555 | 493.3182 | 387.8593 |
| CE 18:3 | 18:3 | Cholesteryl | 52.4065 | 35.02608 | 31.69875 | 53.46213 | 24.47565 | 38.48925 |
| CE 20:4 | 20:4 | Cholesteryl | 90.5186 | 116.0675 | 107.9728 | 103.2384 | 139.9774 | 105.2141 |
| CE 22:6 | 22:6 | Cholesteryl | 89.76427 | 96.94046 | 84.66591 | 99.0688 | 120.6795 | 91.53769 |
| Total | | Cholesteryl | 1496.115 | 969.7861 | 1253.898 | 2071.363 | 1534.831 | 1223.146 |
| DAG 14:0-1 | 8:1 | Diacylglyce | 0.207254 | 0.072398 | 0.175645 | 0.099038 | 0.118707 | 0.11638 |
| DAG 16:0-1 | 8:0 | Diacylglyce | 9.64176 | 5.508772 | 8.089575 | 7.809388 | 9.298958 | 8.36281 |
| DAG 16:0-1 | 10:1 | Diacylglyce | 71.97636 | 47.78947 | 67.94595 | 56.53485 | 66.8214 | 64.73554 |
| DAG 16:0-1 | 10:2 | Diacylglyce | 51.05712 | 28.77193 | 41.81467 | 40.96728 | 40.85337 | 44.90083 |
| DAG 16:0-2 | 12:4 | Diacylglyce | 0.638923 | 0.517661 | 0.709807 | 0.686714 | 0.701987 | 0.628017 |
| DAG 16:1-1 | 8:1 | Diacylglyce | 2.077505 | 0.650292 | 1.624402 | 1.619232 | 1.427241 | 1.161074 |
| DAG 16:1-1 | 8:2 | Diacylglyce | 0.414366 | 0.103111 | 0.253205 | 0.310327 | 0.328978 | 0.148314 |
| DAG 16:1-1 | 10:1 | Diacylglyce | 0.688798 | 0.264094 | 0.522162 | 0.496728 | 0.5046 | 0.432149 |
| DAG 16:1-1 | 10:2 | Diacylglyce | 9.339199 | 2.087252 | 6.581467 | 4.811095 | 5.796525 | 3.989256 |
| DAG 16:1-1 | 10:3 | Diacylglyce | 5.372817 | 1.408889 | 3.301158 | 2.842105 | 3.157193 | 2.866116 |
| DAG 18:0-1 | 10:0 | Diacylglyce | 6.202495 | 4.238597 | 5.826255 | 4.572973 | 5.879083 | 5.21405 |
| DAG 18:0-1 | 12:0 | Diacylglyce | 3.290348 | 2.608187 | 2.886487 | 2.724324 | 2.866991 | 3.280165 |
| DAG 18:1-1 | 12:1 | Diacylglyce | 30.18516 | 16.70175 | 22.30193 | 15.76984 | 20.93454 | 19.70083 |
| DAG 18:1-1 | 12:2 | Diacylglyce | 90.29547 | 71.97661 | 84.16216 | 82.01138 | 86.91035 | 80.65289 |
| DAG 18:1-1 | 12:3 | Diacylglyce | 78.90217 | 57.38012 | 69.24324 | 64.34424 | 69.02293 | 70.21488 |
| DAG 18:2-1 | 12:2 | Diacylglyce | 27.70322 | 18.67135 | 21.88263 | 19.8 | 20.38666 | 18.57273 |
| DAG 18:2-1 | 12:4 | Diacylglyce | 45.00591 | 23.17193 | 28.28571 | 25.73258 | 26.71856 | 32.60331 |
| DAG 20:4-1 | 14:4 | Diacylglyce | 8.875903 | 9.291228 | 13.48494 | 12.16472 | 8.053092 | 7.990909 |
| DAG 18:1-18:3 | | Diacylglyce | 3.271438 | 1.294503 | 2.10417 | 1.858634 | 1.837276 | 2.250496 |
| DAG 18:1-20:4 | | Diacylglyce | 1.416835 | 0.784327 | 1.104093 | 0.659061 | 1.025962 | 0.924545 |
| DAG 18:1-22:6 | | Diacylglyce | 3.536179 | 2.19579 | 2.490348 | 1.648165 | 2.047922 | 2.791736 |
| DAG 18:2-20:4 | | Diacylglyce | 0.620486 | 0.318363 | 0.302317 | 0.265519 | 0.447811 | 0.446281 |
| DAG 18:2-20:5 | | Diacylglyce | 0.147498 | 0.065146 | 0.107514 | 0.07922 | 0.116131 | 0.112512 |
| DAG 18:2-22:6 | | Diacylglyce | 2.283152 | 1.544094 | 1.39668 | 1.085377 | 1.318416 | 1.81686 |
| DAG 20:4-20:4 | | Diacylglyce | 0.102398 | 0.121848 | 0.05495 | 0.096401 | 0.112228 | 0.08695 |
| DAG 16:1-18:1 (2) | | Diacylglyce | 10.52817 | 2.54269 | 7.54749 | 5.773827 | 6.171786 | 5.352893 |
| DAG 16:1-18:2 (2) | | Diacylglyce | 5.164806 | 1.29731 | 3.308108 | 3.044381 | 2.83697 | 2.975207 |
| DAG 16:1-18:3 (2) | | Diacylglyce | 0.186548 | 0.028959 | 0.118286 | 0.157161 | 0.081306 | 0.062678 |
| DAG 16:1-20:5 (2) | | Diacylglyce | 0.076822 | 0.031368 | 0.088425 | 0.07264 | 0.073551 | 0.079289 |
| DAG 16:1-22:6 (2) | | Diacylglyce | 1.133185 | 0.347368 | 0.653514 | 0.562532 | 0.601167 | 0.626529 |
| DAG 18:0-16:0 (2) | | Diacylglyce | 5.193171 | 3.466667 | 4.260232 | 3.981508 | 4.365532 | 4.990909 |

| | | | | | | | | |
|---|---|---|---|---|---|---|---|---|
| DAG 18:1-18:2 (2) | | Diacylglyce | 76.77479 | 48.63158 | 61.01931 | 55.20341 | 58.94093 | 62.20661 |
| DAG 16:0-18:3 | | Diacylglyce | 0.199926 | 0.149591 | 0.158896 | 0.157135 | 0.169017 | 0.16624 |
| DAG 18:1-20:4 (2) | | Diacylglyce | 10.47617 | 6.346199 | 7.932046 | 6.46771 | 8.130646 | 7.735537 |
| DAG 16:0-20:4 (2) | | Diacylglyce | 0.591412 | 0.452398 | 0.622471 | 0.614253 | 0.760028 | 0.553636 |
| DAG 16:0-20:5 | | Diacylglyce | 0.075593 | 0.077216 | 0.097969 | 0.101676 | 0.070949 | 0.099744 |
| DAG 16:0-22:6 | | Diacylglyce | 0.813132 | 0.494737 | 0.849498 | 0.785804 | 0.990938 | 0.739835 |
| DAG 18:2-20:4 (2) | | Diacylglyce | 4.394222 | 2.85848 | 2.453282 | 2.455989 | 3.660042 | 3.384298 |
| DAG 18:2-20:5 (2) | | Diacylglyce | 1.037452 | 0.370292 | 0.542548 | 0.703869 | 0.745768 | 0.809504 |
| DAG 18:2-22:6 (2) | | Diacylglyce | 14.84675 | 11.73801 | 9.637066 | 6.055477 | 7.465184 | 13.46033 |
| Total | | Diacylglyce | 634.855 | 408.9982 | 524.9709 | 466.205 | 503.4964 | 516.543 |
| DAG 16:1-22:6 | | Diacylglyce | 0.142629 | 0.04945 | 0.078857 | 0.067442 | 0.065796 | 0.06776 |
| DAG 18:0-18:3 | | Diacylglyce | 0.106085 | 0.053099 | 0.068108 | 0.081883 | 0.052912 | 0.072893 |
| DAG 18:0-20:4 | | Diacylglyce | 7.074721 | 7.307602 | 11.31892 | 9.811664 | 6.784712 | 6.468595 |
| DAG 18:0-20:5 | | Diacylglyce | 0.097529 | 0.048257 | 0.105151 | 0.071309 | 0.069673 | 0.084397 |
| DAG 18:0-22:6 | | Diacylglyce | 0.91501 | 0.785497 | 0.929653 | 0.973229 | 1.156053 | 0.961736 |
| DAG 18:1-20:5 | | Diacylglyce | 0.231624 | 0.113404 | 0.179212 | 0.136011 | 0.127739 | 0.157264 |
| DAG 18:2-18:3 | | Diacylglyce | 1.585844 | 0.600702 | 0.875676 | 0.907169 | 0.837331 | 0.957025 |
| DAG 18:1-22:6 (2) | | Diacylglyce | 28.83782 | 18.71111 | 18.37761 | 12.00854 | 16.47394 | 22.74793 |
| DAG 16:1-20:4 (2) | | Diacylglyce | 0.303506 | 0.158035 | 0.256911 | 0.231081 | 0.279444 | 0.21481 |
| DAG 18:1-18:3 (2) | | Diacylglyce | 6.902167 | 2.977778 | 4.204633 | 4.109531 | 3.552467 | 4.490083 |
| DAG 18:1-20:5 (2) | | Diacylglyce | 1.398162 | 0.662222 | 0.992896 | 0.897952 | 0.908881 | 1.255537 |
| DAG 18:2-18:3 (2) | | Diacylglyce | 2.515036 | 1.160468 | 1.642703 | 1.782589 | 1.436748 | 1.822066 |
| LPC 16:0 | 16:0 | Lysophospl | 118.4278 | 142.2489 | 98.46515 | 128.941 | 136.1336 | 128.6132 |
| LPC 16:1 | 16:1 | Lysophospl | 9.505348 | 4.630196 | 7.355833 | 7.913532 | 8.059584 | 5.197213 |
| LPC 20:6 | | Lysophospl | 0.39385 | 0.54307 | 0.442671 | 0.36283 | 0.379321 | 0.36878 |
| Total | | Lysophospl | 476.2888 | 507.5103 | 431.7325 | 486.7379 | 529.1874 | 485.7816 |
| LPC 14:0 | | Lysophospl | 1.318877 | 0.727277 | 0.879927 | 0.720404 | 0.71586 | 0.959582 |
| LPC 18:3 | | Lysophospl | 3.216578 | 3.149716 | 4.325018 | 3.73543 | 2.355969 | 3.217422 |
| LPC 20:0 | | Lysophospl | 3.391444 | 4.734807 | 2.841966 | 2.878383 | 1.469376 | 3.760976 |
| LPC 20:1 | | Lysophospl | 2.475401 | 1.868452 | 2.070726 | 1.371881 | 1.615663 | 1.9223 |
| LPC 20:2 | | Lysophospl | 3.296791 | 1.961011 | 3.25796 | 1.87863 | 3.842497 | 2.205575 |
| LPC 20:3 | | Lysophospl | 16.82888 | 12.41693 | 18.27733 | 12.44537 | 15.2931 | 15.36585 |
| LPC 20:5 | | Lysophospl | 6.192513 | 3.429438 | 7.369039 | 4.206401 | 2.961227 | 4.034843 |
| LPC 18:0 | | Lysophospl | 81.40107 | 101.7233 | 70.45488 | 91.95059 | 99.48302 | 90.52265 |
| LPC 18:2 | | Lysophospl | 99.52941 | 112.8667 | 87.71533 | 113.6597 | 110.1884 | 107.9791 |
| LPC 14:2 | | Lysophospl | 0.051289 | 0.069203 | 0.072198 | 0.075052 | 0.079295 | 0.040976 |
| LPC 18:1 | | Lysophospl | 68.24599 | 52.55591 | 60.65591 | 57.79001 | 61.03834 | 58.5993 |
| LPC 16:2 | | Lysophospl | 0.136524 | 0.15835 | 0.144343 | 0.129244 | 0.106679 | 0.171449 |
| LPC 20:4 | | Lysophospl | 61.87701 | 64.42704 | 67.40426 | 58.67939 | 85.4655 | 62.8223 |
| PC 32:0 | 32:0 | Phosphatid | 51.93392 | 52.94314 | 49.90603 | 57.10685 | 68.39324 | 64.0447 |
| PC 32:1 | 32:1 | Phosphatid | 44.14127 | 39.05933 | 40.80513 | 46.54247 | 45.64809 | 49.53569 |
| PC 36:0 | 36:0 | Phosphatid | 51.99544 | 56.63659 | 48.87774 | 56.39452 | 56.3444 | 64.74549 |
| PC 36:1 | 36:1 | Phosphatid | 57.33751 | 59.95179 | 52.38548 | 57.51233 | 61.55166 | 68.56092 |
| PC 36:2 | 36:2 | Phosphatid | 75.07605 | 83.38072 | 68.02136 | 78.78356 | 82.36944 | 90.48017 |
| PC 34:1 | | Phosphatid | 78.03931 | 85.07169 | 76.52643 | 85.55616 | 85.57045 | 98.708 |
| PC 40:5 | | Phosphatid | 49.05269 | 53.85538 | 44.92792 | 47.8137 | 56.40075 | 58.69791 |
| PC 40:6 | | Phosphatid | 65.65309 | 70.22002 | 62.63962 | 66.53151 | 70.05009 | 79.26748 |
| PC 38:2 | | Phosphatid | 55.98405 | 59.27318 | 51.26108 | 59.94521 | 59.25235 | 72.62293 |
| PC 34:0 | | Phosphatid | 63.29479 | 70.4759 | 58.03631 | 67.03562 | 71.31246 | 80.5912 |

| | | | | | | | | |
|---|---|---|---|---|---|---|---|---|
| PC 34:2 | | Phosphatid | 116.9923 | 137.6143 | 105.5206 | 134.3562 | 122.9681 | 160.274 |
| PC 34:3 | | Phosphatid | 39.41441 | 41.29543 | 39.62306 | 44.35068 | 42.07514 | 48.73107 |
| PC 40:1 | | Phosphatid | 12.11962 | 13.47219 | 12.15697 | 12.12055 | 14.07765 | 16.05335 |
| PC 36:3 | | Phosphatid | 79.25947 | 81.16687 | 74.89269 | 83.36438 | 80.5886 | 97.22855 |
| PC 36:4 | | Phosphatid | 72.27684 | 75.8492 | 73.04752 | 76.93151 | 85.01816 | 90.90844 |
| PC 36:5 | | Phosphatid | 35.26175 | 37.42398 | 32.68446 | 37.40274 | 43.64183 | 38.77722 |
| PC 38:0 | | Phosphatid | 37.6303 | 38.09147 | 34.36626 | 37.16164 | 35.22229 | 43.28046 |
| PC 38:6 | | Phosphatid | 80.30533 | 79.53152 | 79.43833 | 84.96438 | 83.13588 | 91.92069 |
| PC 40:0 | | Phosphatid | 10.70464 | 11.12485 | 9.629471 | 11.27671 | 11.25423 | 13.44485 |
| PC 40:2 | | Phosphatid | 17.88208 | 17.33251 | 15.78964 | 12.99726 | 12.95053 | 19.89474 |
| PC 40:3 | | Phosphatid | 29.81715 | 28.8356 | 26.30326 | 22.68493 | 20.93049 | 32.04182 |
| PC 38:3 | | Phosphatid | 56.5685 | 59.80717 | 52.11639 | 59.39726 | 60.16531 | 68.65177 |
| PC 38:5 | | Phosphatid | 72.32811 | 75.4932 | 67.75227 | 75.35342 | 73.99499 | 88.11824 |
| PC 38:1 | | Phosphatid | 74.75819 | 78.51916 | 65.10945 | 75.55069 | 77.61302 | 91.06417 |
| PC 30:0 | | Phosphatid | 8.702136 | 8.118912 | 7.668019 | 9.775343 | 17.60551 | 11.3801 |
| PC 40:4 | | Phosphatid | 46.4278 | 48.8937 | 42.52536 | 40.74521 | 41.3087 | 53.22134 |
| PC 38:4 | | Phosphatid | 67.34492 | 69.02967 | 60.20822 | 65.9726 | 72.48466 | 78.82624 |
| Total | | Phosphatid | 1450.302 | 1532.467 | 1352.219 | 1507.627 | 1551.928 | 1771.072 |
| SM 16:0 | 16:0 | Sphingomy | 33.63144 | 31.76143 | 32.17512 | 35.23288 | 39.95617 | 36.71377 |
| SM 16:1 | 16:1 | Sphingomy | 11.11478 | 8.026576 | 9.592953 | 11.91233 | 18.73262 | 10.75068 |
| SM 12:0 | 12:0 | Sphingomy | 0.595933 | 0.883535 | 0.664645 | 0.792986 | 0.688779 | 0.782422 |
| SM 24:0 | 24:0 | Sphingomy | 72.56394 | 72.55624 | 60.95782 | 72.08767 | 75.201 | 84.43259 |
| Total | | Sphingomy | 147.1183 | 138.2809 | 131.3756 | 152.4094 | 163.8722 | 166.7847 |
| SM 18:0 | | Sphingomy | 29.21219 | 25.05315 | 27.98505 | 32.38356 | 29.29368 | 34.10526 |
| TAG 14:0-1 | 18:0 | Triacylglyce | 2.27E-05 | 2.79E-05 | 2.82E-05 | 3.63E-05 | 5.31E-05 | 2.51E-05 |
| TAG 14:0-1 | 22:1 | Triacylglyce | 0.000298 | 0.000273 | 0.000188 | 0.000468 | 0.000443 | 0.000271 |
| TAG 14:0-1 | 26:2 | Triacylglyce | 0.007024 | 0.00435 | 0.00605 | 0.008246 | 0.009003 | 0.004842 |
| TAG 16:0-1 | 24:0 | Triacylglyce | 0.008844 | 0.010612 | 0.008879 | 0.012478 | 0.01309 | 0.007086 |
| TAG 16:1-1 | 24:3 | Triacylglyce | 0.00507 | 0.002393 | 0.003581 | 0.007037 | 0.00728 | 0.002719 |
| TAG 16:1-1 | 26:2 | Triacylglyce | 0.001063 | 0.000801 | 0.000762 | 0.001314 | 0.001291 | 0.00074 |
| TAG 16:1-1 | 26:3 | Triacylglyce | 0.01794 | 0.009133 | 0.014202 | 0.022163 | 0.02196 | 0.011948 |
| TAG 16:1-1 | 26:4 | Triacylglyce | 0.019852 | 0.011333 | 0.016146 | 0.026943 | 0.025448 | 0.016079 |
| TAG 16:1-1 | 28:1 | Triacylglyce | 0.000887 | 0.00054 | 0.000709 | 0.000969 | 0.000857 | 0.000553 |
| TAG 16:1-1 | 28:3 | Triacylglyce | 0.039509 | 0.017444 | 0.028599 | 0.045594 | 0.047127 | 0.020664 |
| TAG 16:1-1 | 28:5 | Triacylglyce | 0.027816 | 0.015032 | 0.021973 | 0.037418 | 0.034575 | 0.021198 |
| TAG 18:0-1 | 26:0 | Triacylglyce | 0.00396 | 0.004164 | 0.004512 | 0.005235 | 0.005256 | 0.003862 |
| TAG 18:0-1 | 28:0 | Triacylglyce | 0.005873 | 0.005084 | 0.005883 | 0.007082 | 0.007592 | 0.005593 |
| TAG 18:0-1 | 30:0 | Triacylglyce | 0.002355 | 0.002271 | 0.002643 | 0.002144 | 0.001918 | 0.002384 |
| TAG 18:1-1 | 26:1 | Triacylglyce | 0.036977 | 0.025651 | 0.033199 | 0.054183 | 0.054889 | 0.031638 |
| TAG 18:1-1 | 30:1 | Triacylglyce | 0.017233 | 0.013391 | 0.015964 | 0.016116 | 0.014994 | 0.014734 |
| TAG 18:1-1 | 28:2 | Triacylglyce | 0.096374 | 0.050916 | 0.07683 | 0.150988 | 0.147521 | 0.068767 |
| TAG 18:1-1 | 30:2 | Triacylglyce | 0.040274 | 0.030208 | 0.032557 | 0.045023 | 0.043125 | 0.03009 |
| TAG 18:1-1 | 30:3 | Triacylglyce | 0.094282 | 0.049337 | 0.066574 | 0.127753 | 0.129266 | 0.058428 |
| TAG 18:1-1 | 30:4 | Triacylglyce | 0.073651 | 0.041877 | 0.052743 | 0.106239 | 0.10438 | 0.051694 |
| TAG 18:1-1 | 30:5 | Triacylglyce | 0.057088 | 0.032856 | 0.041536 | 0.082222 | 0.078402 | 0.041193 |
| TAG 18:2-1 | 26:2 | Triacylglyce | 0.02923 | 0.019272 | 0.024075 | 0.04949 | 0.047541 | 0.027723 |
| TAG 18:2-1 | 30:2 | Triacylglyce | 0.012097 | 0.011004 | 0.01037 | 0.013331 | 0.012287 | 0.012408 |
| TAG 18:2-1 | 28:4 | Triacylglyce | 0.069539 | 0.035305 | 0.053762 | 0.130569 | 0.122635 | 0.052201 |
| TAG 18:2-1 | 30:4 | Triacylglyce | 0.015798 | 0.012459 | 0.011666 | 0.022265 | 0.019464 | 0.012151 |

| | | | | | | | | |
|---|---|---|---|---|---|---|---|---|
| TAG 18:2-1 | 6:6 | Triacylglyce | 0.039386 | 0.025918 | 0.030128 | 0.058893 | 0.054983 | 0.03199 |
| TAG 16:0-16:0-20:4 | | Triacylglyce | 0.001693 | 0.001338 | 0.001824 | 0.002628 | 0.00343 | 0.001716 |
| TAG 16:0-16:0-20:5 | | Triacylglyce | 0.000622 | 0.000406 | 0.000622 | 0.001207 | 0.00096 | 0.00066 |
| TAG 16:0-20:5-20:5 | | Triacylglyce | 0.000628 | 0.000399 | 0.000527 | 0.000693 | 0.000583 | 0.000592 |
| TAG 16:1-16:1-20:5 | | Triacylglyce | 0.001484 | 0.000484 | 0.001311 | 0.003671 | 0.00222 | 0.001451 |
| TAG 18:2-20:4-20:4 | | Triacylglyce | 0.01336 | 0.01078 | 0.010225 | 0.013761 | 0.014112 | 0.012644 |
| TAG 18:1-18:1-20:4 | | Triacylglyce | 0.011686 | 0.008399 | 0.00942 | 0.015592 | 0.016936 | 0.008642 |
| Total | | Triacylglyce | 0.825549 | 0.503737 | 0.647987 | 1.166401 | 1.139197 | 0.623073 |
| TAG 18:2-18:2-20:5 | | Triacylglyce | 0.009652 | 0.005382 | 0.007721 | 0.015459 | 0.011616 | 0.009135 |
| TAG 16:1-20:4-20:4 | | Triacylglyce | 0.006942 | 0.002354 | 0.005444 | 0.007862 | 0.007607 | 0.005359 |
| TAG 18:2-20:5-20:5 | | Triacylglyce | 0.000516 | 0.000282 | 0.000379 | 0.000601 | 0.000424 | 0.000415 |
| TAG 18:1-18:1-20:5 | | Triacylglyce | 0.005773 | 0.003482 | 0.004606 | 0.007972 | 0.006317 | 0.004802 |
| TAG 18:1-20:4-20:4 | | Triacylglyce | 0.009839 | 0.007628 | 0.007582 | 0.008817 | 0.009057 | 0.007932 |
| TAG 18:0-18:0-20:5 | | Triacylglyce | 0.000143 | 0.000124 | 0.000124 | 0.000222 | 0.000136 | 0.000152 |
| TAG 16:1-16:1-20:4 | | Triacylglyce | 0.001736 | 0.000812 | 0.001413 | 0.002452 | 0.003037 | 0.001623 |
| TAG 18:1-20:5-20:5 | | Triacylglyce | 0.00073 | 0.000379 | 0.000499 | 0.000594 | 0.000486 | 0.000486 |
| TAG 18:2-18:2-20:4 | | Triacylglyce | 0.008952 | 0.007187 | 0.006953 | 0.010342 | 0.013754 | 0.008466 |
| TAG 20:4-20:4-20:4 | | Triacylglyce | 0.001595 | 0.001347 | 0.001275 | 0.001404 | 0.002019 | 0.001492 |
| TAG 16:1-20:5-20:5 | | Triacylglyce | 7.78E-05 | 2.31E-05 | 4.58E-05 | 8.11E-05 | 6.96E-05 | 4.53E-05 |
| TAG 18:0-20:5-20:5 | | Triacylglyce | 0.000145 | 0.000103 | 0.000134 | 0.000106 | 8.99E-05 | 0.000121 |
| TAG 18:0-20:4-20:4 | | Triacylglyce | 0.001618 | 0.001506 | 0.001275 | 0.001835 | 0.002074 | 0.00144 |
| TAG 18:0-18:0-20:4 | | Triacylglyce | 0.002105 | 0.001984 | 0.002183 | 0.002998 | 0.003048 | 0.002212 |
| TAG 16:0-20:4-20:4 | | Triacylglyce | 0.023812 | 0.017687 | 0.020866 | 0.033906 | 0.035839 | 0.022706 |

| | | | | | | | | |
|---|---|---|---|---|---|---|---|---|
| 29 (Control | 36 (Control | 3 (Control - | 7 (Control - | 12 (Control | 17 (Control | 19 (Control | 28 (Control | 32 (Control |
| CON-Veh-I | CON-Veh-I | CON-GH | CON-GH | CON-GH | CON-GH | CON-GH-Ir | CON-GH-Ir | CON-GH-Ir |
| 15.54545 | 18.46664 | 31.52932 | 24.76519 | 27.49257 | 29.48475 | 21.92308 | 34.5485 | 28.55918 |
| 4.733182 | 7.931322 | 11.29536 | 6.852963 | 8.842769 | 9.187801 | 6.657692 | 9.478585 | 12.66792 |
| 428.7273 | 530.5907 | 492.6025 | 433.1883 | 438.9416 | 534.9438 | 489.0865 | 527.4705 | 534.9057 |
| 376.7727 | 440.782 | 468.0414 | 380.6602 | 430.0689 | 448.6998 | 409.4712 | 438.4633 | 482.8816 |
| 28.70455 | 40.0697 | 32.81257 | 28.22206 | 33.82095 | 46.76244 | 36.15865 | 42.47699 | 43.53345 |
| 0.597818 | 0.749528 | 1.145818 | 0.891493 | 1.063298 | 0.908957 | 0.78101 | 1.137102 | 0.883791 |
| 855.081 | 1038.59 | 1037.427 | 874.5802 | 940.2301 | 1069.988 | 964.0781 | 1053.575 | 1103.432 |
| 76.85087 | 61.54 | 83.77516 | 126.3839 | 77.91319 | 61.37119 | 83.80012 | 50.19852 | 134.9045 |
| 67.44209 | 62.35573 | 70.5177 | 130.0117 | 82.46179 | 47.73258 | 75.28686 | 25.65032 | 131.341 |
| 65.7184 | 56.73952 | 73.07425 | 112.8006 | 57.56919 | 45.46884 | 67.5731 | 23.10596 | 117.2567 |
| 465.1249 | 486.8338 | 620.9022 | 878.2751 | 510.5441 | 306.8048 | 472.3011 | 195.8001 | 1023.238 |
| 381.2566 | 460.3409 | 482.2484 | 661.8702 | 433.9515 | 276.5025 | 403.5287 | 200.3205 | 788.5553 |
| 29.45269 | 37.25607 | 30.02915 | 53.10148 | 36.61626 | 35.72917 | 43.10672 | 23.76896 | 56.26962 |
| 102.4678 | 123.8741 | 129.077 | 139.0391 | 137.7053 | 135.5685 | 186.4035 | 110.9017 | 137.1954 |
| 79.64421 | 75.43613 | 75.00625 | 113.9817 | 75.05197 | 91.52023 | 118.3714 | 98.71801 | 114.7961 |
| 1267.958 | 1364.376 | 1564.63 | 2215.464 | 1411.813 | 1000.698 | 1450.371 | 728.464 | 2503.557 |
| 0.13508 | 0.075925 | 0.292873 | 0.193019 | 0.187179 | 0.045728 | 0.080418 | 0.063832 | 0.06887 |
| 7.031984 | 4.240459 | 12.43801 | 10.65242 | 8.44264 | 4.430964 | 5.302865 | 6.017418 | 5.699233 |
| 63.00618 | 40.88092 | 86.28078 | 80.30161 | 73.61856 | 39.26812 | 48.03997 | 47.27578 | 42.21483 |
| 38.12491 | 29.18221 | 60.57019 | 56.21376 | 46.49456 | 25.41027 | 29.14057 | 24.87548 | 28.45013 |
| 0.621908 | 0.69934 | 0.923715 | 0.697335 | 0.716345 | 0.62601 | 0.551632 | 0.563239 | 0.474859 |
| 1.067893 | 0.654146 | 2.620302 | 2.229312 | 1.8838 | 0.484898 | 0.820253 | 0.602842 | 1.023376 |
| 0.207698 | 0.142502 | 0.660389 | 0.479649 | 0.385062 | 0.060093 | 0.139755 | 0.075163 | 0.246982 |
| 0.465257 | 0.279684 | 1.116285 | 0.939268 | 0.770123 | 0.27361 | 0.334097 | 0.28217 | 0.404194 |
| 4.902128 | 2.616069 | 12.62981 | 9.693119 | 7.77955 | 1.473948 | 3.168288 | 1.737021 | 3.482609 |
| 2.705559 | 1.88858 | 7.630238 | 5.152269 | 4.409282 | 1.050359 | 1.923278 | 1.263713 | 2.113044 |
| 5.020728 | 3.336585 | 9.926566 | 8.617862 | 6.646556 | 3.655735 | 4.01972 | 4.513063 | 3.692072 |
| 3.130542 | 2.19254 | 4.989201 | 4.359004 | 3.380711 | 2.161267 | 2.914057 | 3.003209 | 2.432992 |
| 19.61098 | 9.565567 | 46.31534 | 36.13177 | 27.77665 | 11.16481 | 17.29727 | 13.84446 | 12.94757 |
| 82.6987 | 70.70875 | 100.5097 | 100.1991 | 94.50326 | 68.02252 | 76.38907 | 77.94041 | 72.78261 |
| 68.68909 | 45.09039 | 91.02376 | 85.15081 | 77.45613 | 44.41098 | 59.98401 | 46.58824 | 48.0844 |
| 19.41826 | 14.02812 | 38.17711 | 31.94144 | 27.01958 | 15.61858 | 20.72219 | 20.03789 | 18.52941 |
| 25.42485 | 17.0858 | 43.07559 | 41.4817 | 30.88325 | 16.90809 | 20.69101 | 16.33889 | 15.33913 |
| 10.08099 | 11.64706 | 12.76976 | 10.49429 | 13.41581 | 15.3855 | 8.977215 | 10.68449 | 10.08414 |
| 1.720439 | 1.105567 | 3.649244 | 3.238946 | 2.511124 | 0.960929 | 1.401386 | 1.092101 | 1.23399 |
| 0.887028 | 0.568924 | 1.667041 | 1.398887 | 1.125163 | 0.621956 | 0.694097 | 0.553338 | 0.567621 |
| 2.638847 | 1.198795 | 4.69892 | 4.619912 | 3.015228 | 1.291795 | 1.882745 | 2.141849 | 1.056522 |
| 0.395086 | 0.165151 | 0.581339 | 0.58981 | 0.445627 | 0.22735 | 0.341292 | 0.273727 | 0.223159 |
| 0.054803 | 0.102551 | 0.165745 | 0.172621 | 0.092911 | 0.060093 | 0.063102 | 0.07233 | 0.080724 |
| 1.651503 | 0.808321 | 2.364492 | 2.711859 | 1.846991 | 1.045292 | 1.30497 | 1.659465 | 0.780997 |
| 0.064983 | 0.050617 | 0.124302 | 0.126395 | 0.08346 | 0.080994 | 0.085335 | 0.077995 | 0.05934 |
| 5.967056 | 2.969871 | 15.07387 | 11.52738 | 8.735025 | 1.594032 | 4.007728 | 2.150374 | 4.074169 |
| 2.399424 | 1.887805 | 7.332181 | 5.761054 | 4.106454 | 1.229472 | 1.865716 | 1.094851 | 2.080818 |
| 0.091742 | 0.067274 | 0.217892 | 0.150878 | 0.137343 | 0.028754 | 0.063102 | 0.028354 | 0.078353 |
| 0.042054 | 0.034631 | 0.128346 | 0.116908 | 0.092911 | 0.031364 | 0.048256 | 0.029757 | 0.046289 |
| 0.598929 | 0.27297 | 1.483801 | 1.266852 | 0.858883 | 0.282224 | 0.393338 | 0.382827 | 0.483146 |
| 3.6593 | 2.597991 | 7.959395 | 6.973353 | 4.871356 | 2.756369 | 3.257029 | 3.555997 | 2.805882 |

| | | | | | | | | |
|---|---|---|---|---|---|---|---|---|
| 59.34935 | 38.50502 | 87.26566 | 82.98975 | 70.92966 | 35.77199 | 49.11925 | 39.13522 | 39.13044 |
| 0.124876 | 0.115877 | 0.244613 | 0.198422 | 0.162927 | 0.088847 | 0.186787 | 0.199994 | 0.110394 |
| 7.822649 | 4.911908 | 12.48726 | 10.77628 | 8.693256 | 4.914849 | 5.979214 | 4.128037 | 4.290537 |
| 0.535182 | 0.582095 | 0.820821 | 0.592709 | 0.636983 | 0.563181 | 0.498628 | 0.577265 | 0.48913 |
| 0.038224 | 0.073265 | 0.120311 | 0.081567 | 0.091553 | 0.04322 | 0.060608 | 0.03831 | 0.072506 |
| 0.815621 | 0.683329 | 1.505313 | 1.098975 | 0.842959 | 0.508205 | 0.541799 | 0.683697 | 0.75867 |
| 2.437474 | 1.722009 | 4.558963 | 4.580381 | 3.636548 | 1.847375 | 2.283278 | 2.066494 | 1.74844 |
| 0.47267 | 0.583386 | 0.864881 | 0.973265 | 0.791791 | 0.362027 | 0.43531 | 0.453782 | 0.48913 |
| 10.18971 | 5.27604 | 14.20821 | 19.53119 | 11.10805 | 7.169599 | 8.073018 | 10.30222 | 4.147826 |
| 491.1582 | 345.0722 | 758.9482 | 703.7574 | 594.1982 | 342.4352 | 411.0428 | 377.4868 | 355.3572 |
| 0.099402 | 0.066603 | 0.213849 | 0.197078 | 0.111733 | 0.035291 | 0.077924 | 0.04106 | 0.054621 |
| 0.109606 | 0.098574 | 0.143041 | 0.122627 | 0.075394 | 0.06402 | 0.055667 | 0.097879 | 0.051054 |
| 8.405765 | 10.9033 | 11.44536 | 8.860322 | 11.0454 | 14.17959 | 7.921919 | 9.400153 | 8.454476 |
| 0.105776 | 0.05728 | 0.102946 | 0.069338 | 0.070016 | 0.073165 | 0.060608 | 0.082286 | 0.054621 |
| 0.916184 | 0.953716 | 1.558704 | 1.076574 | 1.063031 | 0.939395 | 0.902998 | 1.106402 | 0.949719 |
| 0.117241 | 0.107897 | 0.201849 | 0.203903 | 0.177729 | 0.091457 | 0.11131 | 0.079426 | 0.103281 |
| 0.758051 | 0.498164 | 1.450108 | 1.508521 | 1.180769 | 0.491232 | 0.484957 | 0.50356 | 0.582813 |
| 20.26328 | 9.782496 | 32.94169 | 35.81552 | 21.82712 | 11.33709 | 14.34484 | 16.14912 | 7.936573 |
| 0.252272 | 0.158488 | 0.485184 | 0.353411 | 0.362088 | 0.145039 | 0.194199 | 0.214185 | 0.221985 |
| 3.772958 | 2.229469 | 7.241469 | 6.910103 | 4.735606 | 1.810894 | 2.417588 | 1.884981 | 2.449105 |
| 0.793878 | 0.661894 | 1.192225 | 1.552006 | 1.028571 | 0.471724 | 0.517095 | 0.627594 | 0.601918 |
| 1.264077 | 0.956298 | 2.499525 | 2.68287 | 1.935489 | 0.864912 | 0.872059 | 0.894912 | 1.018542 |
| 132.4221 | 142.7464 | 114.3904 | 121.7342 | 125.3517 | 132.2124 | 136.2414 | 126.4776 | 158.9309 |
| 7.545729 | 6.573186 | 8.52749 | 9.743522 | 7.434534 | 4.863092 | 5.455862 | 3.655102 | 10.26201 |
| 0.447839 | 0.427696 | 0.41259 | 0.311561 | 0.338263 | 0.374055 | 0.406552 | 0.362755 | 0.447926 |
| 499.8131 | 480.7088 | 498.4556 | 457.8868 | 475.9027 | 466.3899 | 542.5653 | 454.4367 | 579.0228 |
| 0.746332 | 0.706837 | 0.95506 | 1.276944 | 0.70322 | 0.739302 | 0.841448 | 0.724592 | 1.736722 |
| 3.790955 | 3.106907 | 3.168128 | 2.38804 | 3.792585 | 2.082041 | 3.026897 | 2.428163 | 6.849243 |
| 4.090452 | 2.925131 | 2.34741 | 2.093023 | 1.747564 | 2.612389 | 3.02069 | 4.933469 | 2.490849 |
| 1.759196 | 1.699292 | 2.172112 | 1.397143 | 1.346186 | 1.396523 | 2.030483 | 1.486286 | 1.334773 |
| 2.601005 | 1.925572 | 2.924303 | 2.338206 | 2.070763 | 2.068922 | 3.343448 | 2.558571 | 1.934852 |
| 16.36985 | 11.87185 | 17.43426 | 17.27043 | 12.80212 | 10.59761 | 17.20966 | 15.9649 | 14.02554 |
| 4.367839 | 3.825653 | 4.363347 | 3.685714 | 3.836441 | 3.183967 | 4.706897 | 5.947347 | 3.55023 |
| 84.54271 | 95.73535 | 86.56574 | 97.09635 | 90.74364 | 97.2431 | 100.6552 | 96.28163 | 115.0388 |
| 105.2462 | 96.52931 | 107.8884 | 78.19934 | 102.2606 | 105.8824 | 112.6552 | 89.48571 | 119.4944 |
| 0.080864 | 0.070036 | 0.041084 | 0.051409 | 0.042292 | 0.047357 | 0.046945 | 0.039784 | 0.067213 |
| 64.1407 | 52.10911 | 65.4502 | 53.10299 | 60.63559 | 49.68038 | 72.08276 | 53.74286 | 60.6241 |
| 0.163799 | 0.177932 | 0.205418 | 0.161322 | 0.10228 | 0.109218 | 0.214345 | 0.131676 | 0.210217 |
| 71.49749 | 60.27858 | 81.60956 | 67.03655 | 62.69492 | 53.29724 | 80.62759 | 50.21633 | 82.02502 |
| 56.30488 | 58.39289 | 54.25737 | 53.14392 | 51.45614 | 50.86047 | 51.51142 | 47.31118 | 52.6366 |
| 44.96707 | 45.09426 | 45.71695 | 44.32577 | 43.88039 | 38.72093 | 38.7522 | 36.43505 | 43.9966 |
| 55.04268 | 58.18724 | 50.8433 | 54.96798 | 49.24231 | 51.9593 | 52.08084 | 50.663 | 51.55404 |
| 58.8622 | 62.07172 | 57.33528 | 59.47777 | 53.26375 | 56.16628 | 59.74692 | 52.56138 | 56.20085 |
| 74.32683 | 81.59695 | 78.3344 | 86.3351 | 75.02623 | 76.0814 | 82.53427 | 77.03268 | 81.38553 |
| 85.55488 | 86.66963 | 82.2527 | 84.41032 | 80.64203 | 79.61861 | 83.67311 | 77.13156 | 80.16 |
| 49.81829 | 52.92034 | 49.27809 | 50.12247 | 47.39408 | 49.81395 | 49.56063 | 49.98077 | 47.89787 |
| 68.74024 | 74.23929 | 67.24132 | 70.01927 | 64.48519 | 65.2186 | 68.8471 | 64.8613 | 64.64681 |
| 56.76585 | 62.25452 | 57.90254 | 64.44638 | 54.86827 | 56.44884 | 56.21441 | 51.59242 | 55.5983 |
| 68.41098 | 71.80578 | 64.35249 | 66.65092 | 62.7385 | 65.57442 | 65.97891 | 61.43038 | 61.65447 |

| | | | | | | | | |
|---|---|---|---|---|---|---|---|---|
| 128.3049 | 137.0993 | 116.7085 | 126.9008 | 122.5726 | 124.3256 | 124.1125 | 119.1431 | 133.6851 |
| 41.58659 | 46.06538 | 40.03385 | 42.71433 | 39.69647 | 39.38023 | 41.50439 | 37.72041 | 41.55574 |
| 13.73049 | 13.92701 | 11.5553 | 11.50389 | 11.05896 | 11.37558 | 12.67487 | 12.48778 | 11.09106 |
| 84.44634 | 87.19518 | 79.66851 | 81.60149 | 75.82849 | 77.53605 | 79.58172 | 75.6089 | 78.08681 |
| 80.22073 | 83.0365 | 75.68719 | 76.75598 | 70.89309 | 71.86395 | 73.25483 | 65.66218 | 77.32085 |
| 36.27439 | 35.44018 | 37.14503 | 39.63693 | 33.97913 | 34.15814 | 35.89455 | 38.24444 | 33.64085 |
| 40.68659 | 42.23802 | 33.97257 | 31.57973 | 31.0646 | 35.24651 | 38.37258 | 36.2373 | 31.97617 |
| 85.23659 | 84.83021 | 76.00233 | 78.60243 | 71.27898 | 75.32791 | 74.41476 | 78.64433 | 78.78128 |
| 12.02927 | 12.96731 | 9.090867 | 9.648492 | 9.224937 | 9.800581 | 12.12654 | 11.67701 | 10.76426 |
| 18.59268 | 17.43447 | 13.79282 | 13.19366 | 11.18082 | 12.17093 | 15.2478 | 18.13348 | 10.69277 |
| 31.01707 | 28.09394 | 25.32711 | 26.82375 | 20.37123 | 22.02907 | 25.72935 | 27.37819 | 20.02723 |
| 58.32439 | 63.88829 | 57.28276 | 59.11968 | 54.41128 | 57.98721 | 57.84886 | 53.0953 | 56.13957 |
| 74.7878 | 78.44367 | 70.98103 | 72.54834 | 68.7402 | 71.05814 | 73.32865 | 67.52101 | 69.39574 |
| 77.09268 | 77.20978 | 72.37817 | 78.16599 | 68.26291 | 69.26861 | 74.49912 | 69.79511 | 71.87745 |
| 9.534512 | 10.04367 | 9.626612 | 9.038607 | 9.631143 | 8.898488 | 8.72478 | 8.357814 | 8.644085 |
| 48.04024 | 47.36782 | 44.04669 | 45.06435 | 36.91396 | 39.47442 | 43.42355 | 42.55534 | 37.22553 |
| 70.33171 | 73.78229 | 69.69945 | 73.07429 | 66.11001 | 66.34884 | 71.1775 | 64.63389 | 66.43404 |
| 1529.031 | 1592.296 | 1450.513 | 1509.873 | 1384.216 | 1416.713 | 1470.816 | 1395.895 | 1433.07 |
| 35.96707 | 38.02222 | 37.50219 | 35.82095 | 35.40085 | 34.70233 | 34.6819 | 32.12414 | 35.56085 |
| 12.09512 | 13.41288 | 10.53633 | 11.16481 | 10.42934 | 9.658256 | 11.5993 | 9.166603 | 9.682723 |
| 0.578524 | 0.808658 | 0.380274 | 0.600821 | 0.556705 | 0.806965 | 0.72 | 0.869399 | 0.79057 |
| 71.17683 | 71.92003 | 68.19726 | 77.4498 | 66.30296 | 62.57093 | 67.07557 | 64.16918 | 67.02638 |
| 149.7261 | 155.1025 | 148.8553 | 151.9161 | 143.8662 | 134.0582 | 141.4511 | 130.8995 | 143.7601 |
| 29.90854 | 30.93875 | 32.23928 | 26.8797 | 31.1763 | 26.31977 | 27.37434 | 24.57017 | 30.69957 |
| 1.86E-05 | 3.63E-05 | 0.000119 | 0.00013 | 6.25E-05 | 2.1E-05 | 1.21E-05 | 2.53E-05 | 4.56E-05 |
| 0.000257 | 0.000354 | 0.000898 | 0.000683 | 0.000645 | 0.000179 | 0.000171 | 0.000179 | 0.000906 |
| 0.006107 | 0.005911 | 0.010593 | 0.010125 | 0.007652 | 0.003256 | 0.003772 | 0.003419 | 0.012077 |
| 0.009798 | 0.009157 | 0.015756 | 0.014941 | 0.011619 | 0.005912 | 0.006594 | 0.010853 | 0.015906 |
| 0.003721 | 0.004234 | 0.01103 | 0.009357 | 0.007276 | 0.001261 | 0.001593 | 0.00102 | 0.010856 |
| 0.000912 | 0.000979 | 0.00176 | 0.001613 | 0.001289 | 0.000557 | 0.000599 | 0.000736 | 0.002259 |
| 0.014202 | 0.014968 | 0.025394 | 0.025491 | 0.020199 | 0.008161 | 0.009178 | 0.006735 | 0.03315 |
| 0.016236 | 0.019605 | 0.023823 | 0.026368 | 0.021063 | 0.011324 | 0.011252 | 0.007574 | 0.03623 |
| 0.000789 | 0.000693 | 0.001368 | 0.001421 | 0.000839 | 0.000283 | 0.000363 | 0.000398 | 0.001294 |
| 0.031987 | 0.031619 | 0.056373 | 0.062418 | 0.038612 | 0.01163 | 0.017703 | 0.011568 | 0.074085 |
| 0.023094 | 0.026451 | 0.031142 | 0.03772 | 0.026908 | 0.014539 | 0.017781 | 0.010681 | 0.052498 |
| 0.004597 | 0.004637 | 0.006029 | 0.006134 | 0.005073 | 0.0029 | 0.003509 | 0.0042 | 0.006058 |
| 0.006274 | 0.005903 | 0.007911 | 0.008823 | 0.00659 | 0.003816 | 0.004736 | 0.004808 | 0.008191 |
| 0.002497 | 0.002323 | 0.00256 | 0.002614 | 0.002183 | 0.001664 | 0.002461 | 0.002269 | 0.002361 |
| 0.03475 | 0.036243 | 0.056077 | 0.066007 | 0.043479 | 0.021793 | 0.023686 | 0.021788 | 0.079681 |
| 0.016991 | 0.014837 | 0.022379 | 0.025174 | 0.015268 | 0.008474 | 0.011426 | 0.011331 | 0.019904 |
| 0.083413 | 0.091366 | 0.126302 | 0.194065 | 0.097912 | 0.050865 | 0.050626 | 0.043458 | 0.23312 |
| 0.038228 | 0.038893 | 0.054447 | 0.058762 | 0.041481 | 0.022312 | 0.030094 | 0.025817 | 0.071022 |
| 0.081247 | 0.085431 | 0.128376 | 0.177204 | 0.094228 | 0.043675 | 0.050878 | 0.0396 | 0.228912 |
| 0.066743 | 0.074321 | 0.104449 | 0.134051 | 0.077153 | 0.0389 | 0.042251 | 0.034266 | 0.201308 |
| 0.050061 | 0.059374 | 0.075485 | 0.098326 | 0.059576 | 0.031392 | 0.034009 | 0.027153 | 0.162005 |
| 0.02757 | 0.031012 | 0.04618 | 0.056066 | 0.036515 | 0.019803 | 0.017691 | 0.01884 | 0.082164 |
| 0.012279 | 0.011959 | 0.014853 | 0.021293 | 0.010776 | 0.007331 | 0.007398 | 0.010102 | 0.018178 |
| 0.055403 | 0.076598 | 0.082745 | 0.138809 | 0.071273 | 0.037769 | 0.034596 | 0.03143 | 0.202907 |
| 0.01455 | 0.019143 | 0.019149 | 0.025208 | 0.01582 | 0.011547 | 0.011696 | 0.011721 | 0.040522 |

| | | | | | | | | |
|---|---|---|---|---|---|---|---|---|
| 0.033916 | 0.042302 | 0.046402 | 0.060991 | 0.040086 | 0.025491 | 0.026101 | 0.022728 | 0.106208 |
| 0.001605 | 0.00158 | 0.002143 | 0.0025 | 0.001934 | 0.001506 | 0.001703 | 0.001345 | 0.004035 |
| 0.000524 | 0.000753 | 0.000592 | 0.000961 | 0.000616 | 0.000504 | 0.000338 | 0.000464 | 0.001477 |
| 0.000566 | 0.00047 | 0.00055 | 0.000746 | 0.000472 | 0.000319 | 0.000394 | 0.000327 | 0.000926 |
| 0.000969 | 0.002474 | 0.001316 | 0.001842 | 0.001626 | 0.000798 | 0.000445 | 0.000485 | 0.004213 |
| 0.013309 | 0.015037 | 0.013201 | 0.018947 | 0.013589 | 0.010664 | 0.010401 | 0.009394 | 0.022849 |
| 0.010724 | 0.012822 | 0.013267 | 0.015325 | 0.011782 | 0.009581 | 0.009322 | 0.006859 | 0.025803 |
| 0.729238 | 0.823485 | 1.07444 | 1.412171 | 0.856855 | 0.462645 | 0.492807 | 0.427006 | 1.905839 |
| 0.007396 | 0.013215 | 0.007278 | 0.010926 | 0.008509 | 0.006189 | 0.004737 | 0.004352 | 0.0212 |
| 0.005779 | 0.006806 | 0.006491 | 0.009774 | 0.006417 | 0.002799 | 0.003142 | 0.002376 | 0.009493 |
| 0.000393 | 0.000544 | 0.000393 | 0.000572 | 0.000353 | 0.000262 | 0.000322 | 0.000273 | 0.000957 |
| 0.00505 | 0.007308 | 0.005462 | 0.007661 | 0.005509 | 0.003878 | 0.003489 | 0.00316 | 0.014905 |
| 0.009614 | 0.009675 | 0.010156 | 0.014557 | 0.009834 | 0.007614 | 0.007829 | 0.006705 | 0.01838 |
| 0.000138 | 0.000163 | 0.000123 | 0.000179 | 0.000129 | 0.000133 | 0.000104 | 0.000147 | 0.000301 |
| 0.001214 | 0.001814 | 0.002376 | 0.002324 | 0.002248 | 0.001158 | 0.000988 | 0.000628 | 0.003699 |
| 0.000534 | 0.000493 | 0.000646 | 0.000656 | 0.00047 | 0.000293 | 0.000497 | 0.0003 | 0.000891 |
| 0.008085 | 0.00942 | 0.010601 | 0.011394 | 0.009649 | 0.00646 | 0.007727 | 0.00456 | 0.018793 |
| 0.001578 | 0.001591 | 0.001647 | 0.002088 | 0.001689 | 0.00133 | 0.001316 | 0.001074 | 0.002701 |
| 5.42E-05 | 4.53E-05 | 7.26E-05 | 8.35E-05 | 5.04E-05 | 1.88E-05 | 3.43E-05 | 2.47E-05 | 0.00015 |
| 0.000126 | 9.83E-05 | 0.000119 | 0.000109 | 0.000109 | 8.2E-05 | 0.000142 | 8.41E-05 | 0.000147 |
| 0.00146 | 0.001723 | 0.001512 | 0.002929 | 0.001695 | 0.001812 | 0.00148 | 0.001902 | 0.003814 |
| 0.002247 | 0.002705 | 0.002134 | 0.002628 | 0.002334 | 0.002659 | 0.002077 | 0.001893 | 0.003459 |
| 0.022235 | 0.026402 | 0.022757 | 0.042177 | 0.024265 | 0.019732 | 0.016144 | 0.017954 | 0.045799 |

| 33 (Control | 4 (JAK2A - | 5 (JAK2A - | 10 (JAK2A | 11 (JAK2A | 15 (JAK2A | 25 (JAK2A | 20 (JAK2A | 26 (JAK2A |
|---|---|---|---|---|---|---|---|---|
| CON-GH-In | JAK2A-Veh | JAK2A-Veh | JAK2A-Veh | JAK2A-Veh | JAK2A-Veh | JAK2A-Veh | JAK2A-Veh | JAK2A-Veh |
| 27.9004 | 24.0338 | 31.11558 | 49.20425 | 18.02262 | 26.24863 | 16.55685 | 16.84811 | 22.89146 |
| 9.83317 | 8.495131 | 12.27368 | 12.56812 | 6.534897 | 10.51258 | 4.905394 | 5.819121 | 8.869746 |
| 557.8479 | 509.9396 | 552.7329 | 600.425 | 488.496 | 503.58 | 432.551 | 440.4934 | 542.9099 |
| 454.602 | 449.8189 | 442.5203 | 511.5939 | 428.4709 | 433.6274 | 348.8484 | 397.7487 | 433.0254 |
| 40.25789 | 51.79074 | 61.85543 | 83.0366 | 33.64726 | 56.14583 | 33.41545 | 41.64842 | 64.79446 |
| 1.122739 | 0.896596 | 1.220131 | 1.490437 | 0.630113 | 0.988261 | 0.602187 | 0.57983 | 0.964711 |
| 1091.564 | 1044.975 | 1101.718 | 1258.318 | 975.8018 | 1031.103 | 836.8793 | 903.1376 | 1073.456 |
| 103.1488 | 74.07386 | 69.43723 | 43.73872 | 78.99367 | 119.4837 | 59.54343 | 90.02755 | 56.75799 |
| 90.40835 | 130.4402 | 71.80375 | 45.962 | 87.63079 | 136.0297 | 64.24695 | 131.7355 | 60.68493 |
| 96.28857 | 63.59823 | 43.59596 | 37.74014 | 46.34655 | 54.23587 | 46.38909 | 57.74105 | 34.14155 |
| 705.1361 | 370.6883 | 322.8283 | 226.0048 | 355.5015 | 470.4009 | 298.4925 | 357.686 | 248.3105 |
| 580.4265 | 322.9365 | 300.0289 | 211.7672 | 343.9853 | 437.0976 | 277.2491 | 311.1295 | 231.8265 |
| 46.11888 | 70.72378 | 42.79942 | 35.23896 | 52.77041 | 56.59456 | 43.7117 | 64.29752 | 42.6621 |
| 137.2051 | 129.8552 | 123.8672 | 133.2684 | 122.3592 | 130.6083 | 113.0395 | 127.6033 | 121.7352 |
| 87.95826 | 114.5938 | 93.73737 | 107.658 | 104.3652 | 90.68649 | 100.1694 | 111.5152 | 112.0548 |
| 1846.691 | 1276.91 | 1068.098 | 841.3781 | 1191.953 | 1495.137 | 1002.842 | 1251.736 | 908.1735 |
| 0.072322 | 0.160481 | 0.117529 | 0.025957 | 0.121482 | 0.095116 | 0.097396 | 0.084927 | 0.049866 |
| 5.68046 | 8.532073 | 5.501279 | 2.773103 | 8.500134 | 5.877826 | 5.461735 | 5.141904 | 4.449753 |
| 41.1954 | 63.00608 | 36.52941 | 23.70679 | 57.94126 | 43.96003 | 43.09348 | 45.58962 | 30.0212 |
| 26.64368 | 38.79541 | 22.89821 | 14.98242 | 40.06142 | 30.35603 | 25.32347 | 25.98764 | 19.01004 |
| 0.483908 | 0.698366 | 0.56624 | 0.54767 | 0.677944 | 0.705159 | 0.500659 | 0.494611 | 0.60882 |
| 0.869195 | 1.948521 | 0.947417 | 0.543356 | 1.937463 | 1.6399 | 0.950235 | 1.11204 | 0.624594 |
| 0.146115 | 0.557866 | 0.150744 | 0.074157 | 0.46478 | 0.340887 | 0.191258 | 0.274116 | 0.087901 |
| 0.341379 | 0.67819 | 0.530563 | 0.254541 | 0.542884 | 0.504585 | 0.439408 | 0.500396 | 0.213888 |
| 2.806897 | 7.003106 | 2.499744 | 1.411957 | 5.909479 | 4.733292 | 2.972966 | 4.970581 | 1.447378 |
| 1.728506 | 3.950034 | 1.649923 | 0.931398 | 3.962884 | 2.729794 | 1.912818 | 2.91471 | 1.117908 |
| 3.767816 | 5.62971 | 3.719693 | 2.777896 | 5.226969 | 3.433604 | 3.469267 | 3.524351 | 2.729894 |
| 1.874483 | 3.755571 | 2.884143 | 2.217284 | 2.927103 | 2.2007 | 3.110962 | 2.322868 | 2.271435 |
| 10.93793 | 24.67252 | 12.46419 | 8.352863 | 17.52657 | 11.92655 | 15.09966 | 13.65241 | 8.067562 |
| 67.58621 | 82.62255 | 64.56522 | 46.28229 | 77.14286 | 70.51593 | 72.96839 | 71.04326 | 51.49399 |
| 45.81609 | 73.92032 | 44.76982 | 24.51931 | 67.19359 | 52.59463 | 52.46268 | 57.8937 | 29.38516 |
| 13.27586 | 25.88791 | 16.42788 | 15.01358 | 21.58318 | 17.7684 | 17.08971 | 17.06996 | 13.16099 |
| 16.01609 | 32.11074 | 17.34399 | 8.496671 | 30.83311 | 20.14516 | 18.55683 | 25.3424 | 12.61145 |
| 8.418391 | 11.63133 | 8.245013 | 12.10866 | 13.33298 | 11.90631 | 11.08568 | 7.366873 | 9.962968 |
| 1.092644 | 2.406968 | 0.967442 | 0.409134 | 2.131883 | 1.631805 | 1.090652 | 1.713226 | 0.696848 |
| 0.481379 | 1.303876 | 0.721841 | 0.717124 | 0.911055 | 0.717976 | 0.673033 | 0.735575 | 0.538092 |
| 1.052874 | 3.444429 | 1.551637 | 1.096538 | 2.72283 | 1.442698 | 2.101654 | 1.958863 | 1.074657 |
| 0.200368 | 0.610614 | 0.327545 | 0.296724 | 0.485928 | 0.22034 | 0.348379 | 0.339753 | 0.216483 |
| 0.064023 | 0.119084 | 0.066476 | 0.046977 | 0.120232 | 0.082344 | 0.051204 | 0.086062 | 0.05643 |
| 0.726667 | 2.596084 | 1.286701 | 0.961598 | 1.955728 | 0.996127 | 1.570733 | 1.43555 | 0.894785 |
| 0.078253 | 0.221883 | 0.124642 | 0.203944 | 0.127658 | 0.070741 | 0.103618 | 0.081456 | 0.097086 |
| 3.126437 | 7.632681 | 3.257033 | 1.498961 | 6.89239 | 5.396627 | 3.84694 | 5.484549 | 1.632339 |
| 1.80069 | 4.056989 | 1.780435 | 0.923249 | 4.21522 | 3.085072 | 2.034351 | 3.26403 | 0.875194 |
| 0.033195 | 0.119109 | 0.053425 | 0.035856 | 0.137583 | 0.114813 | 0.059944 | 0.082635 | 0.020994 |
| 0.047425 | 0.092783 | 0.028496 | 0.023489 | 0.105356 | 0.096262 | 0.023726 | 0.027545 | 0.027553 |
| 0.334253 | 1.060554 | 0.445166 | 0.27755 | 0.930761 | 0.662211 | 0.509375 | 0.681508 | 0.272989 |
| 2.790805 | 4.650101 | 2.999233 | 2.203862 | 3.765821 | 2.86471 | 2.682448 | 2.84796 | 2.174247 |

| | | | | | | | | |
|---|---|---|---|---|---|---|---|---|
| 37.35632 | 64.68332 | 35.47059 | 19.37816 | 57.84513 | 45.82636 | 45.00605 | 49.79481 | 24.72424 |
| 0.126851 | 0.209364 | 0.167386 | 0.148338 | 0.167335 | 0.183238 | 0.183535 | 0.135412 | 0.171884 |
| 4.802299 | 11.19379 | 5.961637 | 5.241811 | 8.634713 | 6.606371 | 6.553598 | 6.543634 | 4.177527 |
| 0.476782 | 0.603079 | 0.452302 | 0.489667 | 0.648385 | 0.662211 | 0.555615 | 0.498171 | 0.564297 |
| 0.062828 | 0.087752 | 0.087836 | 0.093955 | 0.102881 | 0.112497 | 0.079917 | 0.077051 | 0.077419 |
| 0.585747 | 0.751114 | 0.584194 | 0.438855 | 0.729373 | 0.778239 | 0.53189 | 0.541557 | 0.465074 |
| 1.739081 | 4.951519 | 3.153453 | 2.487883 | 3.943658 | 2.365522 | 2.303564 | 2.712237 | 1.928735 |
| 0.403678 | 0.930263 | 0.268849 | 0.196562 | 0.878852 | 0.497614 | 0.313517 | 0.442991 | 0.268919 |
| 4.586207 | 17.21729 | 7.232225 | 5.574967 | 13.97944 | 5.814866 | 10.34728 | 9.046724 | 5.541201 |
| 331.0898 | 563.703 | 333.706 | 229.6854 | 511.6763 | 391.8316 | 387.3447 | 404.7838 | 255.0799 |
| 0.073494 | 0.134131 | 0.073588 | 0.059321 | 0.133858 | 0.112475 | 0.078658 | 0.07803 | 0.051341 |
| 0.047425 | 0.12414 | 0.078353 | 0.060567 | 0.09173 | 0.061477 | 0.092409 | 0.072289 | 0.102352 |
| 7.135632 | 9.424173 | 6.764962 | 9.973103 | 12.10254 | 10.02873 | 9.257835 | 5.958467 | 8.986007 |
| 0.061655 | 0.147962 | 0.056992 | 0.089017 | 0.104131 | 0.069594 | 0.069918 | 0.074603 | 0.065614 |
| 0.716092 | 1.075625 | 1.027519 | 0.85494 | 1.235728 | 1.10406 | 0.885353 | 0.771174 | 0.714403 |
| 0.067586 | 0.18051 | 0.090483 | 0.060567 | 0.15616 | 0.090461 | 0.118604 | 0.084905 | 0.062892 |
| 0.457701 | 0.941931 | 0.414322 | 0.252144 | 1.094179 | 0.830181 | 0.50308 | 0.859506 | 0.276806 |
| 9.183908 | 29.19379 | 12.92455 | 8.616511 | 21.7466 | 11.98051 | 16.9033 | 17.22571 | 8.176961 |
| 0.164805 | 0.393545 | 0.166189 | 0.196538 | 0.333324 | 0.306259 | 0.244035 | 0.336193 | 0.234878 |
| 2.09954 | 4.640378 | 2.016829 | 0.954168 | 4.320961 | 3.307683 | 2.182999 | 3.455377 | 1.46824 |
| 0.48023 | 1.13688 | 0.409488 | 0.384447 | 1.068224 | 0.714379 | 0.456839 | 0.604079 | 0.373993 |
| 0.972184 | 1.806563 | 0.883197 | 0.418961 | 1.970628 | 1.563223 | 0.794082 | 1.445785 | 0.754601 |
| 115.4617 | 108.9632 | 131.0642 | 129.5623 | 127.6186 | 118.3752 | 148.2153 | 140.5196 | 120.9868 |
| 5.320432 | 10.04053 | 11.50275 | 8.054615 | 10.13304 | 10.19746 | 7.35747 | 12.01708 | 6.854884 |
| 0.35558 | 0.434842 | 0.497798 | 0.435528 | 0.288624 | 0.441749 | 0.464899 | 0.365552 | 0.420027 |
| 424.9455 | 464.2713 | 544.5214 | 537.9999 | 491.219 | 450.5165 | 553.0544 | 548.6289 | 474.8988 |
| 0.485187 | 0.808579 | 0.942569 | 0.828582 | 0.750423 | 0.689535 | 0.918412 | 0.498363 | 0.782481 |
| 5.486641 | 2.748947 | 3.581651 | 4.663559 | 4.025135 | 3.870804 | 2.786003 | 3.606406 | 3.0995 |
| 2.498428 | 3.999474 | 3.702752 | 3.680495 | 2.190033 | 3.171791 | 4.738627 | 4.381495 | 4.355657 |
| 1.301198 | 2.325789 | 2.766972 | 2.711132 | 1.697616 | 1.760226 | 2.178896 | 2.671174 | 1.823716 |
| 1.457328 | 3.031579 | 3.598165 | 4.005899 | 2.418202 | 2.300141 | 3.100942 | 3.384342 | 3.212358 |
| 10.7222 | 17.21053 | 18.86239 | 21.49382 | 14.06652 | 11.17236 | 16.19758 | 15.93523 | 13.48078 |
| 2.636346 | 7.45579 | 5.719266 | 5.420552 | 4.489274 | 4.353174 | 4.690175 | 8.237722 | 6.401817 |
| 86.92338 | 84.63158 | 98.49541 | 97.45005 | 88.94691 | 83.25529 | 97.34051 | 93.20285 | 90.48251 |
| 86.85265 | 92.46316 | 117.9817 | 111.7507 | 103.0856 | 93.93512 | 119.5559 | 113.5089 | 96.04362 |
| 0.054707 | 0.057821 | 0.08989 | 0.050335 | 0.053298 | 0.051504 | 0.06621 | 0.08368 | 0.074224 |
| 49.40275 | 57.80526 | 73.72477 | 75.16841 | 58.38787 | 58.00282 | 64.02961 | 74.04982 | 58.80055 |
| 0.130403 | 0.199579 | 0.156239 | 0.17589 | 0.131733 | 0.158014 | 0.183731 | 0.344626 | 0.119826 |
| 55.85658 | 72.09474 | 71.83486 | 72.54805 | 72.93608 | 58.78138 | 81.23015 | 75.82206 | 67.96002 |
| 53.53534 | 55.9408 | 57.40511 | 52.79934 | 58.55422 | 58.45995 | 55.50487 | 57.08241 | 51.58471 |
| 44.29748 | 46.01389 | 47.00365 | 42.47269 | 51.67153 | 51.4522 | 46.60325 | 48.9398 | 39.32773 |
| 56.12415 | 55.40803 | 56.74818 | 53.9753 | 57.85796 | 53.00258 | 52.48449 | 52.9585 | 50.63811 |
| 59.47634 | 61.67079 | 59.40876 | 56.4952 | 59.69562 | 56.84755 | 59.20591 | 59.0076 | 55.45893 |
| 80.47449 | 80.99185 | 81.68978 | 80.18227 | 80.29803 | 72.45478 | 75.76485 | 74.8194 | 75.48387 |
| 84.11432 | 83.98188 | 86.34307 | 81.41751 | 85.16043 | 80.23773 | 82.41182 | 79.36412 | 80.64625 |
| 49.88445 | 52.61371 | 50.62774 | 48.9849 | 52.74445 | 47.50388 | 49.86824 | 49.66569 | 46.90052 |
| 66.25814 | 71.32588 | 68.67153 | 68.69942 | 70.67597 | 64.50646 | 65.9805 | 68.58095 | 63.65628 |
| 62.67363 | 59.20266 | 61.27007 | 56.13945 | 62.80025 | 55.90698 | 59.36544 | 57.21917 | 54.7563 |
| 67.15427 | 67.41166 | 69.78832 | 64.63794 | 70.08244 | 65.54005 | 64.89572 | 65.5827 | 62.46571 |

| | | | | | | | | |
|---|---|---|---|---|---|---|---|---|
| 134.3085 | 122.2108 | 136.9708 | 128.2679 | 131.6043 | 126.8217 | 129.0044 | 123.4015 | 125.595 |
| 42.18439 | 41.27333 | 42.99635 | 39.37963 | 45.31389 | 44.77519 | 40.72201 | 43.76388 | 40.34264 |
| 11.40627 | 13.254 | 15.03285 | 13.96322 | 13.25174 | 12.1447 | 13.07061 | 13.97078 | 14.238 |
| 80.76214 | 80.89399 | 82.9708 | 75.58715 | 84.16741 | 79.21447 | 81.1675 | 79.62712 | 75.41556 |
| 77.92993 | 80.39384 | 77.35402 | 71.82213 | 84.3272 | 74.80103 | 76.50931 | 76.09234 | 71.41448 |
| 38.45605 | 40.42525 | 34.08394 | 33.54927 | 34.64173 | 33.70543 | 36.91462 | 35.737 | 35.11195 |
| 35.31408 | 40.38176 | 44.04745 | 46.66264 | 41.19341 | 40.2584 | 40.24343 | 40.18703 | 40.1865 |
| 78.01844 | 89.13561 | 85.86131 | 77.48449 | 89.36081 | 91.05943 | 82.6777 | 86.22326 | 79.87531 |
| 10.81217 | 12.71036 | 14.61679 | 14.32885 | 12.4071 | 11.31783 | 12.23043 | 12.37171 | 14.04283 |
| 12.09219 | 17.33132 | 16.4781 | 18.31128 | 15.52315 | 14.45995 | 16.63338 | 18.79953 | 16.40445 |
| 23.62016 | 28.99789 | 26.09124 | 27.21493 | 26.54914 | 23.54522 | 28.74682 | 29.60374 | 24.30903 |
| 59.22188 | 59.36575 | 59.96715 | 55.8924 | 63.15409 | 56.83721 | 55.44106 | 56.98773 | 55.49797 |
| 73.72588 | 75.47931 | 75.45985 | 69.53939 | 76.31452 | 73.05426 | 71.86174 | 72.76797 | 70.35077 |
| 76.22618 | 74.23981 | 70.92701 | 64.43041 | 80.40076 | 68.9199 | 77.59409 | 78.22794 | 66.15451 |
| 9.055317 | 10.19764 | 9.788321 | 10.98875 | 10.25098 | 11.24548 | 10.50328 | 10.25821 | 8.50084 |
| 42.33927 | 48.7756 | 43.42336 | 43.14466 | 46.30691 | 40.04134 | 46.11403 | 46.85681 | 40.33288 |
| 66.9662 | 75.29447 | 67.34672 | 66.8515 | 70.80152 | 62.71835 | 68.51167 | 66.17183 | 61.54839 |
| 1496.432 | 1544.922 | 1542.372 | 1463.223 | 1575.11 | 1470.832 | 1500.031 | 1504.269 | 1420.24 |
| 33.55501 | 36.22833 | 34.56569 | 36.39528 | 34.68738 | 35.91731 | 35.61713 | 34.60082 | 32.99431 |
| 12.19176 | 11.56871 | 10.69161 | 10.30689 | 11.93912 | 10.97674 | 9.386588 | 9.824781 | 9.311792 |
| 0.803971 | 0.820356 | 0.955401 | 1.187812 | 0.792365 | 0.867287 | 0.972479 | 0.64678 | 0.880239 |
| 70.74985 | 69.8798 | 68.60584 | 65.46802 | 73.94039 | 64.48579 | 71.14919 | 71.38983 | 61.58742 |
| 147.6804 | 152.0182 | 148.3441 | 140.6322 | 156.001 | 148.7743 | 148.297 | 151.0315 | 129.7269 |
| 30.37984 | 33.52099 | 33.52555 | 27.27422 | 34.64172 | 36.52713 | 31.17164 | 34.56926 | 24.9531 |
| 8.18E-05 | 7.7E-05 | 0.000144 | 2.08E-05 | 4.27E-05 | 3.96E-05 | 4.3E-05 | 1.69E-05 | 2.86E-05 |
| 0.000554 | 0.000427 | 0.000926 | 0.000129 | 0.000307 | 0.000477 | 0.000405 | 0.000145 | 0.000187 |
| 0.007767 | 0.00548 | 0.006918 | 0.001992 | 0.004546 | 0.005304 | 0.00584 | 0.003605 | 0.002582 |
| 0.013512 | 0.010031 | 0.014585 | 0.005106 | 0.009187 | 0.008035 | 0.011399 | 0.006927 | 0.007134 |
| 0.008245 | 0.010175 | 0.008629 | 0.000979 | 0.004509 | 0.007155 | 0.008174 | 0.003086 | 0.002147 |
| 0.00128 | 0.000972 | 0.001466 | 0.000391 | 0.000794 | 0.000955 | 0.001295 | 0.00054 | 0.000524 |
| 0.020609 | 0.014993 | 0.01686 | 0.004597 | 0.012781 | 0.01557 | 0.016716 | 0.01001 | 0.006961 |
| 0.024165 | 0.01508 | 0.015057 | 0.005489 | 0.015418 | 0.019394 | 0.014907 | 0.012496 | 0.008379 |
| 0.00088 | 0.000653 | 0.000841 | 0.000282 | 0.0006 | 0.000644 | 0.000766 | 0.000463 | 0.00035 |
| 0.040115 | 0.025048 | 0.02828 | 0.009938 | 0.022414 | 0.029157 | 0.028367 | 0.019684 | 0.013378 |
| 0.032582 | 0.019139 | 0.019191 | 0.009569 | 0.021655 | 0.025629 | 0.019346 | 0.019269 | 0.011988 |
| 0.005508 | 0.003471 | 0.005546 | 0.00271 | 0.003473 | 0.003685 | 0.004436 | 0.003908 | 0.003272 |
| 0.007523 | 0.004201 | 0.005381 | 0.003264 | 0.00434 | 0.004447 | 0.004511 | 0.005015 | 0.003405 |
| 0.002425 | 0.002297 | 0.002976 | 0.002123 | 0.001912 | 0.00189 | 0.002754 | 0.002981 | 0.00209 |
| 0.051752 | 0.027752 | 0.029482 | 0.015635 | 0.028332 | 0.03275 | 0.026944 | 0.023095 | 0.018977 |
| 0.01674 | 0.010827 | 0.012868 | 0.007516 | 0.010168 | 0.009664 | 0.012221 | 0.011746 | 0.008328 |
| 0.134187 | 0.056118 | 0.048097 | 0.035322 | 0.060863 | 0.076144 | 0.045812 | 0.046221 | 0.034495 |
| 0.04625 | 0.028178 | 0.033395 | 0.020173 | 0.026466 | 0.02915 | 0.031666 | 0.027827 | 0.021599 |
| 0.123089 | 0.052261 | 0.051936 | 0.031825 | 0.052931 | 0.067603 | 0.050883 | 0.045372 | 0.03473 |
| 0.10277 | 0.042512 | 0.041749 | 0.02628 | 0.044522 | 0.057972 | 0.042041 | 0.03886 | 0.029726 |
| 0.08034 | 0.034006 | 0.032874 | 0.022531 | 0.036927 | 0.047401 | 0.032763 | 0.031871 | 0.024301 |
| 0.046579 | 0.021391 | 0.023938 | 0.011879 | 0.024013 | 0.029755 | 0.021241 | 0.017588 | 0.014863 |
| 0.014169 | 0.007684 | 0.007636 | 0.004786 | 0.008531 | 0.007516 | 0.007561 | 0.007151 | 0.005487 |
| 0.113555 | 0.037833 | 0.032531 | 0.026403 | 0.04879 | 0.063977 | 0.031101 | 0.03468 | 0.027579 |
| 0.021421 | 0.010953 | 0.012481 | 0.008651 | 0.011481 | 0.014322 | 0.011332 | 0.010754 | 0.009819 |

| | | | | | | | | |
|---|---|---|---|---|---|---|---|---|
| 0.05666 | 0.027074 | 0.025852 | 0.020197 | 0.02997 | 0.037205 | 0.025055 | 0.025985 | 0.022558 |
| 0.002567 | 0.002027 | 0.001893 | 0.002003 | 0.001767 | 0.002165 | 0.001472 | 0.002298 | 0.00167 |
| 0.000759 | 0.00057 | 0.00038 | 0.000399 | 0.000608 | 0.000823 | 0.000348 | 0.000454 | 0.00037 |
| 0.000613 | 0.000422 | 0.000441 | 0.00031 | 0.000422 | 0.000557 | 0.000412 | 0.000473 | 0.000351 |
| 0.002168 | 0.001053 | 0.000521 | 0.00039 | 0.001572 | 0.002441 | 0.0004 | 0.000669 | 0.000424 |
| 0.017842 | 0.010677 | 0.007642 | 0.006335 | 0.011538 | 0.012404 | 0.007297 | 0.009507 | 0.007745 |
| 0.016037 | 0.010507 | 0.007973 | 0.010306 | 0.00941 | 0.012936 | 0.007358 | 0.009761 | 0.007902 |
| 1.109143 | 0.553434 | 0.538972 | 0.340791 | 0.571911 | 0.698225 | 0.510159 | 0.480678 | 0.373739 |
| 0.013497 | 0.006599 | 0.00423 | 0.003843 | 0.007919 | 0.010696 | 0.003187 | 0.004548 | 0.004591 |
| 0.007846 | 0.00436 | 0.002213 | 0.00213 | 0.005508 | 0.006251 | 0.002038 | 0.003964 | 0.001858 |
| 0.00062 | 0.000329 | 0.00028 | 0.000261 | 0.00041 | 0.000513 | 0.000287 | 0.000371 | 0.000273 |
| 0.006162 | 0.00464 | 0.002782 | 0.003413 | 0.00389 | 0.005397 | 0.002399 | 0.003662 | 0.003212 |
| 0.011223 | 0.008299 | 0.005089 | 0.00478 | 0.007053 | 0.008009 | 0.005169 | 0.006716 | 0.005343 |
| 0.00015 | 0.00012 | 9.9E-05 | 0.000116 | 0.000132 | 0.000147 | 8.66E-05 | 0.000125 | 0.000102 |
| 0.002656 | 0.001824 | 0.001241 | 0.001018 | 0.001793 | 0.002334 | 0.000801 | 0.001172 | 0.000754 |
| 0.000586 | 0.000432 | 0.000406 | 0.00029 | 0.000392 | 0.000513 | 0.000443 | 0.000513 | 0.000335 |
| 0.013348 | 0.008381 | 0.006814 | 0.007715 | 0.008225 | 0.009441 | 0.005051 | 0.006847 | 0.006283 |
| 0.002259 | 0.00156 | 0.001226 | 0.001699 | 0.001581 | 0.001513 | 0.001026 | 0.001184 | 0.001198 |
| 7.29E-05 | 4.67E-05 | 3.92E-05 | 1.93E-05 | 5.85E-05 | 6.94E-05 | 3.1E-05 | 4.06E-05 | 3.07E-05 |
| 0.000108 | 0.000142 | 0.000144 | 0.000129 | 9.66E-05 | 0.000105 | 0.000146 | 0.000147 | 0.000133 |
| 0.002249 | 0.001844 | 0.001122 | 0.001751 | 0.001477 | 0.001602 | 0.00102 | 0.001359 | 0.001387 |
| 0.003032 | 0.001997 | 0.002133 | 0.002099 | 0.001958 | 0.002744 | 0.001654 | 0.002197 | 0.001815 |
| 0.032589 | 0.01897 | 0.012665 | 0.013997 | 0.021127 | 0.021726 | 0.011951 | 0.015374 | 0.013075 |

| 31 (JAK2A | 35 (JAK2A | 8 (JAK2A - | 1 (JAK2A - | 9 (JAK2A - | 14 (JAK2A | 18 (JAK2A | 27 (JAK2A | 22 (JAK2A |
|---|---|---|---|---|---|---|---|---|
| JAK2A-Veh | JAK2A-Veh | JAK2A-GH | JAK2A-GH | JAK2A-GH | JAK2A-GH | JAK2A-GH | JAK2A-GH | JAK2A-GH |
| 15.69296 | 21.8812 | 28.56765 | 24.83271 | 28.4713 | 30.04088 | 26.56941 | 23.21404 | 25.96502 |
| 5.656056 | 7.765338 | 9.327527 | 7.730855 | 8.605565 | 8.893948 | 9.232941 | 6.072983 | 7.118273 |
| 408.5493 | 528.5761 | 497.5894 | 524.3123 | 596.9739 | 509.9158 | 543.6706 | 512.8421 | 479.1068 |
| 357.2113 | 452.5631 | 472.9549 | 406.8401 | 476.2957 | 473.4108 | 455.4353 | 410.6667 | 415.735 |
| 29.04085 | 51.78845 | 42.17263 | 41.27881 | 70.8887 | 44.0513 | 55.10118 | 51.50877 | 34.60662 |
| 0.640648 | 0.982018 | 1.039549 | 0.774572 | 0.920348 | 0.796329 | 0.669882 | 1.006175 | 0.870726 |
| 816.7911 | 1063.556 | 1051.652 | 1005.769 | 1182.155 | 1067.109 | 1090.679 | 1005.311 | 963.4025 |
| 68.28528 | 52.9193 | 46.83359 | 48.01619 | 45.9314 | 61.00631 | 53.93802 | 46.50784 | 62.85284 |
| 125.22 | 71.84908 | 63.20995 | 68.3864 | 40.26086 | 72.01932 | 52.77521 | 69.71589 | 98.8793 |
| 56.15782 | 36.63602 | 26.23017 | 36 | 27.04902 | 44.40467 | 39.41761 | 25.57472 | 50.44498 |
| 419.6055 | 306.9082 | 267.3406 | 257.0804 | 226.5568 | 260.6414 | 304.1613 | 230.1542 | 401.4698 |
| 417.7238 | 286.6417 | 269.0202 | 237.8467 | 228.9416 | 241.8999 | 279.4884 | 214.835 | 332.0228 |
| 49.35357 | 30.29735 | 28.14495 | 25.07178 | 33.42441 | 42.12961 | 37.85342 | 32.21149 | 45.41834 |
| 116.5402 | 105.5368 | 148.0871 | 115.4992 | 143.5124 | 151.1391 | 168.6375 | 132.7354 | 169.2523 |
| 91.59332 | 78.26321 | 89.58009 | 89.46573 | 110.1251 | 99.35865 | 111.3232 | 110.0777 | 95.37388 |
| 1344.48 | 969.0517 | 938.4467 | 877.3664 | 855.8016 | 972.599 | 1047.595 | 861.8122 | 1255.714 |
| 0.11333 | 0.06239 | 0.125199 | 0.160392 | 0.030286 | 0.076062 | 0.038512 | 0.031961 | 0.141698 |
| 5.879221 | 4.378497 | 5.500717 | 5.942899 | 3.286073 | 4.446206 | 3.663356 | 3.666896 | 7.568895 |
| 50.02597 | 29.75443 | 45.78766 | 50.12592 | 25.94981 | 33.58227 | 22.61144 | 28.02202 | 59.99128 |
| 36.18701 | 18.83348 | 27.96844 | 28.72621 | 14.35257 | 19.42163 | 12.44727 | 16.30778 | 28.30814 |
| 0.659455 | 0.426414 | 0.443673 | 0.596662 | 0.367001 | 0.591135 | 0.552106 | 0.451177 | 0.588401 |
| 1.773584 | 0.898351 | 1.033773 | 1.368843 | 0.51606 | 0.937837 | 0.641936 | 0.645423 | 1.542297 |
| 0.39787 | 0.20186 | 0.190433 | 0.278038 | 0.116469 | 0.165832 | 0.103859 | 0.100939 | 0.452093 |
| 0.395766 | 0.30964 | 0.550846 | 0.613001 | 0.299247 | 0.382968 | 0.296417 | 0.28245 | 0.713721 |
| 4.962857 | 2.526817 | 3.460545 | 4.493411 | 1.486525 | 2.458831 | 1.70225 | 1.68702 | 6.030523 |
| 3.81974 | 1.628467 | 2.498565 | 2.634641 | 0.85325 | 1.763009 | 0.760729 | 1.009388 | 2.870058 |
| 3.892208 | 2.529017 | 4.441894 | 5.357833 | 2.660477 | 3.396911 | 2.391703 | 2.792292 | 5.206395 |
| 2.257247 | 1.958998 | 3.150646 | 3.87145 | 2.122509 | 2.420148 | 1.725556 | 2.106731 | 3.411628 |
| 11.66961 | 7.644227 | 17.77532 | 21.61318 | 8.618319 | 11.59302 | 7.111754 | 8.394219 | 21.98198 |
| 72.14026 | 55.17654 | 78.81779 | 84.09663 | 46.68256 | 67.01948 | 41.61157 | 55.94494 | 83.40698 |
| 55.05195 | 33.6909 | 54.3099 | 60.61493 | 25.11418 | 35.56481 | 17.43206 | 31.34205 | 60.56686 |
| 17.91351 | 12.77043 | 21.04993 | 21.88463 | 11.77792 | 15.87723 | 12.99711 | 13.68149 | 22.26977 |
| 28.35584 | 11.52572 | 20.17963 | 21.03865 | 7.267754 | 13.27092 | 5.315148 | 11.30048 | 18.79797 |
| 10.47273 | 8.92413 | 12.72138 | 10.40469 | 7.52522 | 11.90007 | 14.03796 | 9.276256 | 9.536337 |
| 1.902623 | 0.897251 | 1.32792 | 1.371479 | 0.535257 | 0.832908 | 0.367693 | 0.646662 | 1.264448 |
| 0.718597 | 0.478754 | 0.853773 | 0.912914 | 0.489184 | 0.67092 | 0.427203 | 0.5151 | 0.885349 |
| 1.790649 | 1.037996 | 2.016671 | 2.473353 | 1.051317 | 1.230866 | 0.521106 | 0.88947 | 2.190087 |
| 0.359299 | 0.150839 | 0.435409 | 0.357365 | 0.251593 | 0.26184 | 0.101529 | 0.237655 | 0.412849 |
| 0.115738 | 0.052164 | 0.075925 | 0.055713 | 0.016304 | 0.049878 | 0.017507 | 0.037065 | 0.078253 |
| 1.346494 | 0.679316 | 1.679397 | 1.767584 | 0.785044 | 0.922606 | 0.410685 | 0.673421 | 1.373547 |
| 0.075951 | 0.056716 | 0.151825 | 0.152223 | 0.126949 | 0.108459 | 0.056003 | 0.095835 | 0.124116 |
| 5.383636 | 2.85449 | 4.033859 | 5.357833 | 1.761154 | 3.065682 | 1.969931 | 1.953861 | 6.731686 |
| 3.896883 | 1.843983 | 2.469125 | 2.941142 | 1.035508 | 1.441209 | 0.854632 | 1.18307 | 2.943314 |
| 0.133831 | 0.057859 | 0.063943 | 0.053025 | 0.025634 | 0.061725 | 0.053695 | 0.024278 | 0.090419 |
| 0.090444 | 0.038573 | 0.047492 | 0.044855 | 0.023307 | 0.042407 | 0.021951 | 0.034315 | 0.055334 |
| 0.804156 | 0.506903 | 0.692626 | 0.819619 | 0.330866 | 0.423828 | 0.236003 | 0.254205 | 0.692791 |
| 3.165195 | 1.788564 | 3.563845 | 4.042753 | 2.006876 | 2.792478 | 1.860189 | 1.9883 | 3.906105 |

| | | | | | | | | |
|---|---|---|---|---|---|---|---|---|
| 51.19481 | 28.06109 | 46.27834 | 52.55051 | 20.38269 | 32.68771 | 15.31414 | 21.42161 | 52.61337 |
| 0.150709 | 0.107736 | 0.145162 | 0.212047 | 0.122296 | 0.168322 | 0.121373 | 0.118778 | 0.178116 |
| 5.245714 | 3.940867 | 7.16901 | 7.742899 | 4.008783 | 5.362525 | 3.559271 | 4.11287 | 7.806977 |
| 0.555896 | 0.332291 | 0.496872 | 0.391625 | 0.375132 | 0.503855 | 0.486713 | 0.388493 | 0.527703 |
| 0.059073 | 0.040838 | 0.051934 | 0.09817 | 0.065225 | 0.06743 | 0.066524 | 0.065187 | 0.068808 |
| 0.725844 | 0.562541 | 0.617991 | 0.664656 | 0.343513 | 0.518845 | 0.515902 | 0.472732 | 0.56407 |
| 2.73974 | 1.533244 | 3.509613 | 3.336457 | 1.928733 | 1.988583 | 0.873413 | 2.257619 | 2.90407 |
| 0.94161 | 0.380012 | 0.609986 | 0.449341 | 0.202675 | 0.293029 | 0.112028 | 0.181462 | 0.333314 |
| 9.605455 | 4.151985 | 9.906456 | 11.1347 | 5.054454 | 6.116857 | 1.927618 | 4.080661 | 8.997384 |
| 430.3559 | 262.4601 | 419.1828 | 456.336 | 217.3367 | 308.4298 | 193.7151 | 246.5641 | 460.666 |
| 0.096452 | 0.093002 | 0.114534 | 0.114141 | 0.052396 | 0.102246 | 0.045526 | 0.047273 | 0.109282 |
| 0.059073 | 0.065798 | 0.085248 | 0.118252 | 0.085031 | 0.076062 | 0.085192 | 0.06645 | 0.122808 |
| 8.467013 | 7.521075 | 10.61148 | 8.380674 | 5.630364 | 9.45816 | 12.38165 | 7.977977 | 8.879651 |
| 0.08079 | 0.056716 | 0.071923 | 0.061168 | 0.05592 | 0.076086 | 0.087545 | 0.043458 | 0.056669 |
| 1.021091 | 0.798509 | 0.99891 | 0.996457 | 0.626725 | 0.844271 | 0.855311 | 0.753944 | 0.879855 |
| 0.125369 | 0.076002 | 0.109214 | 0.084255 | 0.052419 | 0.107226 | 0.04668 | 0.085677 | 0.129532 |
| 1.098234 | 0.379352 | 0.558077 | 0.477013 | 0.23059 | 0.408838 | 0.170361 | 0.240206 | 0.587616 |
| 14.97273 | 7.439707 | 15.67059 | 20.61435 | 8.789962 | 9.794224 | 3.491389 | 6.481487 | 17.32762 |
| 0.330312 | 0.225632 | 0.290273 | 0.347877 | 0.166562 | 0.256763 | 0.164546 | 0.248011 | 0.33593 |
| 4.156364 | 1.738644 | 2.732281 | 2.783016 | 1.024893 | 1.711995 | 0.634695 | 1.195953 | 2.547471 |
| 1.050078 | 0.451265 | 0.632712 | 0.4894 | 0.210806 | 0.362901 | 0.169184 | 0.274769 | 0.619273 |
| 1.92787 | 0.820061 | 1.104017 | 1.11716 | 0.462309 | 0.750705 | 0.267228 | 0.472732 | 0.943169 |
| 150.6992 | 139.7682 | 148.2618 | 107.1031 | 139.1146 | 129.833 | 128.7087 | 126.2737 | 119.0415 |
| 9.596339 | 11.30979 | 9.091178 | 10.69624 | 9.044049 | 8.887265 | 5.81068 | 8.232617 | 9.258986 |
| 0.370323 | 0.443652 | 0.379465 | 0.312868 | 0.336574 | 0.393445 | 0.424485 | 0.425506 | 0.349862 |
| 513.1169 | 538.6677 | 569.6796 | 460.5636 | 536.0965 | 510.4915 | 468.1195 | 474.6142 | 499.8829 |
| 1.781989 | 1.414215 | 1.183472 | 0.586341 | 0.957264 | 0.859603 | 0.538951 | 0.603146 | 0.756866 |
| 4.039292 | 5.06047 | 3.759818 | 5.105423 | 4.593103 | 3.372651 | 2.319029 | 3.437651 | 3.010138 |
| 2.822453 | 4.302679 | 3.026295 | 2.692526 | 3.34772 | 3.568059 | 2.596893 | 2.990247 | 2.82765 |
| 1.589359 | 1.871843 | 1.541833 | 1.345911 | 1.807008 | 2.096868 | 1.656524 | 1.789428 | 1.939171 |
| 1.758267 | 2.537124 | 2.430051 | 2.314411 | 2.506785 | 2.222756 | 2.132039 | 2.227583 | 2.665438 |
| 13.82453 | 14.08704 | 14.38976 | 14.38241 | 16.49833 | 13.62589 | 11.89573 | 12.42538 | 15.78065 |
| 4.269921 | 4.403062 | 5.036312 | 3.784661 | 4.969522 | 5.03737 | 5.744272 | 3.930362 | 2.881106 |
| 98.687 | 98.25697 | 111.2578 | 86.5618 | 105.1568 | 102.7766 | 93.00583 | 92.55794 | 95.24424 |
| 110.46 | 115.9913 | 113.6756 | 101.5105 | 112.9655 | 104.7871 | 95.53981 | 109.1138 | 103.2811 |
| 0.067959 | 0.068004 | 0.06657 | 0.072563 | 0.059887 | 0.062981 | 0.036944 | 0.060371 | 0.053235 |
| 52.73704 | 64.48114 | 72.71713 | 59.79482 | 66.27364 | 61.66597 | 56.09709 | 62.2968 | 65.69585 |
| 0.164251 | 0.17258 | 0.166969 | 0.21403 | 0.0897 | 0.185035 | 0.115375 | 0.111209 | 0.193917 |
| 60.24893 | 74.49973 | 82.6955 | 64.08598 | 68.37597 | 71.11691 | 61.49709 | 48.13844 | 76.90323 |
| 57.63021 | 52.83306 | 52.06735 | 52.07489 | 54.77267 | 55.2536 | 51.09282 | 59.77919 | 52.57799 |
| 52.27929 | 45.59806 | 45.82722 | 45.32426 | 49.44015 | 50.16515 | 43.35737 | 50.18528 | 48.19305 |
| 54.5448 | 50.41167 | 48.51007 | 50.47149 | 56.24184 | 53.52924 | 52.05336 | 58.71701 | 51.50244 |
| 57.30656 | 53.21232 | 58.29754 | 56.17021 | 60.76904 | 59.18895 | 55.39483 | 64.78173 | 56.81816 |
| 76.28289 | 68.7812 | 74.31521 | 77.12681 | 83.6445 | 81.58448 | 76.05677 | 91.5533 | 81.66964 |
| 79.12017 | 73.62399 | 82.3141 | 79.49617 | 87.51874 | 82.3779 | 79.79676 | 93.33503 | 81.3387 |
| 50.11088 | 44.7423 | 50.87497 | 48.70468 | 53.3688 | 50.39788 | 49.89725 | 55.14213 | 49.40305 |
| 65.61343 | 60.71961 | 65.85923 | 66.68936 | 71.90206 | 67.65207 | 68.22935 | 77.5736 | 67.13933 |
| 57.63021 | 51.79255 | 52.83246 | 59.31575 | 61.79202 | 58.55422 | 55.60942 | 64.21066 | 56.7044 |
| 65.80761 | 61.59481 | 60.70218 | 64.01362 | 69.89964 | 66.56244 | 64.56089 | 73.05076 | 64.59523 |

| | | | | | | | | |
|---|---|---|---|---|---|---|---|---|
| 130.5364 | 118.6386 | 108.2087 | 128.4766 | 138.9722 | 125.0426 | 124.5643 | 151.4467 | 121.2066 |
| 44.32844 | 39.88006 | 37.88794 | 43.14894 | 44.61911 | 42.17808 | 40.44508 | 46.86168 | 40.49871 |
| 13.38807 | 12.64182 | 11.15871 | 12.92936 | 12.87424 | 12.22921 | 12.26228 | 13.8769 | 12.19305 |
| 81.15913 | 73.01135 | 74.93127 | 75.13532 | 79.8029 | 78.26271 | 76.71076 | 85.64848 | 76.52973 |
| 78.19239 | 68.39222 | 70.53933 | 71.97957 | 75.93954 | 76.96151 | 71.8365 | 81.50254 | 71.68974 |
| 33.70213 | 29.35819 | 36.78499 | 32.52766 | 38.30713 | 37.02615 | 36.50071 | 40.24873 | 33.94197 |
| 37.87714 | 40.08428 | 33.76428 | 39.12511 | 44.89117 | 37.10021 | 40.32245 | 44.10914 | 37.40649 |
| 85.71172 | 75.94814 | 76.33232 | 79.16936 | 81.93591 | 77.59624 | 78.19245 | 89.34899 | 73.72709 |
| 11.2736 | 13.72123 | 10.78112 | 11.62213 | 13.2769 | 11.19248 | 12.68124 | 15.35025 | 10.18052 |
| 15.25442 | 14.73258 | 10.84074 | 12.45957 | 13.35308 | 13.55157 | 13.41697 | 14.71066 | 14.12698 |
| 25.43842 | 22.12318 | 19.05824 | 21.2834 | 22.52721 | 23.74963 | 21.89838 | 25.00127 | 24.04481 |
| 60.44591 | 54.94327 | 55.26691 | 55.75149 | 59.65901 | 60.29974 | 56.40647 | 65.14721 | 55.90807 |
| 76.02397 | 67.01135 | 69.18797 | 68.22128 | 74.93833 | 72.77226 | 72.38831 | 79.96066 | 71.19334 |
| 75.99161 | 63.71475 | 70.05244 | 65.72936 | 70.25877 | 73.22715 | 67.99432 | 78.15609 | 70.30394 |
| 10.20557 | 9.41329 | 9.152526 | 9.036255 | 8.817171 | 9.624684 | 10.24922 | 10.44137 | 10.12881 |
| 44.79233 | 38.63533 | 38.68286 | 38.57362 | 41.43047 | 42.83397 | 39.11666 | 43.66371 | 41.87417 |
| 64.63171 | 59.67909 | 66.03809 | 65.86213 | 72.57678 | 70.34969 | 67.34034 | 75.28934 | 71.51393 |
| 1505.279 | 1365.238 | 1390.269 | 1430.418 | 1543.529 | 1489.264 | 1438.375 | 1649.092 | 1446.41 |
| 33.91789 | 31.65316 | 43.55175 | 32.5583 | 35.4341 | 36.08463 | 35.26426 | 40.77411 | 35.40017 |
| 9.440695 | 9.080713 | 8.634833 | 10.0606 | 8.695284 | 10.25307 | 9.745444 | 12.40355 | 11.87245 |
| 0.737261 | 1.539384 | 0.828407 | 0.911694 | 1.0948 | 1.204937 | 1.087255 | 0.90948 | 0.865096 |
| 70.18759 | 61.4295 | 64.52774 | 63.08426 | 67.85369 | 69.02733 | 63.33466 | 70.68655 | 65.43292 |
| 147.8022 | 136.27 | 146.5872 | 139.2957 | 145.0839 | 150.4859 | 139.0859 | 157.2331 | 147.6781 |
| 33.51873 | 32.56726 | 29.04444 | 32.68085 | 32.00605 | 33.91596 | 29.65427 | 32.45939 | 34.10744 |
| 4.44E-05 | 3.76E-05 | 8.06E-05 | 7.98E-05 | 2.33E-05 | 3.31E-05 | 4.15E-05 | 2.46E-05 | 0.000107 |
| 0.000303 | 0.00044 | 0.000603 | 0.000699 | 0.000156 | 0.000327 | 0.000417 | 9.32E-05 | 0.000656 |
| 0.005042 | 0.004534 | 0.006155 | 0.006587 | 0.002376 | 0.003764 | 0.005215 | 0.001998 | 0.006415 |
| 0.008298 | 0.009164 | 0.013411 | 0.010991 | 0.005685 | 0.009243 | 0.012007 | 0.004739 | 0.013422 |
| 0.006426 | 0.005732 | 0.006936 | 0.007162 | 0.001276 | 0.003348 | 0.006001 | 0.001423 | 0.009107 |
| 0.000933 | 0.000929 | 0.001426 | 0.001364 | 0.000432 | 0.000689 | 0.00119 | 0.000352 | 0.001386 |
| 0.015613 | 0.013105 | 0.015395 | 0.015712 | 0.005909 | 0.009891 | 0.013082 | 0.0057 | 0.016543 |
| 0.02022 | 0.015421 | 0.014985 | 0.014104 | 0.00624 | 0.011212 | 0.013117 | 0.007167 | 0.014591 |
| 0.000655 | 0.000614 | 0.000783 | 0.000732 | 0.000297 | 0.000487 | 0.000662 | 0.000261 | 0.000785 |
| 0.029838 | 0.023876 | 0.027745 | 0.027027 | 0.010534 | 0.01806 | 0.024197 | 0.011846 | 0.030711 |
| 0.027041 | 0.02118 | 0.020321 | 0.018397 | 0.009725 | 0.015179 | 0.016446 | 0.0122 | 0.019091 |
| 0.003429 | 0.004088 | 0.005036 | 0.004287 | 0.00318 | 0.003679 | 0.004431 | 0.002628 | 0.004206 |
| 0.004511 | 0.004595 | 0.005065 | 0.004413 | 0.003409 | 0.003847 | 0.004414 | 0.002704 | 0.004394 |
| 0.001689 | 0.002482 | 0.002762 | 0.002832 | 0.002131 | 0.002195 | 0.002663 | 0.001773 | 0.001935 |
| 0.033103 | 0.026677 | 0.028155 | 0.026941 | 0.017523 | 0.023353 | 0.024913 | 0.016013 | 0.029261 |
| 0.010228 | 0.010895 | 0.01237 | 0.012758 | 0.007078 | 0.008883 | 0.010193 | 0.006934 | 0.012025 |
| 0.078982 | 0.053566 | 0.046661 | 0.044647 | 0.033964 | 0.042769 | 0.042093 | 0.034676 | 0.049092 |
| 0.026214 | 0.028688 | 0.032847 | 0.032684 | 0.020705 | 0.024782 | 0.028923 | 0.020047 | 0.031051 |
| 0.067053 | 0.05031 | 0.049602 | 0.048598 | 0.032411 | 0.041473 | 0.044747 | 0.033467 | 0.052488 |
| 0.05986 | 0.043885 | 0.042249 | 0.041675 | 0.026356 | 0.034919 | 0.036421 | 0.028329 | 0.043309 |
| 0.049767 | 0.037628 | 0.034122 | 0.031173 | 0.021723 | 0.029091 | 0.029493 | 0.023831 | 0.032561 |
| 0.03141 | 0.023483 | 0.023392 | 0.021749 | 0.012497 | 0.018702 | 0.021161 | 0.011375 | 0.023096 |
| 0.00968 | 0.007757 | 0.008238 | 0.007822 | 0.004745 | 0.005493 | 0.006564 | 0.004514 | 0.007293 |
| 0.071395 | 0.042322 | 0.032573 | 0.029008 | 0.025559 | 0.030922 | 0.028841 | 0.026522 | 0.032525 |
| 0.015295 | 0.013765 | 0.013352 | 0.012715 | 0.007233 | 0.009621 | 0.011379 | 0.008323 | 0.01082 |

| | | | | | | | | |
|---|---|---|---|---|---|---|---|---|
| 0.039044 | 0.029746 | 0.026802 | 0.023908 | 0.020247 | 0.024038 | 0.023762 | 0.022046 | 0.024659 |
| 0.001756 | 0.002315 | 0.002119 | 0.001669 | 0.001833 | 0.002056 | 0.00206 | 0.001641 | 0.001596 |
| 0.000872 | 0.000633 | 0.000459 | 0.000315 | 0.000336 | 0.000474 | 0.000404 | 0.000299 | 0.000322 |
| 0.000547 | 0.000604 | 0.000371 | 0.000357 | 0.000328 | 0.0004 | 0.000332 | 0.000338 | 0.000311 |
| 0.002754 | 0.001495 | 0.000765 | 0.000326 | 0.000336 | 0.000679 | 0.000437 | 0.000246 | 0.000502 |
| 0.0139 | 0.011549 | 0.010697 | 0.00666 | 0.006466 | 0.009807 | 0.006429 | 0.006033 | 0.007376 |
| 0.010554 | 0.011966 | 0.009949 | 0.007272 | 0.007982 | 0.009885 | 0.009618 | 0.009143 | 0.00817 |
| 0.723358 | 0.568488 | 0.548473 | 0.498676 | 0.335744 | 0.448274 | 0.470556 | 0.341203 | 0.528024 |
| 0.012355 | 0.0096 | 0.006526 | 0.002564 | 0.002728 | 0.005443 | 0.003755 | 0.002958 | 0.003381 |
| 0.007797 | 0.004949 | 0.003581 | 0.002256 | 0.002073 | 0.002913 | 0.001943 | 0.001669 | 0.002872 |
| 0.00058 | 0.000493 | 0.000336 | 0.000244 | 0.000195 | 0.000314 | 0.000219 | 0.000197 | 0.000206 |
| 0.005628 | 0.005101 | 0.004271 | 0.002431 | 0.002573 | 0.003724 | 0.002911 | 0.002519 | 0.002986 |
| 0.008609 | 0.007533 | 0.006884 | 0.005232 | 0.004691 | 0.006566 | 0.00443 | 0.004276 | 0.00572 |
| 0.000147 | 0.00018 | 0.000132 | 7.88E-05 | 0.000109 | 0.000109 | 0.000115 | 8.36E-05 | 7.69E-05 |
| 0.001949 | 0.002043 | 0.001601 | 0.000852 | 0.000889 | 0.001364 | 0.001324 | 0.000846 | 0.001136 |
| 0.000479 | 0.000562 | 0.000453 | 0.000383 | 0.000297 | 0.000439 | 0.000321 | 0.000288 | 0.00033 |
| 0.008372 | 0.009581 | 0.009311 | 0.005121 | 0.005806 | 0.008607 | 0.007792 | 0.007236 | 0.006236 |
| 0.001439 | 0.001837 | 0.001848 | 0.001114 | 0.001163 | 0.001651 | 0.001392 | 0.001088 | 0.00109 |
| 8E-05 | 7.94E-05 | 5.77E-05 | 4.48E-05 | 2.3E-05 | 3.5E-05 | 3.27E-05 | 2.79E-05 | 3.97E-05 |
| 0.000103 | 0.000164 | 0.000127 | 0.000131 | 0.000122 | 0.000148 | 0.000116 | 0.000105 | 9.42E-05 |
| 0.001765 | 0.001726 | 0.001448 | 0.001067 | 0.00137 | 0.001393 | 0.001306 | 0.001171 | 0.000995 |
| 0.002224 | 0.00289 | 0.002201 | 0.001651 | 0.002065 | 0.001895 | 0.00237 | 0.001607 | 0.001573 |
| 0.025374 | 0.018266 | 0.014269 | 0.010844 | 0.012943 | 0.014375 | 0.010874 | 0.010445 | 0.011482 |

| 23 (JAK2A | 30 (JAK2A | 34 (JAK2A | Average (C | Average (C | Average (C | Average (C | Average (J | Average (J |
|---|---|---|---|---|---|---|---|---|
| JAK2A-GH | JAK2A-GH | JAK2A-GH-Ins | | | | | | |
| 23.7775 | 22.40708 | 22.65351 | 26.38945 | 36.37608 | 28.31796 | 28.23279 | 29.72498 | 18.77412 |
| 8.026205 | 6.581037 | 7.322811 | 10.13267 | 8.434683 | 9.044724 | 9.659343 | 10.07688 | 6.603131 |
| 544.5241 | 536.4349 | 527.201 | 487.4935 | 496.8112 | 474.9191 | 527.3277 | 531.0347 | 470.616 |
| 450.0371 | 433.2743 | 429.3958 | 423.9341 | 431.1402 | 431.8676 | 446.3545 | 453.2063 | 397.8794 |
| 52.86527 | 40.88496 | 47.97041 | 46.07039 | 37.76457 | 35.4045 | 40.60675 | 57.29517 | 44.13752 |
| 0.765093 | 0.688597 | 0.900518 | 0.818908 | 1.143316 | 1.002392 | 0.98116 | 1.045108 | 0.753879 |
| 1079.995 | 1040.271 | 1035.444 | 994.8389 | 1011.67 | 980.5562 | 1053.162 | 1082.383 | 938.764 |
| 64.7737 | 45.91135 | 47.01268 | 87.65647 | 75.1071 | 87.36085 | 93.013 | 77.14543 | 65.50671 |
| 89.01382 | 72.81425 | 43.77235 | 88.98636 | 66.89345 | 82.68095 | 80.67164 | 94.37329 | 90.74731 |
| 33.63316 | 30.33303 | 23.76245 | 68.97303 | 58.21319 | 72.22821 | 76.05607 | 49.10335 | 46.21311 |
| 281.334 | 258.5935 | 224.7634 | 518.2212 | 474.5442 | 579.1316 | 599.1189 | 349.0848 | 326.2005 |
| 278.0181 | 238.2842 | 252.3229 | 443.7461 | 430.6938 | 463.6431 | 493.2077 | 323.1631 | 304.9141 |
| 42.66603 | 30.66996 | 18.5846 | 43.14836 | 32.41841 | 38.86901 | 42.31604 | 51.62543 | 46.06445 |
| 170.1953 | 133.836 | 160.1794 | 104.4493 | 117.8833 | 135.3475 | 142.9264 | 127.9917 | 116.891 |
| 115.1977 | 86.94658 | 96.48624 | 92.60986 | 91.82438 | 88.89004 | 104.9609 | 102.2082 | 98.71918 |
| 1074.832 | 897.3888 | 866.884 | 1447.791 | 1347.578 | 1548.151 | 1632.271 | 1174.695 | 1095.256 |
| 0.08377 | 0.070986 | 0.047547 | 0.138584 | 0.111523 | 0.1797 | 0.07136 | 0.104113 | 0.081582 |
| 4.276391 | 4.816901 | 2.903969 | 7.762374 | 7.233553 | 8.991008 | 5.674994 | 6.236883 | 5.062222 |
| 36.71199 | 40.82897 | 18.35032 | 61.06166 | 58.86101 | 69.86727 | 44.6815 | 45.02871 | 39.69694 |
| 21.67342 | 22.37746 | 11.79104 | 40.65275 | 38.26533 | 47.1722 | 27.27746 | 29.4187 | 25.06833 |
| 0.348875 | 0.608934 | 0.505198 | 0.638276 | 0.662813 | 0.740851 | 0.51841 | 0.639076 | 0.537992 |
| 0.826354 | 0.821891 | 0.552676 | 1.492858 | 1.077589 | 1.804578 | 0.828917 | 1.403331 | 1.071761 |
| 0.159842 | 0.158125 | 0.124802 | 0.270252 | 0.206873 | 0.396298 | 0.152004 | 0.317687 | 0.230601 |
| 0.299481 | 0.397183 | 0.254443 | 0.492946 | 0.420423 | 0.774822 | 0.34046 | 0.502152 | 0.37182 |
| 3.25513 | 2.699396 | 1.204225 | 5.704753 | 4.325995 | 7.894106 | 2.798703 | 4.311516 | 3.37612 |
| 1.910359 | 1.682173 | 0.793982 | 3.231242 | 2.654362 | 4.560537 | 1.757135 | 2.644807 | 2.278729 |
| 3.257355 | 3.62173 | 1.925608 | 5.21008 | 4.862611 | 7.21168 | 3.998168 | 4.157574 | 3.228947 |
| 2.425216 | 2.58833 | 1.862919 | 2.877337 | 2.86756 | 3.722546 | 2.556185 | 2.79696 | 2.384302 |
| 13.20964 | 12.50946 | 4.899872 | 21.23967 | 17.45298 | 30.34714 | 13.75681 | 14.98854 | 11.2267 |
| 67.68356 | 70.21328 | 30.09987 | 82.11141 | 80.24267 | 90.80866 | 73.67458 | 68.22577 | 64.56449 |
| 48.72682 | 42.73642 | 15.50397 | 67.46744 | 63.25432 | 74.51042 | 50.11818 | 52.59953 | 45.69688 |
| 13.91496 | 16.52958 | 10.18694 | 22.0143 | 18.10144 | 28.18918 | 18.14134 | 19.33619 | 15.60092 |
| 17.57726 | 14.87324 | 4.784635 | 30.54903 | 25.45813 | 33.08716 | 17.09628 | 21.78593 | 19.27845 |
| 5.43115 | 12.64708 | 10.59257 | 10.9542 | 9.443012 | 13.01634 | 9.54106 | 11.44486 | 9.562475 |
| 1.129172 | 0.914125 | 0.277029 | 2.132186 | 1.728444 | 2.590061 | 1.20503 | 1.509447 | 1.26012 |
| 0.617429 | 0.58165 | 0.380512 | 0.991079 | 0.851615 | 1.203262 | 0.574109 | 0.874374 | 0.62881 |
| 1.475377 | 1.618914 | 0.516031 | 2.46762 | 2.169325 | 3.406464 | 1.533497 | 2.051627 | 1.592764 |
| 0.277676 | 0.245312 | 0.091521 | 0.376671 | 0.363582 | 0.461031 | 0.259636 | 0.38823 | 0.282951 |
| 0.042452 | 0.064757 | 0.034479 | 0.099845 | 0.096499 | 0.122842 | 0.070045 | 0.087023 | 0.072319 |
| 1.094685 | 1.29006 | 0.486069 | 1.577325 | 1.398775 | 1.992159 | 1.118025 | 1.559248 | 1.185375 |
| 0.080321 | 0.102109 | 0.048722 | 0.093899 | 0.078695 | 0.103788 | 0.075231 | 0.149774 | 0.082965 |
| 3.163906 | 3.015694 | 1.336056 | 6.598044 | 5.115401 | 9.232576 | 3.339677 | 4.935538 | 3.840391 |
| 1.945958 | 1.3934 | 0.696492 | 3.203651 | 2.524851 | 4.60729 | 1.710519 | 2.812193 | 2.382888 |
| 0.051642 | 0.058527 | 0.034479 | 0.122738 | 0.07575 | 0.133717 | 0.050751 | 0.092157 | 0.071053 |
| 0.040161 | 0.029891 | 0.02021 | 0.067314 | 0.057381 | 0.092382 | 0.042932 | 0.069277 | 0.041568 |
| 0.455451 | 0.46334 | 0.21872 | 0.67415 | 0.524899 | 0.97294 | 0.398391 | 0.675248 | 0.554986 |
| 2.509765 | 3.172636 | 1.627836 | 4.225394 | 3.903433 | 5.640118 | 3.102428 | 3.296745 | 2.531683 |

| | | | | | | | | |
|---|---|---|---|---|---|---|---|---|
| 41.42893 | 34.91348 | 12.93419 | 60.40727 | 54.75048 | 69.23927 | 41.18531 | 44.64071 | 39.7562 |
| 0.119347 | 0.125771 | 0.10936 | 0.166387 | 0.144002 | 0.173702 | 0.156006 | 0.175132 | 0.149855 |
| 4.599011 | 5.169417 | 3.187452 | 7.80553 | 7.150185 | 9.217911 | 4.800022 | 7.527664 | 5.292268 |
| 0.291471 | 0.485795 | 0.52548 | 0.570133 | 0.607735 | 0.653423 | 0.510451 | 0.571129 | 0.501254 |
| 0.065503 | 0.090905 | 0.074881 | 0.088114 | 0.070546 | 0.084163 | 0.058563 | 0.096984 | 0.066859 |
| 0.440766 | 0.605795 | 0.554059 | 0.735793 | 0.807431 | 0.988863 | 0.642478 | 0.656355 | 0.565381 |
| 2.112386 | 2.065594 | 0.897234 | 3.040493 | 2.800956 | 3.655817 | 1.959323 | 3.380407 | 2.243504 |
| 0.229617 | 0.416016 | 0.154533 | 0.663541 | 0.652832 | 0.747991 | 0.445475 | 0.554428 | 0.46941 |
| 7.54487 | 7.675654 | 2.022868 | 10.56933 | 9.097815 | 13.00426 | 6.777317 | 9.963756 | 7.738528 |
| 334.1433 | 342.3221 | 158.6519 | 508.7573 | 464.0674 | 599.8347 | 368.7442 | 406.1205 | 348.0049 |
| 0.084326 | 0.098366 | 0.032082 | 0.084595 | 0.07489 | 0.139488 | 0.061775 | 0.102675 | 0.079496 |
| 0.091802 | 0.084676 | 0.080827 | 0.077294 | 0.083496 | 0.10127 | 0.063006 | 0.083254 | 0.078384 |
| 5.273177 | 10.57787 | 9.578489 | 8.878227 | 8.140593 | 11.38267 | 8.228045 | 9.658701 | 8.038079 |
| 0.051642 | 0.082189 | 0.079652 | 0.080561 | 0.079282 | 0.078866 | 0.064792 | 0.093539 | 0.069528 |
| 0.639234 | 0.89167 | 0.85114 | 0.900847 | 0.996922 | 1.159426 | 0.918803 | 1.059574 | 0.838106 |
| 0.096386 | 0.07596 | 0.028533 | 0.165063 | 0.127535 | 0.168734 | 0.090401 | 0.115636 | 0.093554 |
| 0.476143 | 0.488209 | 0.14619 | 0.992348 | 0.762643 | 1.157657 | 0.507258 | 0.706551 | 0.623396 |
| 12.3508 | 12.06761 | 3.987196 | 19.48377 | 17.31691 | 25.48035 | 11.90361 | 16.89239 | 12.94368 |
| 0.138282 | 0.224137 | 0.156907 | 0.237383 | 0.226253 | 0.336431 | 0.198793 | 0.279171 | 0.27421 |
| 2.19738 | 1.725875 | 0.669065 | 4.548527 | 3.511244 | 5.174518 | 2.212804 | 3.048004 | 2.600325 |
| 0.414289 | 0.550262 | 0.186592 | 0.987808 | 0.905048 | 1.061132 | 0.556709 | 0.742684 | 0.587251 |
| 0.842373 | 0.775292 | 0.242458 | 1.775199 | 1.369797 | 1.995699 | 0.939424 | 1.328514 | 1.14848 |
| 163.1598 | 119.4324 | 151.4446 | 122.0207 | 134.9788 | 123.4222 | 134.2779 | 123.1167 | 140.0378 |
| 9.213402 | 8.182329 | 11.23337 | 7.351227 | 6.843928 | 7.642159 | 6.173353 | 9.98568 | 9.427112 |
| 0.435387 | 0.363742 | 0.406126 | 0.435605 | 0.405909 | 0.359117 | 0.393203 | 0.419708 | 0.412891 |
| 604.2404 | 483.7138 | 537.659 | 475.5674 | 498.8727 | 474.6587 | 500.2426 | 497.7056 | 525.6733 |
| 0.877036 | 0.571247 | 0.773421 | 0.911621 | 0.782153 | 0.918632 | 0.946987 | 0.803937 | 1.079092 |
| 5.481186 | 3.223909 | 3.552801 | 3.606685 | 3.117813 | 2.857698 | 4.447736 | 3.778019 | 3.718334 |
| 2.051675 | 1.628981 | 2.492968 | 3.46165 | 3.061484 | 2.200097 | 3.235859 | 3.348909 | 4.120182 |
| 1.725077 | 1.31052 | 1.715042 | 1.946615 | 1.749113 | 1.577991 | 1.538185 | 2.252347 | 2.026998 |
| 2.247912 | 2.267775 | 2.919905 | 2.598598 | 2.643662 | 2.350548 | 2.32355 | 3.070797 | 2.798607 |
| 12.47474 | 13.09958 | 11.70965 | 14.99213 | 14.72516 | 14.5261 | 14.48057 | 16.56112 | 14.70503 |
| 4.597423 | 6.872557 | 4.917283 | 5.299348 | 3.797391 | 3.767367 | 4.210205 | 5.487611 | 5.600539 |
| 116.7448 | 97.1289 | 115.0155 | 86.38246 | 92.57093 | 92.91221 | 99.72476 | 90.55585 | 95.59397 |
| 130.2216 | 101.5634 | 101.7783 | 103.4428 | 104.9858 | 98.55768 | 102.122 | 103.8432 | 111.1119 |
| 0.056227 | 0.053906 | 0.050889 | 0.066935 | 0.067793 | 0.045535 | 0.052162 | 0.060569 | 0.072015 |
| 80.04897 | 58.20998 | 61.98093 | 59.81195 | 58.97186 | 57.21729 | 58.96312 | 64.61783 | 62.81963 |
| 0.144765 | 0.16215 | 0.237926 | 0.142115 | 0.154965 | 0.14456 | 0.17166 | 0.164291 | 0.197003 |
| 74.76031 | 69.64241 | 67.43027 | 63.09692 | 70.01597 | 66.15957 | 67.18138 | 69.63902 | 71.95218 |
| 55.61511 | 51.20319 | 59.67679 | 52.97249 | 61.78393 | 52.42947 | 51.24863 | 56.63188 | 54.92705 |
| 47.58205 | 45.44565 | 48.97714 | 42.63705 | 46.31128 | 43.16101 | 40.87033 | 47.72279 | 46.54963 |
| 52.9693 | 50.87536 | 57.54588 | 53.47607 | 58.57995 | 51.75323 | 52.60551 | 55.39841 | 52.20752 |
| 60.19481 | 55.41377 | 61.77388 | 56.79678 | 62.76163 | 56.56077 | 56.99637 | 58.82358 | 56.83826 |
| 81.85006 | 78.03415 | 79.71187 | 76.31542 | 82.19335 | 78.94428 | 80.35675 | 79.12334 | 74.22644 |
| 83.65643 | 80.76949 | 86.69089 | 81.2984 | 89.12574 | 81.73091 | 81.26975 | 83.42812 | 79.03327 |
| 49.53719 | 48.92886 | 53.58847 | 48.91242 | 54.45932 | 49.15215 | 49.33093 | 50.49494 | 48.25753 |
| 67.2928 | 67.05179 | 74.07454 | 66.26106 | 73.07428 | 66.7411 | 66.15334 | 68.77585 | 64.91015 |
| 61.63991 | 59.41947 | 61.34544 | 56.61588 | 62.72391 | 58.41651 | 56.51969 | 59.06388 | 56.15273 |
| 67.47344 | 64.75697 | 71.51519 | 64.71065 | 73.0301 | 64.82908 | 64.05451 | 67.49208 | 64.06931 |

| | | | | | | | | |
|---|---|---|---|---|---|---|---|---|
| 138.8784 | 123.1417 | 138.0019 | 123.6208 | 137.1616 | 122.6269 | 127.8123 | 129.1751 | 125.4352 |
| 43.73554 | 40.856 | 45.85406 | 41.1709 | 44.61454 | 40.45622 | 40.74123 | 42.74768 | 41.80741 |
| 11.90083 | 11.89414 | 13.58597 | 12.46733 | 14.44712 | 11.37343 | 11.91499 | 13.5293 | 13.46186 |
| 79.58678 | 76.74331 | 82.89132 | 79.67085 | 87.36467 | 78.65864 | 78.50989 | 80.56676 | 78.07613 |
| 74.81582 | 75.65737 | 77.94175 | 74.52627 | 84.79596 | 73.80005 | 73.54195 | 77.73964 | 74.12015 |
| 35.10744 | 34.42231 | 37.65738 | 35.69323 | 38.5334 | 36.22981 | 36.55897 | 35.28112 | 34.16478 |
| 39.72963 | 35.21116 | 44.83934 | 36.81242 | 40.35684 | 32.96585 | 35.47503 | 42.50873 | 39.71567 |
| 85.48406 | 77.6346 | 88.44973 | 81.05989 | 86.28084 | 75.30291 | 77.4647 | 86.58033 | 82.08722 |
| 12.11334 | 10.50085 | 15.72816 | 10.68392 | 12.42391 | 9.441219 | 11.34499 | 13.07619 | 12.72796 |
| 11.06139 | 10.75697 | 13.56342 | 16.00037 | 17.2181 | 12.58456 | 14.04156 | 16.42076 | 16.36487 |
| 19.20071 | 18.65566 | 21.25274 | 26.91023 | 28.02083 | 23.63779 | 24.18873 | 26.47968 | 26.04424 |
| 59.06848 | 55.75185 | 64.25431 | 56.97233 | 62.75744 | 57.20023 | 56.5764 | 59.04332 | 56.66319 |
| 72.54191 | 71.15993 | 77.76135 | 72.73175 | 78.83618 | 70.83193 | 70.99282 | 73.96947 | 71.60316 |
| 68.66352 | 66.00683 | 73.57845 | 73.48437 | 80.74491 | 72.01892 | 73.09947 | 71.78358 | 72.33658 |
| 9.059504 | 10.28571 | 10.95446 | 8.566103 | 12.14095 | 9.298713 | 8.695499 | 10.49423 | 9.776239 |
| 37.17946 | 36.50199 | 41.00595 | 44.64802 | 47.48453 | 41.37485 | 41.38592 | 44.33837 | 43.34628 |
| 67.88784 | 68.07627 | 74.78484 | 65.63885 | 73.85623 | 68.80815 | 67.30291 | 68.60251 | 64.10854 |
| 1493.826 | 1425.155 | 1577.005 | 1460.654 | 1611.082 | 1440.329 | 1449.053 | 1519.292 | 1459.011 |
| 35.31995 | 34.21742 | 38.26621 | 33.20022 | 37.66481 | 35.85658 | 33.98048 | 35.5588 | 33.75666 |
| 10.68949 | 10.67501 | 9.479737 | 10.16166 | 13.74783 | 10.44718 | 10.6601 | 11.09662 | 9.408914 |
| 1.129516 | 0.936266 | 1.233448 | 0.734275 | 0.714596 | 0.586191 | 0.795985 | 0.924644 | 0.955228 |
| 66.07084 | 62.28799 | 69.58722 | 69.54142 | 75.68261 | 68.63024 | 67.25525 | 68.47597 | 67.14871 |
| 146.2983 | 137.9391 | 152.695 | 142.2961 | 158.8714 | 144.674 | 140.9478 | 149.154 | 142.6255 |
| 33.08855 | 29.82242 | 34.12841 | 28.65849 | 31.06156 | 29.15376 | 28.25598 | 33.09792 | 31.356 |
| 5.65E-05 | 9.15E-05 | 4.38E-05 | 2.88E-05 | 3.33E-05 | 8.33E-05 | 4.12E-05 | 6.48E-05 | 3.41E-05 |
| 0.000459 | 0.000518 | 0.000409 | 0.000307 | 0.000331 | 0.000601 | 0.000453 | 0.000453 | 0.000296 |
| 0.004898 | 0.004813 | 0.005026 | 0.006417 | 0.006466 | 0.007906 | 0.006759 | 0.004848 | 0.00432 |
| 0.008087 | 0.010737 | 0.012298 | 0.010203 | 0.009783 | 0.012057 | 0.011716 | 0.009389 | 0.008585 |
| 0.005154 | 0.00442 | 0.006504 | 0.00452 | 0.004488 | 0.007231 | 0.005429 | 0.006289 | 0.005113 |
| 0.000895 | 0.000934 | 0.001169 | 0.000985 | 0.00098 | 0.001305 | 0.001219 | 0.000915 | 0.000844 |
| 0.012354 | 0.011745 | 0.01308 | 0.015859 | 0.015769 | 0.019811 | 0.017418 | 0.01296 | 0.012481 |
| 0.012675 | 0.012953 | 0.012812 | 0.018569 | 0.019342 | 0.020644 | 0.019805 | 0.014088 | 0.014285 |
| 0.000468 | 0.000583 | 0.0007 | 0.000776 | 0.000723 | 0.000978 | 0.000734 | 0.000604 | 0.00057 |
| 0.022711 | 0.020453 | 0.02503 | 0.032786 | 0.032849 | 0.042258 | 0.035868 | 0.022967 | 0.023029 |
| 0.017106 | 0.017989 | 0.017457 | 0.02556 | 0.02633 | 0.027577 | 0.028385 | 0.019037 | 0.019765 |
| 0.00316 | 0.003996 | 0.004554 | 0.004468 | 0.004588 | 0.005034 | 0.004819 | 0.003777 | 0.003827 |
| 0.003166 | 0.004439 | 0.004373 | 0.005981 | 0.00634 | 0.006785 | 0.006315 | 0.004327 | 0.004408 |
| 0.002034 | 0.002247 | 0.002558 | 0.002353 | 0.002281 | 0.002255 | 0.002379 | 0.002239 | 0.002399 |
| 0.023219 | 0.02596 | 0.026191 | 0.037502 | 0.03938 | 0.046839 | 0.044227 | 0.02679 | 0.025759 |
| 0.008837 | 0.011274 | 0.011076 | 0.015676 | 0.015389 | 0.017824 | 0.01485 | 0.010209 | 0.010684 |
| 0.042081 | 0.046643 | 0.042334 | 0.093777 | 0.097767 | 0.117286 | 0.115348 | 0.055309 | 0.051815 |
| 0.026204 | 0.028829 | 0.029737 | 0.037015 | 0.037584 | 0.044251 | 0.043296 | 0.027473 | 0.027199 |
| 0.044387 | 0.046413 | 0.045635 | 0.084486 | 0.088593 | 0.110871 | 0.11062 | 0.051311 | 0.04967 |
| 0.037668 | 0.039988 | 0.039156 | 0.068627 | 0.074285 | 0.088638 | 0.095149 | 0.042607 | 0.042874 |
| 0.030725 | 0.032881 | 0.0308 | 0.053426 | 0.057257 | 0.066195 | 0.075877 | 0.034748 | 0.035266 |
| 0.018492 | 0.021969 | 0.02091 | 0.030517 | 0.033461 | 0.039641 | 0.041318 | 0.022195 | 0.021717 |
| 0.005429 | 0.007415 | 0.006876 | 0.0117 | 0.012233 | 0.013563 | 0.012462 | 0.007231 | 0.007527 |
| 0.029757 | 0.034058 | 0.030984 | 0.072294 | 0.076709 | 0.082649 | 0.095622 | 0.041907 | 0.041416 |
| 0.010133 | 0.012693 | 0.011571 | 0.015547 | 0.016327 | 0.017931 | 0.02134 | 0.011578 | 0.012193 |

| | | | | | | | | |
|---|---|---|---|---|---|---|---|---|
| 0.023698 | 0.025948 | 0.024462 | 0.038581 | 0.040798 | 0.043243 | 0.052924 | 0.028059 | 0.028478 |
| 0.001235 | 0.001798 | 0.001863 | 0.001871 | 0.002083 | 0.002021 | 0.002412 | 0.001971 | 0.001902 |
| 0.000293 | 0.000479 | 0.000353 | 0.000714 | 0.000724 | 0.000668 | 0.00076 | 0.000556 | 0.000535 |
| 0.000351 | 0.000386 | 0.000347 | 0.000562 | 0.000553 | 0.000521 | 0.000565 | 0.00043 | 0.000477 |
| 0.000406 | 0.000685 | 0.000477 | 0.001738 | 0.001778 | 0.001396 | 0.001828 | 0.001195 | 0.001148 |
| 0.008414 | 0.012059 | 0.007873 | 0.012031 | 0.013776 | 0.0141 | 0.015121 | 0.009719 | 0.01 |
| 0.007518 | 0.009118 | 0.009902 | 0.011274 | 0.012281 | 0.012489 | 0.014505 | 0.010227 | 0.009508 |
| 0.450801 | 0.509183 | 0.490534 | 0.785919 | 0.828748 | 0.951528 | 0.983699 | 0.540667 | 0.531284 |
| 0.004169 | 0.005819 | 0.004541 | 0.009553 | 0.010341 | 0.008225 | 0.010946 | 0.006658 | 0.006856 |
| 0.002583 | 0.003911 | 0.002315 | 0.00565 | 0.006388 | 0.00637 | 0.005714 | 0.004092 | 0.004121 |
| 0.000273 | 0.00033 | 0.000272 | 0.000444 | 0.000444 | 0.000395 | 0.000543 | 0.000358 | 0.000401 |
| 0.003009 | 0.003892 | 0.003502 | 0.005458 | 0.005869 | 0.005628 | 0.006929 | 0.004024 | 0.004 |
| 0.005862 | 0.008001 | 0.00553 | 0.008467 | 0.00907 | 0.01054 | 0.011034 | 0.006646 | 0.006674 |
| 8.93E-05 | 0.000131 | 0.000115 | 0.000153 | 0.000147 | 0.000141 | 0.000175 | 0.000123 | 0.000128 |
| 0.000838 | 0.001094 | 0.001279 | 0.001603 | 0.001922 | 0.002026 | 0.001993 | 0.001642 | 0.001344 |
| 0.000424 | 0.000389 | 0.000389 | 0.000551 | 0.0005 | 0.000516 | 0.000569 | 0.000407 | 0.000466 |
| 0.0058 | 0.008068 | 0.007579 | 0.008359 | 0.009931 | 0.009526 | 0.011107 | 0.008115 | 0.007227 |
| 0.001039 | 0.001813 | 0.001587 | 0.001405 | 0.00167 | 0.001689 | 0.001838 | 0.001516 | 0.001337 |
| 3.75E-05 | 3.74E-05 | 3.63E-05 | 5.69E-05 | 5.36E-05 | 5.63E-05 | 7.04E-05 | 4.66E-05 | 5.24E-05 |
| 0.000117 | 0.000125 | 0.00015 | 0.000122 | 0.000109 | 0.000105 | 0.00012 | 0.000123 | 0.000138 |
| 0.001045 | 0.001725 | 0.001494 | 0.001558 | 0.001674 | 0.001987 | 0.002361 | 0.00156 | 0.001451 |
| 0.001441 | 0.002005 | 0.002134 | 0.002318 | 0.002553 | 0.002439 | 0.002615 | 0.002186 | 0.002156 |
| 0.012003 | 0.017325 | 0.01305 | 0.024068 | 0.026795 | 0.027233 | 0.028122 | 0.017697 | 0.016808 |

| Average (J. | Average (J. | SEM (Cont | SEM (Cont | SEM (Cont | SEM (Cont | SEM (JAK2 | SEM (JAK2 | SEM (JAK2 |
|---|---|---|---|---|---|---|---|---|
| 27.69639 | 23.60343 | 4.499254 | 19.17276 | 1.442731 | 2.58066 | 5.304693 | 1.495423 | 0.90352 |
| 8.758167 | 7.024262 | 1.50786 | 1.955485 | 0.909691 | 1.228968 | 1.146072 | 0.737638 | 0.286939 |
| 534.4924 | 520.0218 | 15.55712 | 23.03068 | 24.06849 | 14.29253 | 20.36434 | 27.19777 | 17.40352 |
| 456.9874 | 427.8218 | 5.775012 | 25.8874 | 18.74679 | 15.34304 | 15.04983 | 20.35129 | 13.06647 |
| 50.69852 | 45.56721 | 3.764809 | 3.041195 | 3.977185 | 1.632243 | 7.982045 | 6.458768 | 5.624206 |
| 0.840136 | 0.846222 | 0.102973 | 0.439593 | 0.061447 | 0.088461 | 0.146024 | 0.090173 | 0.063798 |
| 1079.473 | 1024.885 | 28.82238 | 63.67734 | 44.80456 | 31.54081 | 48.32994 | 54.88263 | 29.17909 |
| 51.1451 | 53.41168 | 14.76745 | 6.502675 | 13.8455 | 17.73789 | 12.2082 | 6.631499 | 2.833775 |
| 59.33035 | 74.83912 | 17.04683 | 1.593135 | 17.34355 | 21.8299 | 17.22582 | 15.54278 | 5.764932 |
| 34.62029 | 32.74967 | 11.83676 | 2.567044 | 14.65658 | 20.3767 | 4.491648 | 4.843886 | 3.52399 |
| 263.1561 | 279.263 | 113.6221 | 26.91168 | 119.058 | 175.5624 | 39.4104 | 29.08831 | 12.41308 |
| 251.4394 | 263.0966 | 79.06291 | 27.50697 | 79.34279 | 125.3869 | 36.30353 | 30.98846 | 9.689554 |
| 33.32483 | 33.91008 | 5.694674 | 3.318394 | 4.963532 | 6.793278 | 6.071101 | 5.518043 | 3.106335 |
| 145.3751 | 153.2397 | 5.345781 | 8.76633 | 2.20898 | 15.76233 | 2.084452 | 3.755632 | 8.595804 |
| 99.97055 | 100.8164 | 3.312547 | 10.2047 | 9.222989 | 7.098334 | 4.428659 | 6.375355 | 4.747023 |
| 938.3617 | 991.3262 | 234.0337 | 69.02213 | 252.3012 | 371.3791 | 108.6029 | 85.45935 | 34.38069 |
| 0.08609 | 0.075192 | 0.031667 | 0.012574 | 0.050818 | 0.003486 | 0.02219 | 0.011497 | 0.025029 |
| 4.56785 | 4.646611 | 0.852452 | 1.100771 | 1.725708 | 0.14616 | 1.073732 | 0.289486 | 0.511706 |
| 35.61142 | 36.78092 | 5.501265 | 6.043895 | 10.52243 | 1.737982 | 7.134157 | 4.155854 | 5.388367 |
| 20.58322 | 20.09157 | 4.572143 | 3.332299 | 7.827817 | 0.95819 | 4.759157 | 3.16316 | 3.370562 |
| 0.510115 | 0.500517 | 0.042829 | 0.021895 | 0.06398 | 0.022731 | 0.033951 | 0.042128 | 0.045119 |
| 0.89969 | 0.877728 | 0.300694 | 0.16037 | 0.464905 | 0.086901 | 0.281595 | 0.192221 | 0.150524 |
| 0.170926 | 0.19916 | 0.064937 | 0.043285 | 0.125781 | 0.035485 | 0.091389 | 0.051288 | 0.031099 |
| 0.428496 | 0.389456 | 0.087369 | 0.049194 | 0.181401 | 0.025002 | 0.068838 | 0.05024 | 0.065282 |
| 2.720313 | 2.975259 | 1.523777 | 0.678947 | 2.361053 | 0.379864 | 1.040817 | 0.695124 | 0.562089 |
| 1.702039 | 1.653192 | 0.819737 | 0.271834 | 1.357616 | 0.182245 | 0.607196 | 0.483912 | 0.394615 |
| 3.649764 | 3.360676 | 0.475591 | 0.54087 | 1.363576 | 0.185369 | 0.544526 | 0.257283 | 0.555442 |
| 2.658062 | 2.478965 | 0.149029 | 0.240669 | 0.616708 | 0.259454 | 0.285922 | 0.192558 | 0.382618 |
| 13.34232 | 12.19903 | 3.312374 | 2.646418 | 7.433114 | 1.327387 | 2.828439 | 1.481458 | 2.759119 |
| 63.64561 | 61.46973 | 3.806887 | 3.434554 | 7.719832 | 2.299089 | 6.272729 | 4.631219 | 8.464829 |
| 38.60718 | 39.77522 | 4.522305 | 6.063493 | 10.41058 | 3.32214 | 8.710242 | 5.88333 | 8.278344 |
| 16.71736 | 15.31655 | 2.009552 | 1.407429 | 4.77167 | 1.685258 | 1.969457 | 1.088291 | 2.054458 |
| 13.41442 | 13.46672 | 4.930718 | 3.198167 | 6.033632 | 1.216213 | 4.401593 | 3.34847 | 3.219451 |
| 11.31786 | 9.49668 | 1.116349 | 0.880592 | 1.008053 | 0.514837 | 0.851244 | 0.653347 | 1.116374 |
| 0.887051 | 0.846287 | 0.415855 | 0.236718 | 0.591835 | 0.073472 | 0.368266 | 0.234071 | 0.203157 |
| 0.670799 | 0.596008 | 0.170055 | 0.098693 | 0.223117 | 0.044224 | 0.113638 | 0.05107 | 0.096027 |
| 1.458663 | 1.337976 | 0.396625 | 0.361076 | 0.80456 | 0.28145 | 0.44291 | 0.224539 | 0.349128 |
| 0.281547 | 0.253003 | 0.082021 | 0.067269 | 0.084608 | 0.031238 | 0.070442 | 0.041957 | 0.056176 |
| 0.043065 | 0.051401 | 0.018164 | 0.014192 | 0.027616 | 0.00412 | 0.014454 | 0.012607 | 0.011523 |
| 1.113063 | 0.983556 | 0.253959 | 0.222441 | 0.362196 | 0.222646 | 0.314706 | 0.169939 | 0.263281 |
| 0.119092 | 0.090221 | 0.014073 | 0.013444 | 0.012466 | 0.005563 | 0.02784 | 0.008266 | 0.017782 |
| 3.237692 | 3.240241 | 1.670144 | 0.736035 | 2.857471 | 0.45145 | 1.138944 | 0.739338 | 0.668799 |
| 1.748323 | 1.632447 | 0.791441 | 0.245255 | 1.304374 | 0.213773 | 0.641292 | 0.536359 | 0.408869 |
| 0.051604 | 0.051869 | 0.034243 | 0.006642 | 0.039169 | 0.011987 | 0.019967 | 0.018545 | 0.006839 |
| 0.036002 | 0.035982 | 0.012438 | 0.011158 | 0.021638 | 0.00441 | 0.017807 | 0.012468 | 0.005523 |
| 0.500588 | 0.416901 | 0.165928 | 0.084209 | 0.264175 | 0.031044 | 0.145777 | 0.090002 | 0.110309 |
| 2.853228 | 2.640928 | 0.362049 | 0.513154 | 1.156992 | 0.185893 | 0.419593 | 0.245366 | 0.425798 |

| | | | | | | | | |
|---|---|---|---|---|---|---|---|---|
| 33.44268 | 32.66232 | 6.013992 | 5.463655 | 11.67955 | 2.677592 | 8.061501 | 5.576342 | 7.175098 |
| 0.15384 | 0.130274 | 0.01136 | 0.013775 | 0.032858 | 0.022009 | 0.010189 | 0.013422 | 0.016915 |
| 5.568498 | 4.975145 | 0.960369 | 0.75089 | 1.630663 | 0.418493 | 1.076856 | 0.558031 | 0.830716 |
| 0.450839 | 0.443788 | 0.039791 | 0.051673 | 0.057823 | 0.022716 | 0.042451 | 0.04386 | 0.027798 |
| 0.069857 | 0.073057 | 0.00681 | 0.012599 | 0.015928 | 0.007228 | 0.004762 | 0.007495 | 0.00762 |
| 0.532181 | 0.527484 | 0.081404 | 0.066903 | 0.21039 | 0.04878 | 0.063906 | 0.043305 | 0.055205 |
| 2.32736 | 2.04738 | 0.461174 | 0.444837 | 0.641699 | 0.132102 | 0.482895 | 0.231637 | 0.490083 |
| 0.333412 | 0.262988 | 0.14203 | 0.076629 | 0.133946 | 0.017854 | 0.151551 | 0.121689 | 0.088788 |
| 6.828017 | 6.064287 | 1.846305 | 1.767654 | 2.609105 | 1.466826 | 2.373557 | 1.21848 | 1.668674 |
| 319.0001 | 308.4695 | 48.2408 | 40.00218 | 92.37753 | 16.98694 | 60.22942 | 37.08512 | 52.46009 |
| 0.085769 | 0.074266 | 0.02027 | 0.00818 | 0.041305 | 0.008556 | 0.015465 | 0.007954 | 0.015227 |
| 0.089957 | 0.089313 | 0.011254 | 0.01277 | 0.018837 | 0.011746 | 0.01174 | 0.008184 | 0.00729 |
| 9.292464 | 8.457432 | 1.022569 | 1.013939 | 1.091738 | 0.475411 | 0.855478 | 0.598433 | 1.130324 |
| 0.070528 | 0.062722 | 0.01298 | 0.010426 | 0.00807 | 0.006033 | 0.015817 | 0.004076 | 0.005583 |
| 0.864335 | 0.803168 | 0.040389 | 0.053965 | 0.136624 | 0.080359 | 0.061713 | 0.053416 | 0.068011 |
| 0.079959 | 0.083217 | 0.02605 | 0.010706 | 0.026436 | 0.010182 | 0.022528 | 0.012171 | 0.013198 |
| 0.368976 | 0.387673 | 0.209462 | 0.097182 | 0.233326 | 0.026888 | 0.160129 | 0.154171 | 0.073355 |
| 11.6721 | 10.44294 | 3.478481 | 2.823342 | 5.596615 | 1.981542 | 3.763297 | 2.134792 | 2.955578 |
| 0.245204 | 0.220653 | 0.030399 | 0.026187 | 0.070537 | 0.012751 | 0.042627 | 0.024297 | 0.035635 |
| 1.777376 | 1.667149 | 0.832559 | 0.471811 | 1.251371 | 0.134806 | 0.695377 | 0.517235 | 0.435854 |
| 0.373 | 0.409037 | 0.153432 | 0.12726 | 0.224834 | 0.034745 | 0.158354 | 0.121546 | 0.086299 |
| 0.740284 | 0.655205 | 0.280307 | 0.180552 | 0.409099 | 0.033976 | 0.293387 | 0.232856 | 0.169632 |
| 130.6042 | 135.8704 | 9.24141 | 3.010064 | 3.71273 | 9.248962 | 4.17002 | 5.214318 | 6.858236 |
| 8.705882 | 9.224141 | 1.014908 | 0.629505 | 1.039465 | 1.423055 | 0.552273 | 1.029097 | 0.794966 |
| 0.369367 | 0.396125 | 0.039411 | 0.01897 | 0.021945 | 0.021439 | 0.034818 | 0.019691 | 0.019984 |
| 508.9901 | 520.0221 | 15.98735 | 10.8826 | 8.744094 | 36.24611 | 18.97641 | 14.459 | 20.53748 |
| 0.825126 | 0.716343 | 0.140656 | 0.059744 | 0.131743 | 0.273486 | 0.042308 | 0.229936 | 0.119577 |
| 3.830005 | 3.741137 | 0.272883 | 0.294896 | 0.386502 | 1.03859 | 0.312362 | 0.398062 | 0.485085 |
| 3.046298 | 2.398304 | 0.442537 | 0.584627 | 0.184377 | 0.579297 | 0.318807 | 0.333438 | 0.185824 |
| 1.689629 | 1.695848 | 0.229392 | 0.064799 | 0.1984 | 0.168965 | 0.227005 | 0.186408 | 0.12663 |
| 2.321208 | 2.465723 | 0.392333 | 0.422946 | 0.201441 | 0.407927 | 0.329686 | 0.296341 | 0.067753 |
| 14.15842 | 13.098 | 1.507864 | 0.982319 | 1.692975 | 1.413728 | 1.807081 | 0.565556 | 0.741021 |
| 4.914427 | 4.639746 | 0.902426 | 0.300245 | 0.242725 | 0.717459 | 0.557473 | 0.762563 | 0.316071 |
| 99.75178 | 103.3383 | 6.738248 | 3.246001 | 2.601987 | 5.853185 | 3.174512 | 1.605512 | 4.417222 |
| 105.6957 | 109.1917 | 6.163493 | 2.994534 | 6.885297 | 8.193376 | 4.95315 | 4.051741 | 3.449728 |
| 0.059789 | 0.054926 | 0.005351 | 0.009253 | 0.002383 | 0.00587 | 0.00744 | 0.003219 | 0.006088 |
| 63.30973 | 65.64651 | 3.273425 | 2.553158 | 3.573026 | 4.945836 | 4.020155 | 3.522454 | 2.866729 |
| 0.154222 | 0.169994 | 0.006228 | 0.016352 | 0.024189 | 0.023469 | 0.011278 | 0.038467 | 0.022761 |
| 69.55429 | 67.37493 | 1.855736 | 5.682197 | 5.89431 | 8.252263 | 2.720971 | 3.60867 | 3.683359 |
| 53.05227 | 55.77046 | 1.515904 | 2.743374 | 0.777861 | 1.376229 | 1.067732 | 1.179884 | 0.823724 |
| 46.82283 | 48.07663 | 1.675086 | 1.084922 | 1.530831 | 1.951007 | 1.739283 | 2.140165 | 1.289726 |
| 52.1612 | 54.322 | 1.867332 | 2.154029 | 1.207952 | 1.208823 | 0.884211 | 0.767844 | 1.317585 |
| 57.96412 | 59.79647 | 1.586876 | 2.057049 | 1.295336 | 1.68369 | 0.962694 | 1.130953 | 0.982301 |
| 78.54555 | 82.5638 | 3.24471 | 3.303285 | 2.558373 | 1.185442 | 1.688918 | 1.381649 | 1.751243 |
| 82.30074 | 85.15811 | 2.340935 | 3.204729 | 1.044788 | 1.638696 | 1.140737 | 1.472984 | 1.438214 |
| 50.64872 | 51.31994 | 1.860464 | 1.950265 | 0.611433 | 0.486064 | 1.019652 | 1.053815 | 0.769966 |
| 68.06641 | 70.62641 | 1.560743 | 2.374212 | 1.238405 | 0.966365 | 1.190746 | 1.308584 | 1.041112 |
| 57.62077 | 60.66397 | 1.983657 | 3.485219 | 2.103276 | 2.292945 | 1.366847 | 1.31535 | 1.551081 |
| 65.14775 | 68.27832 | 2.664506 | 2.629256 | 0.840251 | 1.470774 | 1.094205 | 0.85702 | 1.516172 |

| | | | | | | | | |
|---|---|---|---|---|---|---|---|---|
| 125.0529 | 134.535 | 7.542231 | 8.236522 | 2.163766 | 3.714105 | 2.465125 | 2.110892 | 4.946895 |
| 41.65583 | 43.5612 | 1.140453 | 1.699702 | 0.76444 | 1.018738 | 1.101715 | 0.927946 | 1.160734 |
| 12.29076 | 12.69018 | 0.335065 | 0.540106 | 0.111427 | 0.391913 | 0.475335 | 0.290895 | 0.318936 |
| 76.96859 | 80.27992 | 1.800008 | 3.556175 | 1.256655 | 1.111163 | 1.507918 | 1.646173 | 0.92968 |
| 73.45129 | 76.32144 | 1.109877 | 2.262745 | 1.428803 | 2.824118 | 2.170736 | 1.82086 | 1.260277 |
| 36.22933 | 36.27557 | 1.123861 | 1.844511 | 1.347945 | 1.132718 | 1.299735 | 1.308588 | 0.975627 |
| 39.04064 | 40.25915 | 0.837187 | 1.792548 | 0.98957 | 1.330564 | 1.243799 | 0.460351 | 1.836901 |
| 78.64526 | 82.9289 | 1.315922 | 1.934199 | 1.515712 | 1.030115 | 2.424605 | 1.912765 | 0.942291 |
| 11.91077 | 12.77463 | 0.37172 | 0.488322 | 0.168728 | 0.334441 | 0.617088 | 0.510151 | 0.465581 |
| 12.72439 | 12.84388 | 1.094628 | 1.508664 | 0.575361 | 1.663663 | 0.67294 | 0.703517 | 0.508794 |
| 21.70337 | 21.63104 | 1.591089 | 2.507057 | 1.479373 | 1.586072 | 0.884554 | 1.391523 | 0.777078 |
| 57.47672 | 60.02598 | 1.77153 | 2.280222 | 1.003687 | 1.320561 | 1.277715 | 1.005578 | 1.042519 |
| 71.50163 | 74.52344 | 1.813306 | 3.242134 | 0.785005 | 1.514645 | 1.234167 | 1.476829 | 1.231369 |
| 69.45241 | 71.34177 | 2.906611 | 3.441556 | 2.228367 | 1.418552 | 2.67668 | 3.068083 | 1.249921 |
| 9.375971 | 10.17397 | 0.455275 | 1.862624 | 0.192757 | 0.143469 | 0.269663 | 0.367667 | 0.255148 |
| 40.12752 | 40.04506 | 1.846867 | 2.438823 | 1.920552 | 1.406449 | 1.488132 | 1.633232 | 0.85282 |
| 68.43341 | 71.51044 | 1.915102 | 1.803006 | 1.641202 | 1.384571 | 2.108133 | 1.582605 | 1.311174 |
| 1458.371 | 1518.298 | 40.03019 | 54.9099 | 26.84284 | 21.98406 | 22.13293 | 28.43255 | 26.48152 |
| 36.57861 | 36.79557 | 0.787326 | 0.873921 | 0.59507 | 0.742551 | 0.388724 | 0.678449 | 1.844845 |
| 9.477846 | 11.02405 | 0.859035 | 1.748206 | 0.308985 | 0.731089 | 0.294514 | 0.120748 | 0.341706 |
| 1.025419 | 1.014761 | 0.064364 | 0.052146 | 0.087667 | 0.030627 | 0.071368 | 0.156494 | 0.068048 |
| 65.56553 | 66.8131 | 2.863366 | 3.044848 | 3.163642 | 1.348477 | 1.685479 | 2.311526 | 1.213059 |
| 144.1077 | 148.3687 | 4.662902 | 3.931395 | 3.908091 | 3.587699 | 2.534813 | 4.101167 | 2.192675 |
| 31.46031 | 32.72124 | 1.517529 | 1.069817 | 1.494823 | 1.438877 | 1.555958 | 1.695592 | 0.919673 |
| 5.17E-05 | 6.47E-05 | 2.81E-06 | 7.55E-06 | 2.55E-05 | 1.52E-05 | 2.18E-05 | 5.1E-06 | 1.2E-05 |
| 0.000441 | 0.000427 | 5.89E-05 | 4.29E-05 | 0.000151 | 0.000176 | 0.000133 | 5.8E-05 | 9.69E-05 |
| 0.004819 | 0.00463 | 0.000823 | 0.00089 | 0.001679 | 0.002028 | 0.000811 | 0.000566 | 0.000779 |
| 0.010267 | 0.009857 | 0.000863 | 0.001245 | 0.002235 | 0.001995 | 0.001543 | 0.000812 | 0.001332 |
| 0.004945 | 0.005322 | 0.001002 | 0.000982 | 0.002133 | 0.002442 | 0.001623 | 0.001104 | 0.00114 |
| 0.00102 | 0.000947 | 0.000128 | 0.000115 | 0.000268 | 0.000377 | 0.000173 | 0.000144 | 0.000196 |
| 0.011998 | 0.011884 | 0.00277 | 0.002161 | 0.004075 | 0.006053 | 0.002192 | 0.001799 | 0.001844 |
| 0.011931 | 0.01204 | 0.003291 | 0.002192 | 0.00329 | 0.006529 | 0.0023 | 0.001936 | 0.001555 |
| 0.000592 | 0.000559 | 9.56E-05 | 6.6E-05 | 0.000266 | 0.000221 | 9.05E-05 | 7.34E-05 | 8.93E-05 |
| 0.021512 | 0.02215 | 0.006206 | 0.005436 | 0.011391 | 0.014139 | 0.003471 | 0.002999 | 0.003232 |
| 0.016014 | 0.016769 | 0.004738 | 0.002955 | 0.004882 | 0.009242 | 0.002646 | 0.002407 | 0.001798 |
| 0.004123 | 0.003709 | 0.00028 | 0.000285 | 0.00075 | 0.000585 | 0.000472 | 0.000213 | 0.000319 |
| 0.00423 | 0.003815 | 0.000412 | 0.00044 | 0.001091 | 0.000901 | 0.000337 | 0.000267 | 0.000281 |
| 0.002517 | 0.002109 | 0.000106 | 0.000126 | 0.000219 | 4.21E-05 | 0.000199 | 0.000232 | 0.000147 |
| 0.024177 | 0.024129 | 0.006038 | 0.005258 | 0.009536 | 0.01366 | 0.00292 | 0.002338 | 0.001857 |
| 0.010256 | 0.010029 | 0.000813 | 0.000537 | 0.00375 | 0.002106 | 0.000866 | 0.000682 | 0.001066 |
| 0.042027 | 0.042965 | 0.021221 | 0.017232 | 0.029949 | 0.04433 | 0.006772 | 0.007444 | 0.002167 |
| 0.027988 | 0.027173 | 0.003427 | 0.002723 | 0.008183 | 0.010236 | 0.002153 | 0.001657 | 0.002344 |
| 0.043366 | 0.044478 | 0.017137 | 0.014799 | 0.028135 | 0.043552 | 0.005697 | 0.005224 | 0.003096 |
| 0.036324 | 0.03769 | 0.014165 | 0.011079 | 0.020245 | 0.038549 | 0.005035 | 0.004895 | 0.002873 |
| 0.02912 | 0.030159 | 0.010829 | 0.007966 | 0.014065 | 0.031044 | 0.003989 | 0.004206 | 0.002051 |
| 0.0195 | 0.019168 | 0.006643 | 0.00476 | 0.007724 | 0.015165 | 0.00292 | 0.00284 | 0.001906 |
| 0.006572 | 0.006306 | 0.00065 | 9.6E-05 | 0.003 | 0.002359 | 0.000637 | 0.000671 | 0.000665 |
| 0.029381 | 0.030769 | 0.020646 | 0.016237 | 0.021011 | 0.040493 | 0.006631 | 0.007883 | 0.001175 |
| 0.01086 | 0.010708 | 0.002412 | 0.001788 | 0.002882 | 0.006792 | 0.00093 | 0.001014 | 0.001109 |

| | | | | | | | | |
|---|---|---|---|---|---|---|---|---|
| 0.023751 | 0.024163 | 0.007332 | 0.005231 | 0.00736 | 0.019331 | 0.002783 | 0.002883 | 0.001041 |
| 0.001947 | 0.001627 | 0.000272 | 0.00045 | 0.000208 | 0.000599 | 6.69E-05 | 0.000171 | 8.5E-05 |
| 0.000398 | 0.000349 | 0.000172 | 9.16E-05 | 0.0001 | 0.000255 | 8.06E-05 | 9.8E-05 | 3.19E-05 |
| 0.000358 | 0.000347 | 6.43E-05 | 2.82E-05 | 8.88E-05 | 0.000135 | 3.93E-05 | 4.53E-05 | 1.32E-05 |
| 0.000509 | 0.000463 | 0.000681 | 0.000347 | 0.000226 | 0.000891 | 0.000375 | 0.000448 | 9.03E-05 |
| 0.008012 | 0.008351 | 0.000894 | 0.000516 | 0.001741 | 0.003191 | 0.001166 | 0.00123 | 0.000926 |
| 0.008941 | 0.00877 | 0.001595 | 0.001771 | 0.001211 | 0.004236 | 0.000811 | 0.000848 | 0.000551 |
| 0.460344 | 0.463949 | 0.142884 | 0.111285 | 0.199012 | 0.343628 | 0.05741 | 0.057491 | 0.035363 |
| 0.004203 | 0.004174 | 0.002153 | 0.001291 | 0.001017 | 0.004017 | 0.001259 | 0.001756 | 0.000774 |
| 0.002553 | 0.00267 | 0.001207 | 0.000507 | 0.001425 | 0.001746 | 0.00084 | 0.001088 | 0.000306 |
| 0.000261 | 0.000255 | 7.09E-05 | 3.4E-05 | 6.5E-05 | 0.000158 | 4.64E-05 | 5.95E-05 | 2.73E-05 |
| 0.003182 | 0.003182 | 0.00096 | 0.000583 | 0.000777 | 0.002742 | 0.000458 | 0.000598 | 0.000353 |
| 0.005561 | 0.005878 | 0.000539 | 0.000404 | 0.001453 | 0.00263 | 0.00073 | 0.000653 | 0.000495 |
| 0.000109 | 9.93E-05 | 2.32E-05 | 6.38E-06 | 1.29E-05 | 4.31E-05 | 7.96E-06 | 1.65E-05 | 8.62E-06 |
| 0.001206 | 0.001039 | 0.000342 | 0.000392 | 0.000291 | 0.00072 | 0.000233 | 0.000276 | 0.000145 |
| 0.000379 | 0.000364 | 7.42E-05 | 1.15E-05 | 8.57E-05 | 0.000123 | 3.59E-05 | 3.82E-05 | 3.09E-05 |
| 0.007327 | 0.006984 | 0.000798 | 0.001305 | 0.001082 | 0.003141 | 0.00043 | 0.000795 | 0.000805 |
| 0.001433 | 0.001323 | 6.84E-05 | 0.000118 | 0.000156 | 0.000385 | 7.87E-05 | 0.000141 | 0.000141 |
| 3.86E-05 | 3.57E-05 | 1.38E-05 | 5.73E-06 | 1.43E-05 | 2.84E-05 | 8.55E-06 | 1.13E-05 | 5.89E-06 |
| 0.000129 | 0.000118 | 1.02E-05 | 8.7E-06 | 8E-06 | 1.48E-05 | 9.68E-06 | 1.02E-05 | 5.5E-06 |
| 0.001317 | 0.001286 | 0.000117 | 0.000148 | 0.00032 | 0.000509 | 0.000126 | 0.000137 | 6.65E-05 |
| 0.002036 | 0.001752 | 0.00023 | 0.0002 | 0.000125 | 0.000376 | 0.000143 | 0.000214 | 0.000124 |
| 0.012661 | 0.012861 | 0.00351 | 0.003155 | 0.00507 | 0.006948 | 0.001852 | 0.002399 | 0.000778 |

| SEM (JAK2 | Control - G | Control - G | Control - G | Control - G | Control - G | Control - G | Control - G | Control - G |
|---|---|---|---|---|---|---|---|---|
| 0.636106 | 0.961532 | 0.993504 | -1.01198 | 0.696522 | 0.955402 | 1.101205 | 0.661366 | 0.95342 |
| 0.331907 | 0.758377 | 0.962868 | 1.057358 | 0.82503 | 0.955402 | -1.04088 | 0.597694 | 0.95342 |
| 11.50408 | 0.091807 | 0.303802 | 1.110353 | 0.195245 | 0.815246 | 1.081712 | 0.678503 | 0.95342 |
| 6.950235 | 0.553523 | 0.962868 | 1.033545 | 0.361274 | 0.915711 | 1.052887 | 0.745038 | 0.95342 |
| 3.438001 | 0.363639 | 0.84849 | 1.164346 | 0.477957 | 0.934464 | -1.12581 | 0.111624 | 0.95342 |
| 0.055028 | 0.893379 | 0.962868 | -1.02881 | 0.365199 | 0.915711 | 1.213272 | 0.300127 | 0.95342 |
| 19.42757 | 0.262949 | 0.724186 | 1.074046 | 0.366613 | 0.915711 | 1.058626 | 0.823818 | 0.95342 |
| 4.260846 | 0.852519 | 0.962868 | 1.036958 | 0.834552 | 0.955402 | 1.04162 | 0.981662 | 0.98754 |
| 9.418981 | 0.694528 | 0.962868 | -1.12004 | 0.522153 | 0.943244 | -1.20343 | 0.803356 | 0.95342 |
| 4.753917 | 0.838494 | 0.962868 | -1.05182 | 0.94674 | 0.979621 | -1.01669 | 0.890976 | 0.976723 |
| 32.20862 | 0.804196 | 0.962868 | -1.06369 | 0.834371 | 0.955402 | 1.05345 | 0.648118 | 0.95342 |
| 20.04293 | 0.967438 | 0.993504 | -1.00867 | 0.86643 | 0.955402 | 1.036238 | 0.834466 | 0.95342 |
| 4.782053 | 0.762007 | 0.962868 | 1.062528 | 0.844608 | 0.955402 | -1.04001 | 0.61845 | 0.95342 |
| 8.333628 | 0.662091 | 0.962868 | 1.03813 | 0.001408 | 0.236463 | 1.350127 | 0.00436 | 0.732474 |
| 5.163999 | 0.10638 | 0.330959 | 1.180795 | 0.210274 | 0.815246 | 1.133367 | 0.702285 | 0.95342 |
| 76.79783 | 0.997253 | 0.997253 | -1.00064 | 0.728797 | 0.955402 | 1.067013 | 0.72622 | 0.95342 |
| 0.018904 | 0.073778 | 0.282583 | -2.08573 | 0.153344 | 0.815246 | -1.78749 | 0.699553 | 0.95342 |
| 0.796946 | 0.092226 | 0.303802 | -1.48001 | 0.200925 | 0.815246 | -1.34248 | 0.667822 | 0.95342 |
| 6.96964 | 0.079525 | 0.284259 | -1.50064 | 0.186755 | 0.815246 | -1.35242 | 0.644661 | 0.95342 |
| 2.814743 | 0.032631 | 0.271366 | -1.64491 | 0.096775 | 0.815246 | -1.46289 | 0.60048 | 0.95342 |
| 0.047397 | 0.001611 | 0.220027 | -1.42908 | 0.070333 | 0.815246 | -1.23122 | 0.11859 | 0.95342 |
| 0.174217 | 0.068613 | 0.274712 | -1.86628 | 0.124105 | 0.815246 | -1.68592 | 0.759992 | 0.95342 |
| 0.06418 | 0.117811 | 0.35986 | -2.09694 | 0.244697 | 0.822183 | -1.72509 | 0.673808 | 0.95342 |
| 0.084552 | 0.014236 | 0.271366 | -2.03007 | 0.244337 | 0.822183 | -1.38004 | 0.16522 | 0.95342 |
| 0.845139 | 0.062996 | 0.271366 | -2.26013 | 0.158598 | 0.815246 | -1.84044 | 0.629518 | 0.95342 |
| 0.367455 | 0.061066 | 0.271366 | -2.12811 | 0.189948 | 0.815246 | -1.68182 | 0.547985 | 0.95342 |
| 0.541663 | 0.015977 | 0.271366 | -1.69409 | 0.224657 | 0.815246 | -1.29074 | 0.196533 | 0.95342 |
| 0.264838 | 0.051003 | 0.271366 | -1.41256 | 0.444783 | 0.934464 | -1.14032 | 0.216724 | 0.95342 |
| 2.868606 | 0.032424 | 0.271366 | -1.97784 | 0.183179 | 0.815246 | -1.51212 | 0.383075 | 0.95342 |
| 8.974674 | 0.090661 | 0.303802 | -1.23256 | 0.395548 | 0.934464 | -1.11451 | 0.381319 | 0.95342 |
| 7.685715 | 0.03619 | 0.271366 | -1.48669 | 0.128794 | 0.815246 | -1.34617 | 0.530356 | 0.95342 |
| 2.009595 | 0.027106 | 0.271366 | -1.49819 | 0.266693 | 0.8768 | -1.21707 | 0.240615 | 0.95342 |
| 2.523571 | 0.00897 | 0.271366 | -1.93534 | 0.025324 | 0.811574 | -1.78688 | 0.659191 | 0.95342 |
| 1.17696 | 0.022159 | 0.271366 | -1.36424 | 0.333128 | 0.90267 | -1.14811 | 0.161752 | 0.95342 |
| 0.176549 | 0.083207 | 0.291224 | -1.92814 | 0.165289 | 0.815246 | -1.68316 | 0.712833 | 0.95342 |
| 0.082876 | 0.002619 | 0.220027 | -1.98581 | 0.020426 | 0.811574 | -1.66609 | 0.405142 | 0.95342 |
| 0.291603 | 0.038722 | 0.271366 | -2.08366 | 0.15863 | 0.815246 | -1.63263 | 0.477044 | 0.95342 |
| 0.05125 | 0.105592 | 0.330959 | -1.69817 | 0.301262 | 0.891111 | -1.39583 | 0.540802 | 0.95342 |
| 0.008577 | 0.129419 | 0.388256 | -1.61317 | 0.318254 | 0.891111 | -1.3648 | 0.589204 | 0.95342 |
| 0.173503 | 0.068678 | 0.274712 | -1.78151 | 0.238353 | 0.822183 | -1.44405 | 0.496747 | 0.95342 |
| 0.012534 | 0.208774 | 0.598882 | -1.36152 | 0.43488 | 0.934464 | -1.20928 | 0.624917 | 0.95342 |
| 0.936139 | 0.077374 | 0.282583 | -2.16673 | 0.172174 | 0.815246 | -1.80552 | 0.668728 | 0.95342 |
| 0.384172 | 0.042639 | 0.271366 | -2.29753 | 0.17565 | 0.815246 | -1.72316 | 0.468726 | 0.95342 |
| 0.011389 | 0.061262 | 0.271366 | -2.29824 | 0.082727 | 0.815246 | -2.15533 | 0.881496 | 0.976723 |
| 0.005834 | 0.074274 | 0.282583 | -1.92985 | 0.27139 | 0.8768 | -1.48859 | 0.470141 | 0.95342 |
| 0.085295 | 0.044551 | 0.271366 | -2.08054 | 0.211025 | 0.815246 | -1.56186 | 0.417335 | 0.95342 |
| 0.409443 | 0.019572 | 0.271366 | -1.6931 | 0.164494 | 0.815246 | -1.3546 | 0.303102 | 0.95342 |

| | | | | | | | | |
|---|---|---|---|---|---|---|---|---|
| 7.055205 | 0.014323 | 0.271366 | -1.68116 | 0.084374 | 0.815246 | -1.46672 | 0.41792 | 0.95342 |
| 0.012244 | 0.665606 | 0.962868 | -1.07653 | 0.602342 | 0.955402 | -1.09305 | 0.928769 | 0.979438 |
| 0.779033 | 0.006978 | 0.271366 | -1.83367 | 0.030098 | 0.811574 | -1.60926 | 0.535781 | 0.95342 |
| 0.045683 | 0.033541 | 0.271366 | -1.28009 | 0.358709 | 0.915711 | -1.11692 | 0.203426 | 0.95342 |
| 0.00479 | 0.057219 | 0.271366 | -1.43713 | 0.029802 | 0.811574 | -1.5046 | 0.761812 | 0.95342 |
| 0.030574 | 0.041844 | 0.271366 | -1.44045 | 0.48051 | 0.934464 | -1.13015 | 0.167283 | 0.95342 |
| 0.324462 | 0.038676 | 0.271366 | -1.76833 | 0.124789 | 0.815246 | -1.5155 | 0.561752 | 0.95342 |
| 0.048932 | 0.198606 | 0.585364 | -1.5771 | 0.346656 | 0.915711 | -1.39263 | 0.721901 | 0.95342 |
| 1.297296 | 0.030411 | 0.271366 | -1.91879 | 0.175911 | 0.815246 | -1.55951 | 0.380188 | 0.95342 |
| 50.61941 | 0.048926 | 0.271366 | -1.56202 | 0.161088 | 0.815246 | -1.36593 | 0.540746 | 0.95342 |
| 0.01486 | 0.055795 | 0.271366 | -1.89657 | 0.41059 | 0.934464 | -1.30725 | 0.255766 | 0.95342 |
| 0.009339 | 0.02731 | 0.271366 | -1.59163 | 0.286345 | 0.878057 | -1.24223 | 0.224568 | 0.95342 |
| 0.902791 | 0.058431 | 0.271366 | -1.37123 | 0.704378 | 0.955402 | -1.06325 | 0.123116 | 0.95342 |
| 0.007733 | 0.291324 | 0.776371 | -1.21472 | 0.313496 | 0.891111 | -1.20397 | 0.961164 | 0.979438 |
| 0.047598 | 0.04857 | 0.271366 | -1.25317 | 0.920861 | 0.979621 | 1.011031 | 0.03927 | 0.95342 |
| 0.016382 | 0.035279 | 0.271366 | -1.81452 | 0.038524 | 0.811574 | -1.79471 | 0.967778 | 0.979438 |
| 0.083058 | 0.04241 | 0.271366 | -2.1008 | 0.089957 | 0.815246 | -1.84617 | 0.714068 | 0.95342 |
| 2.356353 | 0.051099 | 0.271366 | -2.03917 | 0.173467 | 0.815246 | -1.62961 | 0.526577 | 0.95342 |
| 0.035283 | 0.062415 | 0.271366 | -1.55969 | 0.499591 | 0.934464 | -1.16959 | 0.219224 | 0.95342 |
| 0.337473 | 0.052498 | 0.271366 | -2.06866 | 0.068708 | 0.815246 | -1.97308 | 0.896094 | 0.976723 |
| 0.081142 | 0.076202 | 0.282583 | -1.75887 | 0.087137 | 0.815246 | -1.72235 | 0.94593 | 0.979438 |
| 0.129621 | 0.059597 | 0.271366 | -1.95228 | 0.088753 | 0.815246 | -1.82362 | 0.842859 | 0.95342 |
| 9.035435 | 0.322965 | 0.797913 | 1.08191 | 0.221963 | 0.815246 | 1.102676 | 0.80979 | 0.95342 |
| 0.552578 | 0.282026 | 0.764199 | -1.23793 | 0.386542 | 0.934464 | -1.1908 | 0.829574 | 0.95342 |
| 0.016873 | 0.345721 | 0.818043 | 1.096239 | 0.329691 | 0.90267 | -1.0997 | 0.061117 | 0.95342 |
| 23.65326 | 0.460547 | 0.932192 | 1.046106 | 0.467229 | 0.934464 | 1.0454 | 0.991134 | 0.991134 |
| 0.056828 | 0.822158 | 0.962868 | -1.054 | 0.857114 | 0.955402 | -1.04302 | 0.96429 | 0.979438 |
| 0.444816 | 0.05719 | 0.271366 | 1.466354 | 0.503447 | 0.934464 | 1.139745 | 0.202273 | 0.95342 |
| 0.25057 | 0.06695 | 0.274712 | 1.425968 | 0.643781 | 0.955402 | -1.09093 | 0.024715 | 0.95342 |
| 0.104323 | 0.878191 | 0.962868 | -1.02142 | 0.104793 | 0.815246 | -1.25828 | 0.139293 | 0.95342 |
| 0.139553 | 0.77718 | 0.962868 | -1.04977 | 0.472709 | 0.934464 | -1.13172 | 0.66176 | 0.95342 |
| 0.705867 | 0.982148 | 0.993981 | 1.002851 | 0.783229 | 0.955402 | -1.03565 | 0.766131 | 0.95342 |
| 0.658039 | 0.697643 | 0.962868 | 1.074656 | 0.219642 | 0.815246 | -1.25899 | 0.110507 | 0.95342 |
| 5.178701 | 0.334898 | 0.815403 | 1.069139 | 0.038646 | 0.811574 | 1.159341 | 0.244507 | 0.95342 |
| 5.432997 | 0.659415 | 0.962868 | 1.036165 | 0.870074 | 0.955402 | -1.01293 | 0.546436 | 0.95342 |
| 0.001605 | 0.40375 | 0.874424 | 1.128782 | 0.082359 | 0.815246 | -1.29363 | 0.013102 | 0.95342 |
| 3.790727 | 0.769851 | 0.962868 | 1.02601 | 0.819009 | 0.955402 | -1.02028 | 0.602754 | 0.95342 |
| 0.021611 | 0.369706 | 0.850831 | 1.202372 | 0.427194 | 0.934464 | 1.176886 | 0.91635 | 0.976723 |
| 5.101566 | 0.887392 | 0.962868 | 1.015445 | 0.572413 | 0.955402 | 1.064733 | 0.671702 | 0.95342 |
| 1.766427 | 0.55319 | 0.962868 | -1.02384 | 0.407869 | 0.934464 | -1.03354 | 0.812028 | 0.95342 |
| 0.788592 | 0.345693 | 0.818043 | -1.05605 | 0.465605 | 0.934464 | -1.04323 | 0.827936 | 0.95342 |
| 1.602631 | 0.672448 | 0.962868 | 1.016504 | 0.692107 | 0.955402 | -1.01545 | 0.415012 | 0.95342 |
| 1.688194 | 0.846285 | 0.962868 | 1.007155 | 0.926941 | 0.979621 | 1.003377 | 0.918585 | 0.976723 |
| 2.353416 | 0.655248 | 0.962868 | 1.019109 | 0.208198 | 0.815246 | 1.055521 | 0.409701 | 0.95342 |
| 2.294714 | 0.857766 | 0.962868 | -1.00605 | 0.993386 | 0.993386 | 1.000279 | 0.851268 | 0.95342 |
| 1.271325 | 0.917157 | 0.975205 | 1.003725 | 0.768888 | 0.955402 | 1.010567 | 0.849331 | 0.95342 |
| 2.193213 | 0.798702 | 0.962868 | -1.00869 | 0.97379 | 0.979621 | -1.00111 | 0.824141 | 0.95342 |
| 1.249458 | 0.490961 | 0.938136 | -1.03413 | 0.963231 | 0.979621 | -1.00224 | 0.520073 | 0.95342 |
| 1.730121 | 0.736426 | 0.962868 | -1.01264 | 0.821812 | 0.955402 | -1.00843 | 0.911137 | 0.976723 |

| | | | | | | | | |
|---|---|---|---|---|---|---|---|---|
| 5.587237 | 0.481035 | 0.938136 | 1.041448 | 0.51173 | 0.934464 | 1.038518 | 0.960834 | 0.979438 |
| 1.28214 | 0.874116 | 0.962868 | 1.006597 | 0.803418 | 0.955402 | -1.01039 | 0.684065 | 0.95342 |
| 0.430967 | 0.315491 | 0.797913 | 1.046063 | 0.306487 | 0.891111 | -1.04697 | 0.048423 | 0.95342 |
| 1.770491 | 0.956072 | 0.993504 | -1.00181 | 0.666508 | 0.955402 | -1.01431 | 0.706839 | 0.95342 |
| 1.637523 | 0.888785 | 0.962868 | -1.00525 | 0.684055 | 0.955402 | -1.01538 | 0.789079 | 0.95342 |
| 1.181951 | 0.857948 | 0.962868 | 1.009672 | 0.655595 | 0.955402 | 1.024313 | 0.789011 | 0.95342 |
| 1.866826 | 0.247121 | 0.691939 | 1.076114 | 0.533777 | 0.953985 | -1.0377 | 0.080708 | 0.95342 |
| 3.089732 | 0.441805 | 0.916337 | 1.029062 | 0.228075 | 0.815246 | -1.04629 | 0.05387 | 0.95342 |
| 1.176731 | 0.023118 | 0.271366 | 1.200664 | 0.43233 | 0.934464 | 1.062499 | 0.119367 | 0.95342 |
| 0.811831 | 0.320437 | 0.797913 | 1.115777 | 0.184719 | 0.815246 | -1.1395 | 0.024836 | 0.95342 |
| 1.266634 | 0.800355 | 0.962868 | 1.023308 | 0.217674 | 0.815246 | -1.11251 | 0.140624 | 0.95342 |
| 2.003096 | 0.76934 | 0.962868 | -1.01139 | 0.870098 | 0.955402 | -1.00633 | 0.896666 | 0.976723 |
| 1.821767 | 0.959251 | 0.993504 | 1.001769 | 0.490967 | 0.934464 | -1.02422 | 0.459808 | 0.95342 |
| 2.099398 | 0.758922 | 0.962868 | 1.015855 | 0.947629 | 0.979621 | -1.00337 | 0.709612 | 0.95342 |
| 0.311226 | 0.405983 | 0.874424 | -1.06912 | 0.814838 | 0.955402 | 1.018903 | 0.289307 | 0.95342 |
| 1.380735 | 0.975763 | 0.993504 | 1.001735 | 0.194734 | 0.815246 | -1.07801 | 0.18494 | 0.95342 |
| 1.579851 | 0.560184 | 0.962868 | -1.02214 | 0.494283 | 0.934464 | 1.026059 | 0.210296 | 0.95342 |
| 41.83064 | 0.854337 | 0.962868 | 1.006229 | 0.832151 | 0.955402 | -1.0072 | 0.692758 | 0.95342 |
| 1.19923 | 0.210322 | 0.598882 | -1.05554 | 0.584211 | 0.955402 | 1.023617 | 0.077036 | 0.95342 |
| 0.511931 | 0.868945 | 0.962868 | 1.0145 | 0.552388 | 0.955402 | 1.05337 | 0.666957 | 0.95342 |
| 0.070936 | 0.028355 | 0.271366 | 1.403145 | 0.5437 | 0.955402 | 1.094243 | 0.100753 | 0.95342 |
| 1.510493 | 0.707947 | 0.962868 | -1.01791 | 0.509558 | 0.934464 | -1.03182 | 0.774244 | 0.95342 |
| 3.246798 | 0.485478 | 0.938136 | -1.02644 | 0.79999 | 0.955402 | -1.00957 | 0.655386 | 0.95342 |
| 0.790831 | 0.660046 | 0.962868 | -1.03177 | 0.843455 | 0.955402 | -1.01425 | 0.808048 | 0.95342 |
| 1.52E-05 | 0.102828 | 0.330959 | -2.0572 | 0.745628 | 0.955402 | 1.150435 | 0.053703 | 0.95342 |
| 9.32E-05 | 0.397064 | 0.874424 | -1.46525 | 0.669425 | 0.955402 | 1.211254 | 0.207079 | 0.95342 |
| 0.00072 | 0.522421 | 0.962868 | -1.21911 | 0.851471 | 0.955402 | -1.0595 | 0.649939 | 0.95342 |
| 0.001561 | 0.880058 | 0.962868 | -1.02908 | 0.50437 | 0.934464 | 1.14831 | 0.414324 | 0.95342 |
| 0.00126 | 0.430751 | 0.904577 | -1.332 | 0.690067 | 0.955402 | 1.200945 | 0.239284 | 0.95342 |
| 0.000173 | 0.723001 | 0.962868 | -1.1243 | 0.76084 | 0.955402 | 1.105811 | 0.511238 | 0.95342 |
| 0.001755 | 0.488161 | 0.938136 | -1.26081 | 0.880728 | 0.960794 | -1.05122 | 0.585945 | 0.95342 |
| 0.001267 | 0.55233 | 0.962868 | -1.1904 | 0.816557 | 0.955402 | -1.0702 | 0.716116 | 0.95342 |
| 9.19E-05 | 0.422469 | 0.898416 | -1.29426 | 0.588998 | 0.955402 | -1.18909 | 0.791044 | 0.95342 |
| 0.00309 | 0.508833 | 0.960493 | -1.26937 | 0.786544 | 0.955402 | -1.10235 | 0.695241 | 0.95342 |
| 0.00119 | 0.745134 | 0.962868 | -1.09058 | 0.957626 | 0.979621 | -1.01426 | 0.785578 | 0.95342 |
| 0.000355 | 0.87022 | 0.962868 | -1.02557 | 0.706522 | 0.955402 | 1.059983 | 0.589879 | 0.95342 |
| 0.000367 | 0.715613 | 0.962868 | -1.05752 | 0.844434 | 0.955402 | 1.030555 | 0.575841 | 0.95342 |
| 0.000136 | 0.62885 | 0.962868 | 1.054844 | 0.919848 | 0.979621 | 1.01092 | 0.701552 | 0.95342 |
| 0.002243 | 0.606868 | 0.962868 | -1.13305 | 0.819631 | 0.955402 | 1.056801 | 0.459165 | 0.95342 |
| 0.000939 | 0.457644 | 0.932192 | -1.1401 | 0.650022 | 0.955402 | -1.08312 | 0.77055 | 0.95342 |
| 0.002459 | 0.621966 | 0.962868 | -1.15403 | 0.86999 | 0.955402 | 1.048607 | 0.51227 | 0.95342 |
| 0.00195 | 0.840403 | 0.962868 | -1.03786 | 0.627829 | 0.955402 | 1.09376 | 0.49375 | 0.95342 |
| 0.003086 | 0.666878 | 0.962868 | -1.13322 | 0.755798 | 0.955402 | 1.094489 | 0.460048 | 0.95342 |
| 0.002516 | 0.76624 | 0.962868 | -1.08833 | 0.629648 | 0.955402 | 1.147379 | 0.437446 | 0.95342 |
| 0.001643 | 0.896098 | 0.962868 | -1.03666 | 0.56939 | 0.955402 | 1.170365 | 0.485089 | 0.95342 |
| 0.002092 | 0.732852 | 0.962868 | -1.10069 | 0.568624 | 0.955402 | 1.174134 | 0.364586 | 0.95342 |
| 0.00057 | 0.725601 | 0.962868 | -1.06737 | 0.951632 | 0.979621 | 1.011319 | 0.680777 | 0.95342 |
| 0.001285 | 0.871716 | 0.962868 | -1.05399 | 0.774044 | 0.955402 | 1.098046 | 0.65415 | 0.95342 |
| 0.000732 | 0.720596 | 0.962868 | 1.077812 | 0.317196 | 0.891111 | 1.235194 | 0.516459 | 0.95342 |

| | | | | | | | | |
|---|---|---|---|---|---|---|---|---|
| 0.000641 | 0.820213 | 0.962868 | 1.054065 | 0.466009 | 0.934464 | 1.184863 | 0.614281 | 0.95342 |
| 0.000109 | 0.566049 | 0.962868 | 1.110304 | 0.287459 | 0.878057 | 1.215754 | 0.618488 | 0.95342 |
| 3.41E-05 | 0.996613 | 0.997253 | -1.00108 | 0.942231 | 0.979621 | -1.01862 | 0.945609 | 0.979438 |
| 1.22E-05 | 0.803379 | 0.962868 | 1.044124 | 0.748899 | 0.955402 | -1.0571 | 0.570142 | 0.95342 |
| 7.12E-05 | 0.809712 | 0.962868 | -1.12456 | 0.772904 | 0.955402 | -1.1511 | 0.961819 | 0.979438 |
| 0.001008 | 0.899823 | 0.962868 | 1.023867 | 0.370645 | 0.915711 | 1.184054 | 0.440279 | 0.95342 |
| 0.000417 | 0.853265 | 0.962868 | 1.036195 | 0.432123 | 0.934464 | 1.163949 | 0.546526 | 0.95342 |
| 0.03324 | 0.765636 | 0.962868 | -1.07835 | 0.731598 | 0.955402 | 1.090702 | 0.522616 | 0.95342 |
| 0.000497 | 0.791735 | 0.962868 | 1.089415 | 0.969338 | 0.979621 | -1.01254 | 0.762337 | 0.95342 |
| 0.000369 | 0.597302 | 0.962868 | -1.19647 | 0.865669 | 0.955402 | -1.05898 | 0.718852 | 0.95342 |
| 2.45E-05 | 0.317903 | 0.797913 | 1.259476 | 0.619395 | 0.955402 | 1.120694 | 0.610852 | 0.95342 |
| 0.000236 | 0.899459 | 0.962868 | 1.031578 | 0.748335 | 0.955402 | 1.082215 | 0.845624 | 0.95342 |
| 0.000601 | 0.975119 | 0.993504 | -1.0055 | 0.279007 | 0.878057 | 1.212062 | 0.265821 | 0.95342 |
| 1.03E-05 | 0.397431 | 0.874424 | 1.161424 | 0.626428 | 0.955402 | 1.089486 | 0.716234 | 0.95342 |
| 8.59E-05 | 0.491404 | 0.938136 | -1.23921 | 0.858341 | 0.955402 | 1.056976 | 0.38775 | 0.95342 |
| 2.41E-05 | 0.694094 | 0.962868 | 1.07452 | 0.942714 | 0.979621 | -1.0132 | 0.642068 | 0.95342 |
| 0.000421 | 0.843931 | 0.962868 | 1.040927 | 0.429483 | 0.934464 | 1.175664 | 0.55137 | 0.95342 |
| 0.000158 | 0.86669 | 0.962868 | 1.027828 | 0.223474 | 0.815246 | 1.22348 | 0.29132 | 0.95342 |
| 2.04E-06 | 0.738184 | 0.962868 | 1.126069 | 0.818075 | 0.955402 | 1.08509 | 0.916808 | 0.976723 |
| 9.5E-06 | 0.29576 | 0.776371 | 1.148379 | 0.916094 | 0.979621 | -1.01289 | 0.251155 | 0.95342 |
| 0.00014 | 0.391566 | 0.874424 | 1.154386 | 0.037111 | 0.811574 | 1.434907 | 0.197995 | 0.95342 |
| 0.000134 | 0.743133 | 0.962868 | 1.043122 | 0.427942 | 0.934464 | 1.108059 | 0.639574 | 0.95342 |
| 0.001192 | 0.940705 | 0.993504 | -1.01522 | 0.646795 | 0.955402 | 1.097712 | 0.594649 | 0.95342 |

| Control - G | Control - Ve | Control - Ve | Control - Ve | JAK2A - GH | JAK2A - GH | JAK2A - GH | JAK2A - GH | JAK2A - GH |
|---|---|---|---|---|---|---|---|---|
| 1.114394 | 0.882358 | 0.986494 | 1.03721 | 0.47609 | 0.707815 | -1.18251 | 0.473088 | 0.998377 |
| -1.10058 | 0.218275 | 0.986494 | -1.25373 | 0.091641 | 0.301875 | -1.34653 | 0.176142 | 0.998377 |
| -1.02648 | 0.758564 | 0.986494 | 1.019113 | 0.799412 | 0.914371 | -1.01405 | 0.594188 | 0.998377 |
| 1.018714 | 0.767671 | 0.986494 | 1.016998 | 0.425512 | 0.668094 | -1.04332 | 0.187874 | 0.998377 |
| -1.31083 | 0.235958 | 0.986494 | -1.22086 | 0.505049 | 0.717734 | 1.111321 | 0.517614 | 0.998377 |
| 1.248232 | 0.390332 | 0.986494 | 1.201126 | 0.475942 | 0.707815 | -1.15485 | 0.957805 | 0.998377 |
| -1.01457 | 0.793079 | 0.986494 | 1.016918 | 0.642696 | 0.811826 | -1.02759 | 0.345117 | 0.998377 |
| 1.004496 | 0.531316 | 0.986494 | -1.13043 | 0.010189 | 0.120731 | -1.65798 | 0.832778 | 0.998377 |
| -1.07446 | 0.447821 | 0.986494 | -1.24601 | 0.878515 | 0.960733 | 1.042694 | 0.403374 | 0.998377 |
| 1.034554 | 0.610581 | 0.986494 | -1.13483 | 0.004508 | 0.076269 | -2.05358 | 0.747365 | 0.998377 |
| 1.120549 | 0.931404 | 0.986494 | -1.02166 | 0.012831 | 0.121829 | -1.86304 | 0.85597 | 0.998377 |
| 1.045227 | 0.957134 | 0.986494 | 1.011438 | 0.013893 | 0.121829 | -1.68564 | 0.844576 | 0.998377 |
| -1.10504 | 0.175849 | 0.908746 | -1.3171 | 0.250237 | 0.502126 | -1.24721 | 0.950451 | 0.998377 |
| 1.300538 | 0.178504 | 0.908746 | 1.12402 | 0.321808 | 0.563163 | 1.084437 | 0.481694 | 0.998377 |
| -1.04185 | 0.935589 | 0.986494 | -1.00855 | 0.653654 | 0.813437 | -1.04111 | 0.922489 | 0.998377 |
| 1.0677 | 0.842364 | 0.986494 | -1.03788 | 0.024882 | 0.144142 | -1.51668 | 0.782013 | 0.998377 |
| 1.166847 | 0.702453 | 0.986494 | -1.16502 | 0.853209 | 0.960733 | -1.07263 | 0.853329 | 0.998377 |
| 1.102441 | 0.684338 | 0.986494 | -1.09678 | 0.247198 | 0.502126 | -1.28676 | 0.958745 | 0.998377 |
| 1.109599 | 0.84985 | 0.986494 | -1.04354 | 0.216768 | 0.477919 | -1.30658 | 0.982434 | 0.998377 |
| 1.124425 | 0.81471 | 0.986494 | -1.05377 | 0.109739 | 0.335202 | -1.41471 | 0.956546 | 0.998377 |
| 1.160706 | 0.703035 | 0.986494 | 1.038442 | 0.769375 | 0.903881 | -1.03575 | 0.867443 | 0.998377 |
| 1.106977 | 0.401937 | 0.986494 | -1.32365 | 0.978641 | 0.984501 | 1.008479 | 0.912887 | 0.998377 |
| 1.215552 | 0.649345 | 0.986494 | -1.23479 | 0.64911 | 0.813437 | 1.221678 | 0.883298 | 0.998377 |
| 1.471027 | 0.642468 | 0.986494 | -1.13555 | 0.800075 | 0.914371 | 1.067902 | 0.606347 | 0.998377 |
| 1.228037 | 0.666789 | 0.986494 | -1.20113 | 0.880672 | 0.960733 | -1.0624 | 0.955192 | 0.998377 |
| 1.26536 | 0.783618 | 0.986494 | -1.11327 | 0.694296 | 0.851399 | -1.15696 | 0.991344 | 0.998377 |
| 1.312497 | 0.708579 | 0.986494 | -1.0807 | 0.262988 | 0.502126 | -1.2495 | 0.633472 | 0.998377 |
| 1.238744 | 0.948124 | 0.986494 | -1.01118 | 0.827313 | 0.939112 | -1.03602 | 0.742087 | 0.998377 |
| 1.307992 | 0.499414 | 0.986494 | -1.23039 | 0.440296 | 0.682203 | -1.25234 | 0.659855 | 0.998377 |
| 1.10592 | 0.849811 | 0.986494 | -1.02329 | 0.198934 | 0.451633 | -1.19855 | 0.805332 | 0.998377 |
| 1.104391 | 0.706761 | 0.986494 | -1.06661 | 0.333774 | 0.578083 | -1.26004 | 0.907061 | 0.998377 |
| 1.230982 | 0.272522 | 0.986494 | -1.21411 | 0.263018 | 0.502126 | -1.20663 | 0.566193 | 0.998377 |
| 1.083084 | 0.378882 | 0.986494 | -1.19997 | 0.507128 | 0.717734 | -1.26952 | 0.99188 | 0.998377 |
| 1.188251 | 0.301258 | 0.986494 | -1.16003 | 0.974222 | 0.984501 | -1.00467 | 0.166922 | 0.998377 |
| 1.145546 | 0.61442 | 0.986494 | -1.20465 | 0.184469 | 0.437411 | -1.6028 | 0.882207 | 0.998377 |
| 1.191898 | 0.533093 | 0.986494 | -1.14005 | 0.965954 | 0.984501 | 1.008522 | 0.550241 | 0.998377 |
| 1.276262 | 0.678797 | 0.986494 | -1.15215 | 0.5383 | 0.717734 | -1.22146 | 0.825379 | 0.998377 |
| 1.216603 | 0.878394 | 0.986494 | -1.05011 | 0.716159 | 0.85939 | -1.11665 | 0.710257 | 0.998377 |
| 1.181984 | 0.930063 | 0.986494 | -1.02748 | 0.228261 | 0.485416 | -1.42999 | 0.276705 | 0.998377 |
| 1.233688 | 0.676971 | 0.986494 | -1.137 | 0.62644 | 0.797288 | -1.15304 | 0.810701 | 0.998377 |
| 1.125891 | 0.457834 | 0.986494 | -1.19791 | 0.529742 | 0.717734 | 1.155801 | 0.228188 | 0.998377 |
| 1.200055 | 0.68586 | 0.986494 | -1.1881 | 0.71015 | 0.85831 | -1.16208 | 0.868332 | 0.998377 |
| 1.333325 | 0.719576 | 0.986494 | -1.15262 | 0.724829 | 0.863626 | -1.14123 | 0.846373 | 0.998377 |
| 1.066307 | 0.502022 | 0.986494 | -1.33675 | 0.97784 | 0.984501 | 1.011411 | 0.904753 | 0.998377 |
| 1.296426 | 0.675402 | 0.986494 | -1.16189 | 0.534054 | 0.717734 | -1.23588 | 0.998377 | 0.998377 |
| 1.332092 | 0.544674 | 0.986494 | -1.23813 | 0.920367 | 0.984501 | -1.0339 | 0.594945 | 0.998377 |
| 1.249887 | 0.651617 | 0.986494 | -1.1019 | 0.319304 | 0.563163 | -1.22692 | 0.677101 | 0.998377 |

| | | | | | | | | |
|---|---|---|---|---|---|---|---|---|
| 1.146207 | 0.60262 | 0.986494 | -1.10332 | 0.410048 | 0.668094 | -1.26094 | 0.935845 | 0.998377 |
| -1.01534 | 0.375671 | 0.986494 | -1.16414 | 0.312694 | 0.558858 | -1.179 | 0.301345 | 0.998377 |
| 1.139447 | 0.682271 | 0.986494 | -1.08995 | 0.994023 | 0.994023 | 1.001494 | 0.554507 | 0.998377 |
| 1.146089 | 0.561292 | 0.986494 | 1.065953 | 0.281257 | 0.518085 | -1.15021 | 0.902782 | 0.998377 |
| -1.04694 | 0.184354 | 0.910923 | -1.24903 | 0.246512 | 0.502126 | 1.247492 | 0.783668 | 0.998377 |
| 1.274565 | 0.54467 | 0.986494 | 1.110636 | 0.238528 | 0.500908 | -1.21593 | 0.955028 | 0.998377 |
| 1.166824 | 0.723571 | 0.986494 | -1.09841 | 0.946607 | 0.984501 | -1.01698 | 0.720829 | 0.998377 |
| 1.132463 | 0.927883 | 0.986494 | 1.032104 | 0.08149 | 0.285216 | -1.80976 | 0.647976 | 0.998377 |
| 1.230377 | 0.594253 | 0.986494 | -1.16174 | 0.785165 | 0.909936 | -1.11758 | 0.756844 | 0.998377 |
| 1.143561 | 0.676111 | 0.986494 | -1.09577 | 0.260385 | 0.502126 | -1.26626 | 0.848699 | 0.998377 |
| 1.450804 | 0.853146 | 0.986494 | -1.06175 | 0.705405 | 0.85831 | 1.123231 | 0.562247 | 0.998377 |
| 1.281264 | 0.726064 | 0.986494 | 1.073181 | 0.058923 | 0.235692 | 1.451769 | 0.929217 | 0.998377 |
| 1.289653 | 0.582604 | 0.986494 | -1.09305 | 0.970137 | 0.984501 | 1.00575 | 0.545623 | 0.998377 |
| 1.008925 | 0.989598 | 0.999821 | 1.002382 | 0.767825 | 0.903881 | -1.05248 | 0.409567 | 0.998377 |
| 1.266992 | 0.366953 | 0.986494 | 1.10556 | 0.220369 | 0.477919 | -1.13898 | 0.497616 | 0.998377 |
| 1.011038 | 0.398323 | 0.986494 | -1.25988 | 0.507158 | 0.717734 | -1.18729 | 0.973373 | 0.998377 |
| 1.137923 | 0.516911 | 0.986494 | -1.25752 | 0.254609 | 0.502126 | -1.46987 | 0.94179 | 0.998377 |
| 1.25132 | 0.738313 | 0.986494 | -1.12522 | 0.528246 | 0.717734 | -1.23594 | 0.812262 | 0.998377 |
| 1.333534 | 0.853293 | 0.986494 | -1.04367 | 0.785361 | 0.909936 | 1.061564 | 0.587693 | 0.998377 |
| 1.048444 | 0.503399 | 0.986494 | -1.27551 | 0.274588 | 0.512565 | -1.46155 | 0.95513 | 0.998377 |
| 1.021208 | 0.796303 | 0.986494 | -1.0832 | 0.186432 | 0.437411 | -1.48298 | 0.669773 | 0.998377 |
| 1.070553 | 0.472078 | 0.986494 | -1.28194 | 0.160755 | 0.412116 | -1.59313 | 0.753842 | 0.998377 |
| 1.019195 | 0.17403 | 0.908746 | 1.115322 | 0.892224 | 0.967055 | 1.010201 | 0.603637 | 0.998377 |
| 1.039576 | 0.707655 | 0.986494 | -1.07412 | 0.023207 | 0.144142 | 1.494187 | 0.668532 | 0.998377 |
| -1.20553 | 0.521908 | 0.986494 | -1.06411 | 0.92981 | 0.984501 | 1.008111 | 0.406629 | 0.998377 |
| -1.00068 | 0.423956 | 0.986494 | 1.050095 | 0.46071 | 0.697291 | 1.043673 | 0.70339 | 0.998377 |
| 1.010525 | 0.581128 | 0.986494 | -1.13813 | 0.42006 | 0.668094 | -1.19715 | 0.597408 | 0.998377 |
| -1.28656 | 0.438909 | 0.986494 | -1.16362 | 0.552786 | 0.731245 | -1.11627 | 0.945386 | 0.998377 |
| -1.55563 | 0.364522 | 0.986494 | -1.18721 | 0.12222 | 0.359133 | -1.32508 | 0.136134 | 0.998377 |
| -1.23189 | 0.531576 | 0.986494 | -1.09069 | 0.423058 | 0.668094 | 1.111497 | 0.958206 | 0.998377 |
| -1.07806 | 0.918535 | 0.986494 | 1.017699 | 0.53799 | 0.717734 | 1.105785 | 0.715666 | 0.998377 |
| -1.0386 | 0.937347 | 0.986494 | -1.01005 | 0.457047 | 0.697291 | -1.09441 | 0.496323 | 0.998377 |
| -1.35299 | 0.115144 | 0.690864 | -1.34752 | 0.560783 | 0.736028 | 1.107875 | 0.593158 | 0.998377 |
| 1.084369 | 0.270207 | 0.986494 | 1.079654 | 0.584532 | 0.761251 | 1.036403 | 0.576938 | 0.998377 |
| -1.04957 | 0.84848 | 0.986494 | 1.014916 | 0.359638 | 0.616523 | 1.069228 | 0.629009 | 0.998377 |
| -1.46023 | 0.938935 | 0.986494 | -1.01111 | 0.616099 | 0.790112 | 1.07115 | 0.635903 | 0.998377 |
| -1.04682 | 0.88581 | 0.986494 | -1.01268 | 0.187462 | 0.437411 | 1.117856 | 0.664544 | 0.998377 |
| -1.02166 | 0.732686 | 0.986494 | 1.072225 | 0.943201 | 0.984501 | -1.01388 | 0.536814 | 0.998377 |
| 1.048539 | 0.341514 | 0.986494 | 1.109657 | 0.97744 | 0.984501 | 1.002881 | 0.735829 | 0.998377 |
| -1.00947 | 0.000586 | 0.049247 | 1.164327 | 0.032769 | 0.164334 | 1.087254 | 0.178716 | 0.998377 |
| 1.012289 | 0.135161 | 0.783 | 1.086175 | 0.003576 | 0.076269 | 1.176321 | 0.561935 | 0.998377 |
| -1.03221 | 0.024541 | 0.242521 | 1.095328 | 0.39855 | 0.656435 | 1.031649 | 0.251318 | 0.998377 |
| -1.00377 | 0.010816 | 0.12114 | 1.104609 | 0.178518 | 0.437411 | 1.048858 | 0.363519 | 0.998377 |
| 1.035728 | 0.086403 | 0.580631 | 1.07739 | 0.520956 | 0.717734 | 1.026204 | 0.199296 | 0.998377 |
| 1.006329 | 0.010602 | 0.12114 | 1.095612 | 0.157501 | 0.412116 | 1.04701 | 0.273395 | 0.998377 |
| 1.006816 | 0.005116 | 0.095506 | 1.113627 | 0.262123 | 0.502126 | 1.039216 | 0.698239 | 0.998377 |
| 1.007565 | 0.007338 | 0.102728 | 1.102019 | 0.054897 | 0.22746 | 1.065941 | 0.247515 | 0.998377 |
| 1.031815 | 0.046969 | 0.415306 | 1.105079 | 0.123986 | 0.359133 | 1.075015 | 0.235987 | 0.998377 |
| 1.004168 | 0.002685 | 0.075188 | 1.129357 | 0.081284 | 0.285216 | 1.065423 | 0.168171 | 0.998377 |

| | | | | | | | | |
|---|---|---|---|---|---|---|---|---|
| -1.00282 | 0.076773 | 0.560775 | 1.110145 | 0.370505 | 0.628736 | 1.050327 | 0.162992 | 0.998377 |
| -1.01705 | 0.064012 | 0.512097 | 1.082526 | 0.101013 | 0.314697 | 1.068396 | 0.235731 | 0.998377 |
| -1.09519 | 0.002488 | 0.075188 | 1.157687 | 0.146741 | 0.412116 | 1.064375 | 0.436717 | 0.998377 |
| -1.01247 | 0.009699 | 0.12114 | 1.09477 | 0.490237 | 0.717734 | 1.021869 | 0.166428 | 0.998377 |
| -1.01007 | 0.001751 | 0.075188 | 1.13698 | 0.283713 | 0.518085 | 1.039227 | 0.26194 | 0.998377 |
| 1.0145 | 0.171585 | 0.908746 | 1.077629 | 0.869453 | 0.960733 | -1.00842 | 0.988038 | 0.998377 |
| -1.11668 | 0.10611 | 0.660239 | 1.096283 | 0.024597 | 0.144142 | 1.134859 | 0.526229 | 0.998377 |
| -1.07669 | 0.102039 | 0.659328 | 1.064037 | 0.070145 | 0.267825 | 1.067786 | 0.135521 | 0.998377 |
| -1.13004 | 0.057988 | 0.487101 | 1.162324 | 0.164356 | 0.412116 | 1.108585 | 0.416382 | 0.998377 |
| -1.27143 | 0.405059 | 0.986494 | 1.076106 | 0.388245 | 0.645992 | -1.09325 | 0.926763 | 0.998377 |
| -1.13844 | 0.610844 | 0.986494 | 1.04127 | 0.221891 | 0.477919 | -1.11824 | 0.970372 | 0.998377 |
| 1.005026 | 0.017869 | 0.187623 | 1.101033 | 0.122426 | 0.359133 | 1.059502 | 0.231581 | 0.998377 |
| -1.02603 | 0.028678 | 0.267662 | 1.082315 | 0.15089 | 0.412116 | 1.049215 | 0.194075 | 0.998377 |
| -1.01928 | 0.074342 | 0.560775 | 1.098658 | 0.600844 | 0.776475 | -1.02581 | 0.57466 | 0.998377 |
| 1.089325 | 0.000355 | 0.049247 | 1.379627 | 0.047867 | 0.206194 | 1.168239 | 0.261494 | 0.998377 |
| -1.07988 | 0.296643 | 0.986494 | 1.061977 | 0.537048 | 0.717734 | -1.03409 | 0.944272 | 0.998377 |
| 1.048778 | 0.003517 | 0.082258 | 1.12568 | 0.098166 | 0.314697 | 1.062142 | 0.198756 | 0.998377 |
| -1.01347 | 0.007029 | 0.102728 | 1.1024 | 0.163233 | 0.412116 | 1.04658 | 0.199107 | 0.998377 |
| 1.080471 | 0.005709 | 0.095917 | 1.134515 | 0.060497 | 0.236359 | 1.081381 | 0.820768 | 0.998377 |
| 1.038315 | 0.002219 | 0.075188 | 1.337971 | 0.661437 | 0.817069 | 1.036969 | 0.06354 | 0.998377 |
| -1.2823 | 0.871127 | 0.986494 | -1.02428 | 0.101153 | 0.314697 | 1.265672 | 0.934898 | 0.998377 |
| -1.01367 | 0.08068 | 0.56476 | 1.088702 | 0.875692 | 0.960733 | -1.00705 | 0.662543 | 0.998377 |
| 1.016711 | 0.003917 | 0.082258 | 1.116485 | 0.148987 | 0.412116 | 1.05265 | 0.37383 | 0.998377 |
| 1.017282 | 0.244048 | 0.986494 | 1.083852 | 0.027194 | 0.152284 | 1.158029 | 0.490872 | 0.998377 |
| 2.366677 | 0.849222 | 0.986494 | 1.085536 | 0.182351 | 0.437411 | 1.741783 | 0.583569 | 0.998377 |
| 1.774786 | 0.81228 | 0.986494 | 1.112364 | 0.965273 | 0.984501 | 1.018683 | 0.836431 | 0.998377 |
| 1.150642 | 0.979422 | 0.999821 | 1.007992 | 0.295512 | 0.533829 | -1.36251 | 0.874131 | 0.998377 |
| 1.181702 | 0.852416 | 0.986494 | -1.04294 | 0.388364 | 0.645992 | -1.18869 | 0.838905 | 0.998377 |
| 1.599656 | 0.988811 | 0.999821 | -1.00711 | 0.96042 | 0.984501 | -1.02012 | 0.853015 | 0.998377 |
| 1.243263 | 0.999821 | 0.999821 | -1.00007 | 0.506253 | 0.717734 | -1.23248 | 0.802817 | 0.998377 |
| 1.199372 | 0.953061 | 0.986494 | 1.019786 | 0.44262 | 0.682203 | -1.27601 | 0.982376 | 0.998377 |
| 1.112311 | 0.807065 | 0.986494 | 1.074047 | 0.220101 | 0.477919 | -1.41168 | 0.923286 | 0.998377 |
| 1.088454 | 0.852388 | 0.986494 | -1.06131 | 0.517981 | 0.717734 | -1.21751 | 0.81281 | 0.998377 |
| 1.15151 | 0.949203 | 0.986494 | 1.023175 | 0.417586 | 0.668094 | -1.32075 | 0.899462 | 0.998377 |
| 1.075247 | 0.804414 | 0.986494 | 1.06826 | 0.156481 | 0.412116 | -1.44032 | 0.787771 | 0.998377 |
| 1.087088 | 0.864962 | 0.986494 | 1.026631 | 0.085997 | 0.294848 | -1.29504 | 0.415724 | 0.998377 |
| 1.089834 | 0.702861 | 0.986494 | 1.060314 | 0.001944 | 0.065328 | -1.63709 | 0.40702 | 0.998377 |
| -1.04345 | 0.775959 | 0.986494 | -1.03189 | 0.271015 | 0.511579 | -1.12782 | 0.082792 | 0.998377 |
| 1.197404 | 0.797782 | 0.986494 | 1.064054 | 0.044374 | 0.199243 | -1.61504 | 0.967293 | 0.998377 |
| 1.052605 | 0.927443 | 0.986494 | -1.01612 | 0.028247 | 0.153078 | -1.46536 | 0.904231 | 0.998377 |
| 1.210128 | 0.791904 | 0.986494 | 1.079545 | 0.009568 | 0.120731 | -2.13459 | 0.935606 | 0.998377 |
| 1.135171 | 0.915046 | 0.986494 | 1.019876 | 0.029673 | 0.155783 | -1.48801 | 0.874295 | 0.998377 |
| 1.240295 | 0.808388 | 0.986494 | 1.072914 | 0.01897 | 0.12748 | -1.97222 | 0.920079 | 0.998377 |
| 1.248731 | 0.701067 | 0.986494 | 1.115563 | 0.016216 | 0.12748 | -1.98246 | 0.872743 | 0.998377 |
| 1.213277 | 0.714109 | 0.986494 | 1.10641 | 0.014503 | 0.121829 | -1.96518 | 0.873001 | 0.998377 |
| 1.292356 | 0.645141 | 0.986494 | 1.138313 | 0.033283 | 0.164334 | -1.80593 | 0.92388 | 0.998377 |
| 1.07945 | 0.791808 | 0.986494 | 1.050229 | 0.000973 | 0.063787 | -1.90178 | 0.818687 | 0.998377 |
| 1.157328 | 0.727009 | 0.986494 | 1.120503 | 0.01078 | 0.120731 | -2.30764 | 0.874924 | 0.998377 |
| 1.146019 | 0.762218 | 0.986494 | 1.065421 | 0.008133 | 0.113864 | -1.75167 | 0.997369 | 0.998377 |

| | | | | | | | | |
|---|---|---|---|---|---|---|---|---|
| 1.124089 | 0.720636 | 0.986494 | 1.086441 | 0.011533 | 0.121097 | -1.80153 | 0.924249 | 0.998377 |
| 1.094974 | 0.662133 | 0.986494 | 1.082823 | 0.075973 | 0.277466 | -1.37022 | 0.258267 | 0.998377 |
| -1.01752 | 0.78507 | 0.986494 | 1.071955 | 0.013108 | 0.121829 | -1.88561 | 0.560407 | 0.998377 |
| -1.10375 | 0.994581 | 0.999821 | 1.001178 | 0.018324 | 0.12748 | -1.50426 | 0.840492 | 0.998377 |
| -1.0236 | 0.680537 | 0.986494 | 1.222536 | 0.039158 | 0.182739 | -2.6949 | 0.849062 | 0.998377 |
| 1.156453 | 0.451896 | 0.986494 | 1.152177 | 0.004003 | 0.076269 | -1.73736 | 0.809562 | 0.998377 |
| 1.123291 | 0.664872 | 0.986494 | 1.08697 | 0.045067 | 0.199243 | -1.46094 | 0.926212 | 0.998377 |
| 1.176159 | 0.763451 | 0.986494 | 1.079134 | 0.021903 | 0.14153 | -1.78077 | 0.967422 | 0.998377 |
| -1.10307 | 0.691869 | 0.986494 | 1.137279 | 0.017497 | 0.12748 | -2.15898 | 0.908689 | 0.998377 |
| 1.129829 | 0.541585 | 0.986494 | 1.230519 | 0.055511 | 0.22746 | -1.8894 | 0.911595 | 0.998377 |
| -1.12384 | 0.887391 | 0.986494 | 1.03295 | 0.005643 | 0.08618 | -1.90648 | 0.922713 | 0.998377 |
| 1.049087 | 0.672736 | 0.986494 | 1.109702 | 0.017586 | 0.12748 | -1.79266 | 0.954037 | 0.998377 |
| 1.218725 | 0.684097 | 0.986494 | 1.074253 | 0.00176 | 0.065328 | -1.77083 | 0.745761 | 0.998377 |
| -1.06603 | 0.945218 | 0.986494 | -1.01215 | 0.00454 | 0.076269 | -1.66486 | 0.533172 | 0.998377 |
| 1.309817 | 0.521391 | 0.986494 | 1.221099 | 0.153613 | 0.412116 | -1.53419 | 0.632957 | 0.998377 |
| -1.0887 | 0.702165 | 0.986494 | -1.07237 | 0.034236 | 0.164334 | -1.46514 | 0.83109 | 0.998377 |
| 1.12944 | 0.427177 | 0.986494 | 1.176628 | 0.090661 | 0.301875 | -1.39876 | 0.870644 | 0.998377 |
| 1.190356 | 0.305703 | 0.986494 | 1.18417 | 0.073756 | 0.275357 | -1.33047 | 0.549316 | 0.998377 |
| -1.03777 | 0.916786 | 0.986494 | 1.037776 | 0.196882 | 0.451633 | -1.55437 | 0.903777 | 0.998377 |
| -1.16318 | 0.374452 | 0.986494 | -1.12115 | 0.87457 | 0.960733 | -1.01867 | 0.418274 | 0.998377 |
| 1.243005 | 0.678661 | 0.986494 | 1.071509 | 0.001139 | 0.063787 | -1.76341 | 0.784576 | 0.998377 |
| 1.062252 | 0.434455 | 0.986494 | 1.106446 | 0.003925 | 0.076269 | -1.46292 | 0.187195 | 0.998377 |
| 1.114419 | 0.562334 | 0.986494 | 1.125259 | 0.000942 | 0.063787 | -2.02548 | 0.965854 | 0.998377 |

| JAK2A - GH | JAK2A - GH | JAK2A - GH | JAK2A - GH | JAK2A - GH | JAK2A - GH | JAK2A - GH | JAK2A - GH | JAK2A - GH |
|---|---|---|---|---|---|---|---|---|
| -1.17251 | 0.431915 | 0.707182 | -1.19064 | 0.930565 | 0.947484 | -1.02062 | 0.944582 | 0.986244 |
| -1.24965 | 0.04519 | 0.407806 | -1.4003 | 0.912494 | 0.943983 | -1.01907 | 0.484334 | 0.785199 |
| -1.02783 | 0.684808 | 0.82303 | -1.02118 | 0.045643 | 0.112765 | 1.125439 | 0.898455 | 0.984719 |
| -1.06817 | 0.249945 | 0.574244 | -1.05933 | 0.282365 | 0.4198 | 1.058166 | 0.86234 | 0.965821 |
| -1.10139 | 0.185458 | 0.540821 | -1.22147 | 0.031337 | 0.084766 | 1.425158 | 0.488348 | 0.785199 |
| 1.010079 | 0.349807 | 0.657092 | -1.19557 | 0.367766 | 0.506432 | -1.2001 | 0.323857 | 0.647714 |
| -1.05326 | 0.320434 | 0.63795 | -1.0561 | 0.112064 | 0.213941 | 1.100878 | 0.959534 | 0.986244 |
| 1.037557 | 0.067095 | 0.41024 | -1.39036 | 0.010112 | 0.039508 | -1.65894 | 0.043171 | 0.576343 |
| 1.242106 | 0.456547 | 0.718501 | -1.2128 | 0.296444 | 0.429333 | -1.33424 | 0.120052 | 0.580313 |
| -1.07407 | 0.062665 | 0.41024 | -1.53094 | 0.005636 | 0.029006 | -2.01104 | 0.117853 | 0.580313 |
| 1.041252 | 0.328863 | 0.63795 | -1.24519 | 0.004435 | 0.029006 | -2.06346 | 0.249135 | 0.621396 |
| 1.037811 | 0.319424 | 0.63795 | -1.20941 | 0.008024 | 0.034563 | -1.76455 | 0.235818 | 0.609498 |
| -1.01118 | 0.019802 | 0.407806 | -1.55024 | 0.434648 | 0.566053 | -1.16083 | 0.022831 | 0.530639 |
| 1.05552 | 0.028935 | 0.407806 | 1.190637 | 0.429369 | 0.563547 | 1.06657 | 0.123137 | 0.580313 |
| 1.008461 | 0.872829 | 0.945616 | -1.0138 | 0.235431 | 0.375135 | 1.124654 | 0.796981 | 0.917846 |
| 1.047367 | 0.333335 | 0.63795 | -1.17713 | 0.013474 | 0.048163 | -1.58954 | 0.216722 | 0.596874 |
| -1.06828 | 0.408327 | 0.692918 | -1.346 | 0.058939 | 0.132023 | -2.09422 | 0.51917 | 0.785199 |
| -1.01055 | 0.185405 | 0.540821 | -1.31399 | 0.006043 | 0.029006 | -1.88452 | 0.202351 | 0.596874 |
| 1.004442 | 0.279349 | 0.594058 | -1.2462 | 0.003382 | 0.02711 | -1.96943 | 0.269984 | 0.621396 |
| -1.01095 | 0.077672 | 0.434965 | -1.43717 | 0.000456 | 0.017216 | -2.30187 | 0.086501 | 0.580313 |
| -1.01918 | 0.021681 | 0.407806 | -1.27683 | 0.000683 | 0.017216 | -1.45232 | 0.031566 | 0.530639 |
| -1.03306 | 0.156827 | 0.497602 | -1.53527 | 0.072657 | 0.157453 | -1.79136 | 0.189587 | 0.596874 |
| 1.062685 | 0.355925 | 0.657092 | -1.47 | 0.178296 | 0.302563 | -1.82404 | 0.286388 | 0.621396 |
| -1.13458 | 0.252941 | 0.574244 | -1.32682 | 0.054315 | 0.124999 | -1.67551 | 0.523466 | 0.785199 |
| 1.021585 | 0.331872 | 0.63795 | -1.45058 | 0.032787 | 0.086026 | -2.45299 | 0.305344 | 0.633306 |
| -1.0038 | 0.217349 | 0.540821 | -1.54782 | 0.021089 | 0.062186 | -2.45282 | 0.221256 | 0.599533 |
| -1.09268 | 0.220873 | 0.540821 | -1.25885 | 0.002089 | 0.023833 | -1.93723 | 0.447726 | 0.783298 |
| -1.05165 | 0.427414 | 0.707182 | -1.12978 | 0.049211 | 0.118107 | -1.39157 | 0.639905 | 0.826954 |
| -1.12812 | 0.353195 | 0.657092 | -1.29155 | 0.010669 | 0.040737 | -2.19563 | 0.621498 | 0.826954 |
| -1.0354 | 0.446283 | 0.718501 | -1.10991 | 0.006702 | 0.031278 | -1.42679 | 0.604598 | 0.819133 |
| 1.030255 | 0.206426 | 0.540821 | -1.32242 | 0.00197 | 0.023833 | -1.92996 | 0.169185 | 0.596874 |
| -1.09418 | 0.12306 | 0.480794 | -1.27955 | 0.004921 | 0.029006 | -1.65216 | 0.321392 | 0.647714 |
| 1.003898 | 0.113547 | 0.480794 | -1.61776 | 0.001088 | 0.0203 | -2.46654 | 0.111395 | 0.580313 |
| -1.19177 | 0.140225 | 0.497602 | -1.20514 | 0.222457 | 0.359354 | -1.15007 | 0.921881 | 0.984719 |
| -1.05009 | 0.115942 | 0.480794 | -1.69944 | 0.004235 | 0.029006 | -2.943 | 0.151961 | 0.596874 |
| -1.11888 | 0.044114 | 0.407806 | -1.47932 | 0.007754 | 0.034281 | -1.75981 | 0.143972 | 0.596874 |
| -1.06974 | 0.146498 | 0.497602 | -1.57162 | 0.011643 | 0.043467 | -2.37918 | 0.21427 | 0.596874 |
| -1.11215 | 0.10952 | 0.480794 | -1.59701 | 0.086556 | 0.173113 | -1.70504 | 0.211883 | 0.596874 |
| 1.354853 | 0.068373 | 0.41024 | -1.68023 | 0.000514 | 0.017216 | -3.12541 | 0.005548 | 0.433774 |
| -1.06818 | 0.104566 | 0.480794 | -1.58029 | 0.031787 | 0.084766 | -1.92303 | 0.162151 | 0.596874 |
| -1.30254 | 0.035369 | 0.407806 | -1.60697 | 0.662109 | 0.777862 | 1.105731 | 0.33595 | 0.661686 |
| -1.06514 | 0.272234 | 0.587858 | -1.52569 | 0.0403 | 0.102583 | -2.36392 | 0.348903 | 0.666087 |
| -1.0709 | 0.152003 | 0.497602 | -1.67507 | 0.022766 | 0.065943 | -2.44842 | 0.212062 | 0.596874 |
| -1.04717 | 0.15965 | 0.497602 | -1.73574 | 0.065954 | 0.145792 | -2.16996 | 0.196237 | 0.596874 |
| -1.00065 | 0.110252 | 0.480794 | -1.6872 | 0.015341 | 0.052598 | -2.38351 | 0.110676 | 0.580313 |
| -1.18242 | 0.151629 | 0.497602 | -1.58286 | 0.080902 | 0.163754 | -1.81921 | 0.357175 | 0.674218 |
| -1.08331 | 0.220317 | 0.540821 | -1.26923 | 0.003174 | 0.02711 | -1.91754 | 0.411958 | 0.760537 |

| | | | | | | | | |
|---|---|---|---|---|---|---|---|---|
| -1.02389 | 0.222835 | 0.540821 | -1.36673 | 0.001526 | 0.023607 | -2.07039 | 0.253656 | 0.621396 |
| -1.17234 | 0.053232 | 0.407806 | -1.35631 | 0.623925 | 0.738165 | -1.08264 | 0.342659 | 0.661686 |
| -1.11786 | 0.031812 | 0.407806 | -1.52325 | 0.018645 | 0.059102 | -1.63789 | 0.107778 | 0.580313 |
| -1.01589 | 0.034235 | 0.407806 | -1.28694 | 0.00239 | 0.025098 | -1.44935 | 0.044598 | 0.576343 |
| 1.045811 | 0.047536 | 0.407806 | -1.32752 | 0.252538 | 0.38222 | -1.20479 | 0.026069 | 0.530639 |
| 1.008747 | 0.193671 | 0.540821 | -1.22609 | 0.001551 | 0.023607 | -1.76681 | 0.175876 | 0.596874 |
| -1.08853 | 0.032263 | 0.407806 | -1.69809 | 0.053681 | 0.124999 | -1.6521 | 0.068836 | 0.580313 |
| -1.15354 | 0.049212 | 0.407806 | -1.88943 | 0.010079 | 0.039508 | -2.47428 | 0.122059 | 0.580313 |
| -1.12594 | 0.121611 | 0.480794 | -1.64302 | 0.024144 | 0.06875 | -1.90454 | 0.209738 | 0.596874 |
| -1.03801 | 0.149511 | 0.497602 | -1.33261 | 0.003988 | 0.029006 | -1.9055 | 0.207796 | 0.596874 |
| -1.18323 | 0.205732 | 0.540821 | -1.45012 | 0.252406 | 0.38222 | -1.42702 | 0.484188 | 0.785199 |
| -1.01613 | 0.630589 | 0.804095 | 1.090669 | 0.691244 | 0.806452 | -1.07894 | 0.569308 | 0.787241 |
| -1.0915 | 0.327965 | 0.63795 | -1.15314 | 0.15399 | 0.266705 | -1.2491 | 0.703998 | 0.838806 |
| -1.14502 | 0.027806 | 0.407806 | -1.45558 | 0.525792 | 0.649507 | -1.11654 | 0.149105 | 0.596874 |
| -1.06957 | 0.008399 | 0.407806 | -1.31989 | 0.009618 | 0.03941 | -1.33449 | 0.040469 | 0.576343 |
| -1.00815 | 0.147034 | 0.497602 | -1.4323 | 0.006019 | 0.029006 | -2.13696 | 0.156039 | 0.596874 |
| 1.023271 | 0.071446 | 0.413896 | -1.79501 | 0.001686 | 0.023607 | -3.15976 | 0.061584 | 0.580313 |
| -1.07786 | 0.115357 | 0.480794 | -1.66227 | 0.016122 | 0.054169 | -2.33823 | 0.176903 | 0.596874 |
| -1.11892 | 0.260954 | 0.576845 | -1.26495 | 0.220336 | 0.359354 | -1.31309 | 0.554102 | 0.787241 |
| -1.01839 | 0.091663 | 0.480794 | -1.75206 | 0.003445 | 0.02711 | -2.96884 | 0.102159 | 0.580313 |
| 1.125494 | 0.041046 | 0.407806 | -1.79951 | 0.00093 | 0.019521 | -2.93571 | 0.015677 | 0.530639 |
| -1.10132 | 0.035173 | 0.407806 | -1.96327 | 0.003305 | 0.02711 | -2.8241 | 0.068195 | 0.580313 |
| 1.037435 | 0.197573 | 0.540821 | 1.096773 | 0.488345 | 0.614759 | 1.053509 | 0.433449 | 0.774675 |
| 1.05953 | 0.530039 | 0.785307 | -1.08256 | 0.40948 | 0.545973 | 1.139191 | 0.294391 | 0.621396 |
| 1.074867 | 0.599708 | 0.804095 | -1.04654 | 0.762241 | 0.858731 | 1.028156 | 0.180531 | 0.596874 |
| 1.020942 | 0.43357 | 0.707182 | 1.043739 | 0.25027 | 0.38222 | 1.069396 | 0.685026 | 0.832444 |
| -1.11711 | 0.558552 | 0.785307 | -1.13059 | 0.584069 | 0.705925 | -1.12951 | 0.954273 | 0.986244 |
| -1.012 | 0.911844 | 0.945616 | -1.01947 | 0.131136 | 0.23689 | 1.32938 | 0.966323 | 0.986244 |
| -1.29112 | 0.053066 | 0.407806 | -1.39979 | 0.073103 | 0.157453 | 1.389417 | 0.63122 | 0.826954 |
| 1.006503 | 0.035444 | 0.407806 | -1.31116 | 0.553235 | 0.673504 | 1.081154 | 0.031586 | 0.530639 |
| 1.057543 | 0.194031 | 0.540821 | -1.22425 | 0.980549 | 0.986421 | -1.00398 | 0.100474 | 0.580313 |
| -1.08084 | 0.067538 | 0.41024 | -1.23923 | 0.936404 | 0.947685 | -1.00968 | 0.235459 | 0.609498 |
| -1.09272 | 0.26009 | 0.576845 | -1.20754 | 0.141739 | 0.250655 | 1.300975 | 0.547429 | 0.787241 |
| 1.034997 | 0.042469 | 0.407806 | 1.13832 | 0.30032 | 0.431228 | 1.070591 | 0.12966 | 0.583378 |
| 1.033076 | 0.461097 | 0.718501 | 1.051505 | 0.355083 | 0.497116 | 1.072425 | 0.797652 | 0.917846 |
| -1.06308 | 0.566524 | 0.785307 | -1.07695 | 0.074604 | 0.158653 | 1.28537 | 0.919962 | 0.984719 |
| 1.03466 | 0.827064 | 0.926947 | 1.017291 | 0.221883 | 0.359354 | 1.10851 | 0.829194 | 0.947651 |
| 1.119702 | 0.953209 | 0.958332 | 1.010762 | 0.766724 | 0.858731 | 1.059129 | 0.575815 | 0.787241 |
| -1.03235 | 0.725986 | 0.846984 | -1.0336 | 0.620681 | 0.738165 | 1.051311 | 0.989524 | 0.989524 |
| 1.049625 | 0.639794 | 0.804095 | -1.01675 | 0.756579 | 0.858731 | 1.011727 | 0.074533 | 0.580313 |
| 1.026778 | 0.869624 | 0.945616 | 1.007415 | 0.117263 | 0.221351 | 1.084841 | 0.676745 | 0.829878 |
| 1.040955 | 0.548071 | 0.785307 | -1.02105 | 0.840361 | 0.916758 | 1.007417 | 0.086031 | 0.580313 |
| 1.030563 | 0.641362 | 0.804095 | 1.015465 | 0.480338 | 0.61134 | 1.025035 | 0.654158 | 0.829878 |
| 1.050556 | 0.272934 | 0.587858 | 1.042847 | 0.910294 | 0.943983 | -1.00453 | 0.845772 | 0.960066 |
| 1.033877 | 0.518785 | 0.785307 | 1.019678 | 0.836402 | 0.916758 | 1.006614 | 0.64641 | 0.828984 |
| 1.01249 | 0.621548 | 0.804095 | 1.015944 | 0.383277 | 0.5235 | 1.03022 | 0.915177 | 0.984719 |
| 1.03613 | 0.407423 | 0.692918 | 1.025599 | 0.54124 | 0.66371 | 1.01991 | 0.736419 | 0.863636 |
| 1.053458 | 0.535985 | 0.785307 | 1.027312 | 0.77274 | 0.859737 | -1.01339 | 0.563589 | 0.787241 |
| 1.047849 | 0.745445 | 0.863688 | 1.010891 | 0.908237 | 0.943983 | 1.004084 | 0.286366 | 0.621396 |

| | | | | | | | | |
|---|---|---|---|---|---|---|---|---|
| 1.075596 | 0.461893 | 0.718501 | 1.038672 | 0.757257 | 0.858731 | 1.016981 | 0.497862 | 0.785199 |
| 1.045578 | 0.619671 | 0.804095 | 1.018632 | 0.476341 | 0.610881 | 1.028563 | 0.483653 | 0.785199 |
| 1.031584 | 0.116456 | 0.480794 | -1.06592 | 0.078534 | 0.162885 | 1.079315 | 0.022822 | 0.530639 |
| 1.042319 | 0.896725 | 0.945616 | -1.00383 | 0.490344 | 0.614759 | -1.02186 | 0.131955 | 0.583378 |
| 1.038731 | 0.596517 | 0.804095 | -1.01793 | 0.893426 | 0.943983 | -1.00477 | 0.103998 | 0.580313 |
| 1.000721 | 0.557794 | 0.785307 | 1.028676 | 0.991866 | 0.991866 | 1.00052 | 0.56782 | 0.787241 |
| 1.031211 | 0.246033 | 0.574077 | -1.05588 | 0.005391 | 0.029006 | 1.184275 | 0.078432 | 0.580313 |
| 1.051772 | 0.188237 | 0.540821 | -1.04529 | 0.219416 | 0.359354 | 1.04473 | 0.007438 | 0.433774 |
| 1.057751 | 0.603658 | 0.804095 | -1.03638 | 0.00355 | 0.02711 | 1.258366 | 0.187824 | 0.596874 |
| 1.009391 | 0.009692 | 0.407806 | -1.27849 | 0.919226 | 0.943983 | 1.011111 | 0.007746 | 0.433774 |
| -1.00334 | 0.018046 | 0.407806 | -1.22415 | 0.352805 | 0.497116 | -1.08913 | 0.019665 | 0.530639 |
| 1.042723 | 0.659817 | 0.812936 | 1.015337 | 0.899217 | 0.943983 | 1.004648 | 0.443121 | 0.783298 |
| 1.041653 | 0.824871 | 0.926947 | 1.006874 | 0.78405 | 0.866581 | 1.009042 | 0.27764 | 0.621396 |
| 1.026115 | 0.910866 | 0.945616 | -1.00514 | 0.46672 | 0.603145 | -1.03617 | 0.501579 | 0.785199 |
| 1.084581 | 0.658854 | 0.812936 | -1.03212 | 0.921507 | 0.943983 | 1.0075 | 0.122558 | 0.580313 |
| -1.00357 | 0.053403 | 0.407806 | -1.10738 | 0.602917 | 0.723501 | -1.02863 | 0.061727 | 0.580313 |
| 1.044701 | 0.212231 | 0.540821 | 1.043325 | 0.880565 | 0.943983 | -1.00536 | 0.968632 | 0.986244 |
| 1.040213 | 0.954488 | 0.958332 | -1.00173 | 0.701502 | 0.807208 | 1.012388 | 0.18071 | 0.596874 |
| 1.008661 | 0.398173 | 0.692918 | 1.032884 | 0.700197 | 0.807208 | 1.015682 | 0.534245 | 0.787241 |
| 1.161123 | 0.902795 | 0.945616 | -1.00958 | 0.238925 | 0.375135 | -1.10372 | 0.049294 | 0.580313 |
| -1.01086 | 0.476712 | 0.734749 | 1.09913 | 0.000239 | 0.017216 | 1.795206 | 0.428335 | 0.773767 |
| 1.018673 | 0.565205 | 0.785307 | -1.02471 | 0.339056 | 0.482723 | -1.04422 | 0.315016 | 0.645399 |
| 1.029568 | 0.868924 | 0.945616 | -1.00529 | 0.910656 | 0.943983 | -1.00393 | 0.293619 | 0.621396 |
| 1.04008 | 0.836312 | 0.930467 | -1.01151 | 0.238683 | 0.375135 | 1.079117 | 0.37233 | 0.695016 |
| 1.236371 | 0.827631 | 0.926947 | 1.087723 | 0.358775 | 0.498133 | -1.46026 | 0.740259 | 0.863636 |
| -1.08632 | 0.909418 | 0.945616 | -1.04667 | 0.510752 | 0.635602 | -1.32408 | 0.926105 | 0.984719 |
| -1.04471 | 0.895556 | 0.945616 | -1.03691 | 0.121245 | 0.223837 | -1.58996 | 0.97834 | 0.988341 |
| -1.04166 | 0.816855 | 0.926947 | 1.049829 | 0.406218 | 0.545956 | -1.17433 | 0.664048 | 0.829878 |
| 1.076255 | 0.634936 | 0.804095 | -1.18189 | 0.294164 | 0.429333 | -1.46242 | 0.510219 | 0.785199 |
| -1.07657 | 0.934544 | 0.958332 | 1.024551 | 0.423036 | 0.559607 | -1.28712 | 0.740155 | 0.863636 |
| -1.0066 | 0.863293 | 0.945616 | -1.05261 | 0.145242 | 0.254173 | -1.59826 | 0.880679 | 0.979828 |
| 1.025501 | 0.68716 | 0.82303 | -1.11122 | 0.057611 | 0.130792 | -1.72332 | 0.618143 | 0.826954 |
| -1.07009 | 0.766945 | 0.882512 | -1.08851 | 0.20847 | 0.350229 | -1.47256 | 0.952407 | 0.986244 |
| 1.04148 | 0.958332 | 0.958332 | -1.01695 | 0.110447 | 0.213276 | -1.74607 | 0.858286 | 0.965821 |
| 1.066317 | 0.70979 | 0.83388 | -1.09288 | 0.048964 | 0.118107 | -1.67496 | 0.522427 | 0.785199 |
| -1.1198 | 0.943202 | 0.958332 | -1.0099 | 0.250069 | 0.38222 | -1.18606 | 0.45701 | 0.783447 |
| -1.12122 | 0.334164 | 0.63795 | -1.1429 | 0.005435 | 0.029006 | -1.54408 | 0.888965 | 0.98254 |
| -1.19307 | 0.570283 | 0.785307 | -1.06164 | 0.285678 | 0.421 | 1.115878 | 0.230896 | 0.609498 |
| -1.00892 | 0.66293 | 0.812936 | -1.09919 | 0.014217 | 0.049761 | -1.81373 | 0.692799 | 0.832444 |
| -1.01908 | 0.886847 | 0.945616 | -1.02261 | 0.005715 | 0.029006 | -1.63936 | 0.982458 | 0.988341 |
| 1.021173 | 0.382968 | 0.68817 | -1.25584 | 0.002128 | 0.023833 | -2.51555 | 0.341384 | 0.661686 |
| -1.02645 | 0.955231 | 0.958332 | -1.00931 | 0.025759 | 0.07119 | -1.50455 | 0.918684 | 0.984719 |
| 1.026374 | 0.62967 | 0.804095 | -1.13356 | 0.005033 | 0.029006 | -2.2939 | 0.560726 | 0.787241 |
| 1.041611 | 0.689537 | 0.82303 | -1.10716 | 0.005252 | 0.029006 | -2.24736 | 0.576373 | 0.787241 |
| 1.040213 | 0.626031 | 0.804095 | -1.12798 | 0.007232 | 0.032836 | -2.11916 | 0.518404 | 0.785199 |
| -1.02429 | 0.596441 | 0.804095 | -1.14261 | 0.018067 | 0.058369 | -1.94063 | 0.663872 | 0.829878 |
| -1.03879 | 0.41468 | 0.696663 | -1.1459 | 0.000644 | 0.017216 | -1.95409 | 0.555577 | 0.787241 |
| 1.046911 | 0.368004 | 0.672008 | -1.30221 | 0.00492 | 0.029006 | -2.54632 | 0.292055 | 0.621396 |
| -1.00062 | 0.691281 | 0.82303 | -1.07728 | 0.020098 | 0.062186 | -1.62421 | 0.693703 | 0.832444 |

| | | | | | | | | |
|---|---|---|---|---|---|---|---|---|
| 1.01989 | 0.527789 | 0.785307 | -1.14025 | 0.01645 | 0.05419 | -1.74312 | 0.468273 | 0.785199 |
| -1.20437 | 0.225342 | 0.540821 | -1.22114 | 0.887601 | 0.943983 | -1.02468 | 0.932218 | 0.984985 |
| -1.14225 | 0.061188 | 0.41024 | -1.55309 | 0.045025 | 0.112765 | -1.65257 | 0.184286 | 0.596874 |
| -1.03171 | 0.199955 | 0.540821 | -1.22347 | 0.049951 | 0.118193 | -1.39642 | 0.276645 | 0.621396 |
| -1.08651 | 0.080789 | 0.437825 | -2.18694 | 0.033284 | 0.086026 | -2.78928 | 0.116548 | 0.580313 |
| 1.041229 | 0.385048 | 0.68817 | -1.15783 | 0.003145 | 0.02711 | -1.76683 | 0.269837 | 0.621396 |
| -1.01605 | 0.399406 | 0.692918 | -1.15695 | 0.080569 | 0.163754 | -1.38764 | 0.452261 | 0.783298 |
| 1.00928 | 0.540969 | 0.785307 | -1.14882 | 0.009531 | 0.03941 | -1.93811 | 0.514562 | 0.785199 |
| 1.033813 | 0.153092 | 0.497602 | -1.52495 | 0.025849 | 0.07119 | -2.04879 | 0.124353 | 0.580313 |
| 1.034209 | 0.23001 | 0.544248 | -1.44543 | 0.012575 | 0.045928 | -2.33794 | 0.191303 | 0.596874 |
| -1.02006 | 0.120342 | 0.480794 | -1.38416 | 0.077308 | 0.162347 | -1.48394 | 0.143709 | 0.596874 |
| 1.012765 | 0.322556 | 0.63795 | -1.24562 | 0.021099 | 0.062186 | -1.75997 | 0.295903 | 0.621396 |
| 1.052348 | 0.457907 | 0.718501 | -1.12444 | 0.000717 | 0.017216 | -1.87378 | 0.289287 | 0.621396 |
| -1.10327 | 0.159944 | 0.497602 | -1.25213 | 0.124221 | 0.226838 | -1.2993 | 0.423282 | 0.772949 |
| -1.14208 | 0.129673 | 0.495114 | -1.53661 | 0.091711 | 0.181264 | -1.66468 | 0.290012 | 0.621396 |
| -1.03544 | 0.528787 | 0.785307 | -1.10869 | 0.1199 | 0.223813 | -1.31686 | 0.675892 | 0.829878 |
| -1.03012 | 0.407224 | 0.692918 | -1.16408 | 0.176098 | 0.301883 | -1.30448 | 0.503891 | 0.785199 |
| -1.09181 | 0.287746 | 0.604266 | -1.17004 | 0.277428 | 0.416142 | -1.1856 | 0.636665 | 0.826954 |
| -1.03912 | 0.555339 | 0.785307 | -1.20657 | 0.401945 | 0.544571 | -1.32838 | 0.638502 | 0.826954 |
| -1.09077 | 0.695657 | 0.82303 | -1.04367 | 0.093132 | 0.181933 | 1.229663 | 0.673146 | 0.829878 |
| -1.04157 | 0.183879 | 0.540821 | -1.22269 | 0.020946 | 0.062186 | -1.46661 | 0.286494 | 0.621396 |
| -1.16679 | 0.058598 | 0.41024 | -1.25227 | 0.139687 | 0.249653 | -1.20197 | 0.54047 | 0.787241 |
| 1.007806 | 0.094753 | 0.480794 | -1.36522 | 0.000686 | 0.017216 | -2.07236 | 0.087181 | 0.580313 |

| JAK2A - Gl | JAK2A - Ve | JAK2A - Ve | JAK2A - Ve | JAK2A - Ve | JAK2A - Ve | JAK2A - Ve | JAK2A - Ve | JAK2A - Ve |
|---|---|---|---|---|---|---|---|---|
| -1.01547 | 0.145367 | 0.34612 | -1.41571 | 0.068655 | 0.851606 | -1.51339 | 0.659828 | 0.842885 |
| -1.12055 | 0.266665 | 0.476593 | -1.21294 | 0.01456 | 0.64007 | -1.51932 | 0.995777 | 0.995777 |
| 1.006511 | 0.365528 | 0.54854 | -1.05566 | 0.032478 | 0.779464 | -1.12838 | 0.13751 | 0.618466 |
| 1.008343 | 0.157793 | 0.34612 | -1.0836 | 0.016127 | 0.64007 | -1.13905 | 0.211982 | 0.618466 |
| -1.10903 | 0.431723 | 0.594504 | 1.132806 | 0.086338 | 0.851606 | -1.29948 | 0.241343 | 0.618466 |
| -1.20762 | 0.188388 | 0.390731 | -1.30818 | 0.106451 | 0.851606 | -1.36801 | 0.262241 | 0.618466 |
| -1.0027 | 0.236695 | 0.449862 | -1.07766 | 0.017452 | 0.64007 | -1.15299 | 0.157617 | 0.618466 |
| -1.44258 | 0.438236 | 0.598566 | -1.15523 | 0.453064 | 0.983907 | -1.14069 | 0.467165 | 0.714789 |
| -1.50642 | 0.366957 | 0.54854 | 1.282157 | 0.935206 | 0.983907 | -1.02118 | 0.856232 | 0.924011 |
| -1.42536 | 0.290965 | 0.514549 | -1.28502 | 0.762601 | 0.983907 | -1.0693 | 0.193721 | 0.618466 |
| -1.29655 | 0.110575 | 0.300122 | -1.4706 | 0.800685 | 0.983907 | -1.05785 | 0.145053 | 0.618466 |
| -1.25513 | 0.08234 | 0.273903 | -1.43115 | 0.788982 | 0.983907 | -1.05199 | 0.147412 | 0.618466 |
| -1.5331 | 0.082581 | 0.273903 | 1.403177 | 0.522213 | 0.983907 | -1.12183 | 0.351486 | 0.653688 |
| 1.128011 | 0.976275 | 0.984707 | -1.00241 | 0.233497 | 0.939037 | -1.09666 | 0.015682 | 0.439091 |
| -1.02238 | 0.456843 | 0.609124 | 1.075087 | 0.68858 | 0.983907 | -1.03534 | 0.302535 | 0.651614 |
| -1.23288 | 0.230516 | 0.445134 | -1.24031 | 0.702339 | 0.983907 | -1.06605 | 0.29231 | 0.646158 |
| -1.25997 | 0.381859 | 0.553037 | -1.39593 | 0.710953 | 0.983907 | -1.14169 | 0.354081 | 0.653688 |
| -1.30027 | 0.142424 | 0.34612 | -1.37985 | 0.489742 | 0.983907 | -1.15117 | 0.210208 | 0.618466 |
| -1.25173 | 0.070137 | 0.273455 | -1.48953 | 0.648742 | 0.983907 | -1.09622 | 0.110075 | 0.618466 |
| -1.42161 | 0.045117 | 0.236282 | -1.55346 | 0.522628 | 0.983907 | -1.13678 | 0.093504 | 0.618466 |
| -1.25281 | 0.048264 | 0.236282 | -1.23201 | 0.08694 | 0.851606 | -1.18789 | 0.989538 | 0.995463 |
| -1.48613 | 0.932964 | 0.949927 | -1.02688 | 0.457998 | 0.983907 | -1.24827 | 0.787243 | 0.907617 |
| -1.56215 | 0.90086 | 0.940028 | 1.05625 | 0.636022 | 0.983907 | -1.21699 | 0.927073 | 0.96738 |
| -1.16944 | 0.588346 | 0.721475 | -1.15107 | 0.234759 | 0.939037 | -1.34204 | 0.918983 | 0.966058 |
| -1.48189 | 0.452066 | 0.607576 | -1.35621 | 0.619065 | 0.983907 | -1.2085 | 0.461109 | 0.714789 |
| -1.54196 | 0.541526 | 0.684033 | -1.25473 | 0.76302 | 0.983907 | -1.11114 | 0.538154 | 0.786173 |
| -1.15207 | 0.048962 | 0.236282 | -1.49388 | 0.218869 | 0.939037 | -1.26014 | 0.214313 | 0.618466 |
| -1.0743 | 0.259648 | 0.473817 | -1.20311 | 0.326379 | 0.983907 | -1.16342 | 0.783099 | 0.907617 |
| -1.14488 | 0.143508 | 0.34612 | -1.54127 | 0.350597 | 0.983907 | -1.29337 | 0.193806 | 0.618466 |
| -1.07196 | 0.102099 | 0.300122 | -1.24283 | 0.678669 | 0.983907 | -1.05671 | 0.145599 | 0.618466 |
| -1.36243 | 0.106153 | 0.300122 | -1.38422 | 0.492046 | 0.983907 | -1.15105 | 0.168443 | 0.618466 |
| -1.16942 | 0.373201 | 0.54854 | -1.16046 | 0.196649 | 0.939037 | -1.2276 | 0.409236 | 0.695906 |
| -1.62407 | 0.262292 | 0.473817 | -1.32055 | 0.626296 | 0.983907 | -1.13007 | 0.115944 | 0.618466 |
| -1.01122 | 0.930692 | 0.949927 | 1.012651 | 0.153591 | 0.922415 | -1.19685 | 0.721215 | 0.871684 |
| -1.61837 | 0.313515 | 0.517358 | -1.42737 | 0.80869 | 0.983907 | -1.08317 | 0.193301 | 0.618466 |
| -1.32213 | 0.147126 | 0.34612 | -1.3415 | 0.101608 | 0.851606 | -1.3695 | 0.579675 | 0.818365 |
| -1.46915 | 0.354353 | 0.54854 | -1.3531 | 0.502005 | 0.983907 | -1.22862 | 0.464468 | 0.714789 |
| -1.43597 | 0.448458 | 0.607576 | -1.25979 | 0.292038 | 0.983907 | -1.35547 | 0.936074 | 0.970743 |
| -2.27647 | 0.307108 | 0.517358 | -1.35263 | 0.517564 | 0.983907 | -1.19652 | 0.610097 | 0.841398 |
| -1.47942 | 0.565991 | 0.704345 | -1.183 | 0.378284 | 0.983907 | -1.27661 | 0.85801 | 0.924011 |
| -1.23373 | 0.741428 | 0.847346 | 1.078781 | 0.01905 | 0.64007 | -1.70551 | 0.069734 | 0.618466 |
| -1.43238 | 0.404251 | 0.575544 | -1.40327 | 0.613715 | 0.983907 | -1.21233 | 0.432502 | 0.712356 |
| -1.56417 | 0.661279 | 0.771492 | -1.17885 | 0.679684 | 0.983907 | -1.15738 | 0.669227 | 0.842885 |
| -1.65755 | 0.60015 | 0.729942 | -1.23944 | 0.438899 | 0.983907 | -1.3495 | 0.616294 | 0.841767 |
| -1.6861 | 0.247214 | 0.456395 | -1.48816 | 0.157134 | 0.922415 | -1.58572 | 0.79884 | 0.91296 |
| -1.33867 | 0.890015 | 0.934516 | 1.047194 | 0.639551 | 0.983907 | -1.15894 | 0.952217 | 0.981426 |
| -1.17162 | 0.044378 | 0.236282 | -1.529 | 0.195749 | 0.939037 | -1.28666 | 0.192102 | 0.618466 |

| | | | | | | | | |
|---|---|---|---|---|---|---|---|---|
| -1.33484 | 0.152349 | 0.34612 | -1.37716 | 0.615172 | 0.983907 | -1.12286 | 0.133063 | 0.618466 |
| -1.15692 | 0.820439 | 0.88355 | 1.037378 | 0.27981 | 0.983907 | -1.18106 | 0.751914 | 0.893617 |
| -1.36265 | 0.134886 | 0.339557 | -1.35545 | 0.08127 | 0.851606 | -1.4005 | 0.788762 | 0.907617 |
| -1.26681 | 0.090203 | 0.291425 | -1.21243 | 0.232083 | 0.939037 | -1.1394 | 0.987028 | 0.995463 |
| -1.38833 | 0.765621 | 0.848093 | -1.05513 | 0.014398 | 0.64007 | -1.45057 | 0.474822 | 0.714789 |
| -1.23681 | 0.036085 | 0.216508 | -1.42964 | 0.373358 | 0.983907 | -1.1485 | 0.488191 | 0.725807 |
| -1.55998 | 0.42249 | 0.591485 | -1.22495 | 0.105017 | 0.851606 | -1.48242 | 0.700353 | 0.869766 |
| -1.63794 | 0.21676 | 0.428419 | -1.51407 | 0.760925 | 0.983907 | -1.09975 | 0.387551 | 0.685353 |
| -1.45925 | 0.60394 | 0.729942 | -1.17565 | 0.370061 | 0.983907 | -1.28755 | 0.816884 | 0.92105 |
| -1.28381 | 0.156105 | 0.34612 | -1.34921 | 0.507052 | 0.983907 | -1.13907 | 0.214926 | 0.618466 |
| -1.22556 | 0.864611 | 0.919333 | 1.053756 | 0.434463 | 0.983907 | -1.25544 | 0.475814 | 0.714789 |
| 1.108259 | 0.809027 | 0.87688 | -1.04727 | 0.80433 | 0.983907 | -1.04566 | 0.71795 | 0.871684 |
| -1.05647 | 0.984707 | 0.984707 | -1.00294 | 0.221902 | 0.939037 | -1.19577 | 0.571625 | 0.813839 |
| -1.27122 | 0.517315 | 0.66853 | -1.11909 | 0.135306 | 0.922415 | -1.28244 | 0.425904 | 0.708435 |
| -1.23403 | 0.098951 | 0.300122 | -1.19384 | 0.023128 | 0.647592 | -1.26517 | 0.138324 | 0.618466 |
| -1.42072 | 0.20155 | 0.4031 | -1.39679 | 0.491438 | 0.983907 | -1.18288 | 0.131249 | 0.618466 |
| -1.83678 | 0.371366 | 0.54854 | -1.35102 | 0.711328 | 0.983907 | -1.12381 | 0.222382 | 0.618466 |
| -1.54219 | 0.363728 | 0.54854 | -1.35833 | 0.463877 | 0.983907 | -1.26142 | 0.567338 | 0.813839 |
| -1.13051 | 0.367745 | 0.54854 | 1.220079 | 0.93039 | 0.983907 | 1.01822 | 0.530152 | 0.781277 |
| -1.72041 | 0.315177 | 0.517358 | -1.41665 | 0.773415 | 0.983907 | -1.09783 | 0.154536 | 0.618466 |
| -2.02534 | 0.113232 | 0.300122 | -1.60914 | 0.460224 | 0.983907 | -1.22801 | 0.238751 | 0.618466 |
| -1.78265 | 0.491947 | 0.645681 | -1.25244 | 0.790956 | 0.983907 | -1.08498 | 0.235476 | 0.618466 |
| 1.057197 | 0.644597 | 0.757289 | 1.035222 | 0.077554 | 0.851606 | 1.136826 | 0.835924 | 0.923917 |
| -1.147 | 0.05158 | 0.236282 | 1.377442 | 0.644554 | 0.983907 | -1.05925 | 0.047406 | 0.618466 |
| -1.12489 | 0.863796 | 0.919333 | 1.01586 | 0.95455 | 0.983907 | -1.00494 | 0.651698 | 0.842885 |
| 1.022329 | 0.375489 | 0.54854 | 1.052815 | 0.307429 | 0.983907 | 1.057611 | 0.444373 | 0.714789 |
| -1.01207 | 0.297829 | 0.517358 | 1.262715 | 0.335394 | 0.983907 | 1.225318 | 0.654966 | 0.842885 |
| -1.00738 | 0.365866 | 0.54854 | 1.18326 | 0.893356 | 0.983907 | -1.02363 | 0.828248 | 0.923917 |
| -1.08417 | 0.055812 | 0.240419 | 1.422497 | 0.211348 | 0.939037 | 1.237374 | 0.856731 | 0.924011 |
| -1.31968 | 0.313629 | 0.517358 | 1.142718 | 0.420242 | 0.983907 | -1.10548 | 0.268216 | 0.618466 |
| -1.29469 | 0.687324 | 0.790893 | 1.067787 | 0.532879 | 0.983907 | -1.10077 | 0.276101 | 0.618466 |
| -1.14654 | 0.980293 | 0.984707 | 1.002986 | 0.400608 | 0.983907 | -1.10105 | 0.461864 | 0.714789 |
| -1.10508 | 0.045215 | 0.236282 | 1.440215 | 0.972006 | 0.983907 | 1.005826 | 0.729764 | 0.875717 |
| 1.099829 | 0.609088 | 0.730905 | 1.033995 | 0.365853 | 0.983907 | 1.057615 | 0.410087 | 0.695906 |
| 1.017839 | 0.426442 | 0.592084 | 1.058352 | 0.318495 | 0.983907 | 1.069997 | 0.958302 | 0.981676 |
| -1.01304 | 0.507087 | 0.660392 | 1.095374 | 0.138986 | 0.922415 | 1.214875 | 0.405095 | 0.695906 |
| -1.01707 | 0.474154 | 0.627228 | 1.06165 | 0.731295 | 0.983907 | -1.02733 | 0.375914 | 0.679071 |
| -1.10778 | 0.31719 | 0.517358 | 1.215684 | 0.489364 | 0.983907 | 1.135033 | 0.476526 | 0.714789 |
| -1.00122 | 0.777418 | 0.848093 | 1.027654 | 0.720307 | 0.983907 | 1.033216 | 0.3431 | 0.653688 |
| -1.06721 | 0.004328 | 0.062906 | -1.12258 | 0.388344 | 0.983907 | -1.03125 | 0.081605 | 0.618466 |
| -1.01922 | 0.916968 | 0.945095 | 1.005147 | 0.587218 | 0.983907 | -1.0252 | 0.032873 | 0.613636 |
| -1.06287 | 0.004097 | 0.062906 | -1.12034 | 0.094823 | 0.851606 | -1.06104 | 0.321706 | 0.653688 |
| -1.01487 | 0.0082 | 0.091844 | -1.10337 | 0.297456 | 0.983907 | -1.03521 | 0.310202 | 0.653688 |
| -1.00739 | 0.01774 | 0.139006 | -1.10545 | 0.100827 | 0.851606 | -1.06573 | 0.348425 | 0.653688 |
| -1.01393 | 0.000789 | 0.026672 | -1.12641 | 0.078527 | 0.851606 | -1.05596 | 0.405197 | 0.695906 |
| 1.003412 | 0.001319 | 0.027698 | -1.12743 | 0.162351 | 0.922415 | -1.04652 | 0.332304 | 0.653688 |
| -1.01027 | 0.000949 | 0.026672 | -1.12494 | 0.063548 | 0.851606 | -1.05978 | 0.249702 | 0.618466 |
| -1.02545 | 0.025835 | 0.180842 | -1.11333 | 0.249299 | 0.951868 | -1.05189 | 0.352184 | 0.653688 |
| -1.03656 | 0.000953 | 0.026672 | -1.13813 | 0.127574 | 0.922415 | -1.05325 | 0.218169 | 0.618466 |

| | | | | | | | | |
|---|---|---|---|---|---|---|---|---|
| -1.03555 | 0.127365 | 0.329189 | -1.08846 | 0.570139 | 0.983907 | -1.02966 | 0.371932 | 0.679071 |
| -1.02645 | 0.113135 | 0.300122 | -1.06588 | 0.556857 | 0.983907 | -1.02211 | 0.346389 | 0.653688 |
| -1.09959 | 0.107222 | 0.300122 | -1.07204 | 0.930411 | 0.983907 | -1.00348 | 0.064857 | 0.618466 |
| -1.04631 | 0.001267 | 0.027698 | -1.11727 | 0.28805 | 0.983907 | -1.0321 | 0.718945 | 0.871684 |
| -1.05736 | 0.000666 | 0.026672 | -1.14424 | 0.164717 | 0.922415 | -1.04849 | 0.254071 | 0.618466 |
| 1.027935 | 0.024581 | 0.179547 | -1.12762 | 0.498164 | 0.983907 | -1.03325 | 0.804441 | 0.913149 |
| -1.08883 | 0.752552 | 0.848093 | -1.01614 | 0.152412 | 0.922415 | -1.07033 | 0.008543 | 0.427761 |
| -1.09941 | 0.160699 | 0.34612 | -1.05147 | 0.119423 | 0.911958 | -1.05417 | 0.074084 | 0.618466 |
| -1.09624 | 0.74934 | 0.848093 | 1.023567 | 0.708242 | 0.983907 | -1.02606 | 0.01001 | 0.427761 |
| -1.29049 | 0.537429 | 0.684001 | -1.05214 | 0.965707 | 0.983907 | -1.00342 | 0.760638 | 0.893617 |
| -1.22007 | 0.342571 | 0.545342 | -1.07589 | 0.823152 | 0.983907 | -1.01672 | 0.834953 | 0.923917 |
| -1.02697 | 0.009647 | 0.101297 | -1.10607 | 0.242691 | 0.948187 | -1.04168 | 0.326153 | 0.653688 |
| -1.03454 | 0.007022 | 0.090742 | -1.09927 | 0.294031 | 0.983907 | -1.03334 | 0.599946 | 0.839924 |
| -1.03139 | 0.028677 | 0.185299 | -1.11748 | 0.883234 | 0.983907 | 1.006752 | 0.626171 | 0.842885 |
| -1.11942 | 0.018203 | 0.139006 | -1.20742 | 0.314836 | 0.983907 | -1.07521 | 0.010594 | 0.427761 |
| -1.10343 | 0.104282 | 0.300122 | -1.09424 | 0.648822 | 0.983907 | -1.02355 | 0.902245 | 0.96546 |
| -1.00132 | 0.000391 | 0.026672 | -1.15245 | 0.05308 | 0.851606 | -1.0694 | 0.226249 | 0.618466 |
| -1.04201 | 0.004493 | 0.062906 | -1.10323 | 0.184003 | 0.936745 | -1.04168 | 0.217815 | 0.618466 |
| 1.024015 | 0.010591 | 0.10466 | -1.11579 | 0.174155 | 0.929685 | -1.05399 | 0.094502 | 0.618466 |
| -1.17224 | 0.000161 | 0.026672 | -1.42937 | 0.043028 | 0.851606 | -1.1781 | 0.242962 | 0.618466 |
| 1.111066 | 0.08109 | 0.273903 | 1.285995 | 0.978272 | 0.983907 | -1.00361 | 0.107461 | 0.618466 |
| -1.04385 | 0.011878 | 0.110864 | -1.12718 | 0.623945 | 0.983907 | -1.02101 | 0.756346 | 0.893617 |
| -1.03502 | 0.003009 | 0.056174 | -1.11391 | 0.177083 | 0.929685 | -1.04577 | 0.181148 | 0.618466 |
| -1.05205 | 0.878955 | 0.928707 | 1.009479 | 0.343102 | 0.983907 | -1.05555 | 0.02802 | 0.613636 |
| -1.13666 | 0.91254 | 0.945095 | 1.046001 | 0.216344 | 0.939037 | -1.62241 | 0.143363 | 0.618466 |
| 1.037881 | 0.679558 | 0.78735 | -1.19231 | 0.419979 | 0.983907 | -1.38429 | 0.54875 | 0.794742 |
| 1.007522 | 0.163429 | 0.347544 | -1.51482 | 0.781955 | 0.983907 | -1.07945 | 0.263804 | 0.618466 |
| 1.093565 | 0.576829 | 0.712554 | -1.1396 | 0.690871 | 0.983907 | -1.09367 | 0.704097 | 0.869766 |
| -1.27201 | 0.772395 | 0.848093 | 1.139164 | 0.564267 | 0.983907 | -1.23009 | 0.415108 | 0.697381 |
| 1.103002 | 0.546486 | 0.685147 | -1.20865 | 0.847323 | 0.983907 | -1.05852 | 0.672301 | 0.842885 |
| -1.04571 | 0.420256 | 0.591485 | -1.29176 | 0.983907 | 0.983907 | 1.006023 | 0.445071 | 0.714789 |
| -1.13956 | 0.246685 | 0.456395 | -1.38435 | 0.838311 | 0.983907 | 1.054827 | 0.27328 | 0.618466 |
| -1.01721 | 0.39156 | 0.56224 | -1.299 | 0.900054 | 0.983907 | -1.03657 | 0.350877 | 0.653688 |
| -1.05913 | 0.305917 | 0.517358 | -1.4228 | 0.927854 | 0.983907 | 1.029554 | 0.297632 | 0.64938 |
| -1.16535 | 0.23832 | 0.449862 | -1.35245 | 0.81981 | 0.983907 | 1.055824 | 0.256571 | 0.618466 |
| 1.10883 | 0.221185 | 0.432083 | -1.19935 | 0.800524 | 0.983907 | 1.035556 | 0.200555 | 0.618466 |
| -1.01933 | 0.017259 | 0.139006 | -1.44042 | 0.867537 | 0.983907 | 1.023143 | 0.03017 | 0.613636 |
| 1.123798 | 0.625263 | 0.744994 | 1.051994 | 0.486245 | 0.983907 | 1.071329 | 0.638948 | 0.842885 |
| -1.08947 | 0.077364 | 0.273903 | -1.51819 | 0.904853 | 0.983907 | -1.02623 | 0.15894 | 0.618466 |
| -1.00346 | 0.032345 | 0.201257 | -1.45054 | 0.742553 | 0.983907 | 1.053012 | 0.012731 | 0.427761 |
| -1.28243 | 0.028163 | 0.185299 | -1.87959 | 0.782891 | 0.983907 | -1.07412 | 0.087296 | 0.618466 |
| 1.016989 | 0.073247 | 0.273455 | -1.38103 | 0.977802 | 0.983907 | -1.0046 | 0.096237 | 0.618466 |
| -1.16345 | 0.049271 | 0.236282 | -1.75168 | 0.918087 | 0.983907 | -1.02704 | 0.100303 | 0.618466 |
| -1.15323 | 0.052038 | 0.236282 | -1.72085 | 0.96364 | 0.983907 | 1.011668 | 0.107312 | 0.618466 |
| -1.17334 | 0.072394 | 0.273455 | -1.62252 | 0.951554 | 0.983907 | 1.015095 | 0.136063 | 0.618466 |
| -1.11552 | 0.107747 | 0.300122 | -1.55061 | 0.962287 | 0.983907 | -1.01195 | 0.269763 | 0.618466 |
| -1.1031 | 0.007608 | 0.091292 | -1.65174 | 0.797886 | 0.983907 | 1.043442 | 0.008326 | 0.427761 |
| -1.3633 | 0.053677 | 0.237308 | -1.85257 | 0.933859 | 0.983907 | -1.02446 | 0.129137 | 0.618466 |
| -1.07661 | 0.156508 | 0.34612 | -1.33175 | 0.782181 | 0.983907 | 1.053149 | 0.173344 | 0.618466 |

| | | | | | | | | |
|---|---|---|---|---|---|---|---|---|
| -1.16293 | 0.114332 | 0.300122 | -1.426 | 0.939215 | 0.983907 | 1.015921 | 0.19699 | 0.618466 |
| -1.01393 | 0.775929 | 0.848093 | -1.05035 | 0.759902 | 0.983907 | -1.05099 | 0.642622 | 0.842885 |
| -1.35968 | 0.165727 | 0.348027 | -1.40595 | 0.796461 | 0.983907 | -1.06054 | 0.3824 | 0.683437 |
| -1.18587 | 0.33162 | 0.535695 | -1.1747 | 0.510971 | 0.983907 | 1.107721 | 0.118943 | 0.618466 |
| -2.01282 | 0.158956 | 0.34612 | -1.94062 | 0.796407 | 0.983907 | -1.11908 | 0.451833 | 0.714789 |
| -1.20557 | 0.058264 | 0.24471 | -1.41595 | 0.854525 | 0.983907 | 1.031208 | 0.189493 | 0.618466 |
| -1.13868 | 0.1923 | 0.390985 | -1.27305 | 0.656655 | 0.983907 | -1.07956 | 0.655547 | 0.842885 |
| -1.15948 | 0.073241 | 0.273455 | -1.55659 | 0.947655 | 0.983907 | -1.01496 | 0.150452 | 0.618466 |
| -1.57651 | 0.100946 | 0.300122 | -1.67685 | 0.922665 | 0.983907 | -1.02854 | 0.247259 | 0.618466 |
| -1.49488 | 0.082954 | 0.273903 | -1.77286 | 0.894969 | 0.983907 | -1.04081 | 0.315959 | 0.653688 |
| -1.35694 | 0.52662 | 0.67536 | -1.14797 | 0.623758 | 0.983907 | 1.105884 | 0.346168 | 0.653688 |
| -1.26152 | 0.083149 | 0.273903 | -1.5154 | 0.904068 | 0.983907 | -1.02688 | 0.2281 | 0.618466 |
| -1.1833 | 0.060871 | 0.249421 | -1.38097 | 0.945842 | 0.983907 | 1.010737 | 0.124353 | 0.618466 |
| -1.13493 | 0.315728 | 0.517358 | -1.18384 | 0.907007 | 0.983907 | 1.018529 | 0.237989 | 0.618466 |
| -1.34546 | 0.193165 | 0.390985 | -1.47565 | 0.378234 | 0.983907 | -1.27933 | 0.846575 | 0.924011 |
| -1.07074 | 0.632751 | 0.748607 | -1.08643 | 0.397121 | 0.983907 | 1.149256 | 0.099997 | 0.618466 |
| -1.13004 | 0.106533 | 0.300122 | -1.37625 | 0.461241 | 0.983907 | -1.14441 | 0.910084 | 0.966058 |
| -1.07166 | 0.135419 | 0.339557 | -1.26656 | 0.340683 | 0.983907 | -1.15081 | 0.637566 | 0.842885 |
| -1.16114 | 0.763302 | 0.848093 | -1.10679 | 0.735705 | 0.983907 | 1.113206 | 0.611015 | 0.841398 |
| 1.045131 | 0.040758 | 0.236117 | 1.273163 | 0.256344 | 0.957016 | 1.122673 | 0.920056 | 0.966058 |
| -1.1739 | 0.344085 | 0.545342 | -1.16256 | 0.608379 | 0.983907 | -1.07947 | 0.974286 | 0.992 |
| -1.07326 | 0.150475 | 0.34612 | -1.1959 | 0.828936 | 0.983907 | -1.0252 | 0.665569 | 0.842885 |
| -1.37588 | 0.016999 | 0.139006 | -1.6235 | 0.719473 | 0.983907 | -1.06749 | 0.125843 | 0.618466 |

| JAK2A - Ve | Transformation |
|---|---|
| 1.108771 | LOG |
| -1.00091 | LOG |
| 1.089317 | NO |
| 1.069049 | NO |
| 1.205751 | LOG |
| 1.256055 | LOG |
| 1.087998 | NO |
| -1.14483 | LOG |
| 1.050807 | LOG |
| -1.36377 | LOG |
| -1.42028 | LOG |
| -1.34504 | LOG |
| 1.195149 | LOG |
| 1.2297 | LOG |
| 1.103642 | NO |
| -1.20753 | LOG |
| -1.42445 | LOG |
| -1.31466 | LOG |
| -1.41795 | LOG |
| -1.44003 | LOG |
| 1.001253 | NO |
| -1.0889 | LOG |
| 1.041024 | LOG |
| 1.026724 | LOG |
| -1.34793 | LOG |
| -1.25712 | LOG |
| -1.28116 | LOG |
| -1.04568 | LOG |
| -1.46621 | LOG |
| -1.20352 | NO |
| -1.28266 | NO |
| -1.14771 | LOG |
| -1.40224 | NO |
| 1.044792 | NO |
| -1.58745 | LOG |
| -1.11674 | LOG |
| -1.26888 | LOG |
| 1.024609 | LOG |
| -1.16154 | LOG |
| -1.05363 | LOG |
| 1.535904 | LOG |
| -1.37522 | LOG |
| -1.174 | LOG |
| -1.22773 | LOG |
| -1.0904 | LOG |
| -1.02018 | LOG |
| -1.30944 | LOG |

| | |
|---|---|
| -1.35319 | NO |
| 1.052459 | LOG |
| -1.05489 | LOG |
| 1.001746 | NO |
| 1.100673 | NO |
| -1.12079 | LOG |
| 1.101764 | LOG |
| -1.33391 | LOG |
| -1.06078 | NO |
| -1.29792 | LOG |
| 1.245981 | LOG |
| 1.071521 | LOG |
| 1.090773 | LOG |
| 1.148699 | LOG |
| 1.171612 | LOG |
| -1.48772 | LOG |
| -1.51177 | LOG |
| -1.21166 | LOG |
| 1.148108 | LOG |
| -1.64592 | LOG |
| -1.41939 | LOG |
| -1.47981 | LOG |
| 1.015639 | LOG |
| 1.358369 | NO |
| -1.04234 | LOG |
| 1.045333 | LOG |
| -1.10442 | LOG |
| 1.040905 | LOG |
| -1.0327 | LOG |
| 1.158204 | LOG |
| 1.196189 | LOG |
| 1.093351 | LOG |
| 1.062598 | LOG |
| 1.055542 | LOG |
| 1.003871 | NO |
| -1.12141 | LOG |
| 1.077014 | LOG |
| 1.148413 | LOG |
| 1.103683 | NO |
| 1.069595 | LOG |
| 1.11928 | NO |
| 1.037345 | LOG |
| 1.036373 | LOG |
| 1.038675 | LOG |
| 1.027092 | LOG |
| 1.033715 | LOG |
| 1.038179 | LOG |
| 1.044097 | LOG |
| 1.045131 | LOG |

| | |
|---|---|
| 1.050171 | LOG |
| 1.038071 | LOG |
| 1.083648 | LOG |
| 1.011312 | LOG |
| 1.041844 | LOG |
| -1.01272 | LOG |
| 1.154739 | NO |
| 1.066767 | LOG |
| 1.220726 | LOG |
| 1.026273 | NO |
| -1.01626 | NO |
| 1.036932 | LOG |
| 1.017408 | LOG |
| -1.024 | LOG |
| 1.228557 | LOG |
| -1.00667 | LOG |
| 1.044565 | LOG |
| 1.040898 | LOG |
| 1.071678 | LOG |
| 1.102771 | LOG |
| 1.260044 | LOG |
| -1.01404 | LOG |
| 1.048195 | NO |
| 1.154908 | NO |
| 1.842205 | LOG |
| 1.291467 | LOG |
| -1.3922 | LOG |
| -1.08675 | NO |
| 1.391381 | NO |
| -1.14191 | LOG |
| -1.27433 | LOG |
| -1.35958 | LOG |
| -1.33 | LOG |
| -1.43167 | LOG |
| -1.33671 | LOG |
| -1.20978 | LOG |
| -1.38992 | LOG |
| -1.05086 | NO |
| -1.39033 | LOG |
| -1.55207 | LOG |
| -1.62095 | LOG |
| -1.34791 | LOG |
| -1.58965 | LOG |
| -1.56058 | LOG |
| -1.48861 | LOG |
| -1.34612 | LOG |
| -1.64106 | LOG |
| -1.61386 | LOG |
| -1.31641 | LOG |

| | |
|---|---|
| -1.33344 | LOG |
| 1.083484 | LOG |
| -1.23671 | LOG |
| -1.29971 | LOG |
| -1.41846 | LOG |
| -1.26728 | LOG |
| -1.08488 | LOG |
| -1.42119 | LOG |
| -1.43352 | LOG |
| -1.38425 | LOG |
| -1.22902 | LOG |
| -1.32984 | LOG |
| -1.29932 | LOG |
| -1.22042 | LOG |
| 1.058646 | LOG |
| -1.33896 | LOG |
| -1.02206 | LOG |
| 1.075952 | LOG |
| -1.18723 | LOG |
| 1.011503 | NO |
| -1.0051 | LOG |
| -1.05429 | LOG |
| -1.35156 | LOG |
